# A spatial human thymus cell atlas mapped to a continuous tissue axis

**DOI:** 10.1101/2023.10.25.562925

**Authors:** Nadav Yayon, Veronika R. Kedlian, Lena Boehme, Chenqu Suo, Brianna Wachter, Rebecca T. Beuschel, Oren Amsalem, Krzysztof Polanski, Simon Koplev, Elizabeth Tuck, Emma Dann, Jolien Van Hulle, Shani Perera, Tom Putteman, Alexander V. Predeus, Monika Dabrowska, Laura Richardson, Catherine Tudor, Alexandra Y. Kreins, Justin Engelbert, Emily Stephenson, Vitalii Kleshchevnikov, Fabrizio De Rita, David Crossland, Marita Bosticardo, Francesca Pala, Elena Prigmore, Nana-Jane Chipampe, Martin Prete, Lijiang Fei, Ken To, Roger A. Barker, Xiaoling He, Filip Van Nieuwerburgh, Omer Bayraktar, Minal Patel, Graham E. Davies, Muzlifah A. Haniffa, Virginie Uhlmann, Luigi D. Notarangelo, Ronald N. Germain, Andrea J. Radtke, John C. Marioni, Tom Taghon, Sarah A. Teichmann

## Abstract

T cells develop from circulating precursors, which enter the thymus and migrate throughout specialised sub-compartments to support maturation and selection. This process starts already in early fetal development and is highly active until the involution of the thymus in adolescence. To map the micro-anatomical underpinnings of this process in pre- vs. post-natal states, we undertook a spatially resolved analysis and established a new quantitative morphological framework for the thymus, the Cortico-Medullary Axis. Using this axis in conjunction with the curation of a multimodal single-cell, spatial transcriptomics and high-resolution multiplex imaging atlas, we show that canonical thymocyte trajectories and thymic epithelial cells are highly organised and fully established by post-conception week 12, pinpoint TEC progenitor states, find that TEC subsets and peripheral tissue genes are associated with Hassall’s Corpuscles and uncover divergence in the pace and drivers of medullary entry between CD4 vs. CD8 T cell lineages. These findings are complemented with a holistic toolkit for spatial analysis and annotation, providing a basis for a detailed understanding of T lymphocyte development.

## Introduction

T cells play a central role in adaptive immunity, aiding the development of the humoral immune response, eliminating infected and cancerous cells through T cell-mediated killing, and regulating specific and targeted immune responses while avoiding prolonged inflammation or autoimmunity. This is achieved by distinct T cell subtypes, which include conventional helper, cytotoxic, and regulatory T cells (Tregs) as well as unconventional T cell types, such as yδT cells, Natural Killer T (NKT) cells, Mucosal-associated invariant T (MAIT) cells, and CD8αα intraepithelial lymphocytes (lELs), that exhibit innate-like characteristics in their antigen recognition and effector functions^1,2^. All T lineage cells originate from bone marrow-derived precursors but develop in the thymus, which originates from the bilateral thymic primordia that migrate and attach to the pericardium in post-conception week (pcw) 8 of human embryonic development, giving rise to the two lobes of the organ^3^. Subsequently, the tissue is seeded by lymphoid precursors as well as vascular and mesenchymal cells, and the precursors of thymic epithelial cells (TECs) begin to differentiate into different functional subtypes^3^. Previous reports by us and others have suggested pcw 12-14 as the earliest time point when mature T cells are first released into the periphery and indicated that a clear morphological division between the internal medulla and the surrounding cortex is established around pcw 12-16^3-6^. Although thymopoiesis remains highly active until adolescence, degeneration and involution of the tissue starts already during paediatric development and involves the gradual infiltration with peripheral lymphocytes and adipocytes, which eventually results in a loss of clear demarcation of cortex and medulla and severe reduction in thymic output in the aged thymus^7^.

The cortex marks the region where bone marrow-derived precursor cells commit to the T lymphocyte lineage, first becoming CD4- CD8- double negative (DN), then CD4+ immature single positive (ISP), and later CD4+ CD8+ double positive (DP) T lineage cells that undergo V(D)J recombination of their T cell receptor (TCR) loci. This is followed by the positive selection of DP thymocytes with a functional TCR through interactions with peptide-Major Histocompatibility Complex (MHC) ligands on cortical TECs (cTECs). Intracellular signals arising from these interactions prompt the lineage bifurcation into CD4+ and CD8+ single positive (SP) thymocytes. Upon migration of positive selected cells into the thymic medulla, they undergo a pruning process termed negative selection, where cells that strongly respond upon TCR engagement with self-ligands found on medullary TECs (mTECs) and other antigen presenting cells, such as dendritic cells (DCs), are deleted or converted into suppressive Tregs to enforce tolerance to self^8^. Beyond the coarse division into cortex and medulla, the thymus harbours secondary morphological structures, such as medullary Hassall’s Corpuscles (HCs), which have been suggested to play an important role in tolerance generation^9^, and less clearly defined regions like the cortico-medullary junction (CMJ), which is highly vascularised and thus often referred to as perivascular space (PVS). The thymus is surrounded by a capsule, which exhibits extensions of connective tissue, called trabeculae/septae, that separate individual thymic lobules^10^.

Our understanding of the thymic architecture and key steps of T cell development is the result of decades of research since the discovery of the thymus as an immunological organ in the 1960s^11^ using a range of increasingly sophisticated and higher resolution methods, from flow cytometry to bulk transcriptomics and more recently single cell RNA sequencing (scRNA-seq). Genetically engineered mouse models and *in vitro* systems have helped elucidate the molecular mechanisms that drive differentiation from bone marrow-derived precursors into the diverse array of mature T cell subsets^12^. Finally, histologic analyses have provided spatial information on the organ subregions in which the various developmental and antigen-driven steps of these maturation processes occur.6,1^3-17^

Recent efforts by us and others to comprehensively catalogue the full range of cell types present in the human thymus using single cell RNA-seq6,^14-16^ have indicated an unexpectedly high degree of diversity not just in the T lineage, but also in other hematopoietic cell types and in the stromal compartment^17-19^. To this end, mapping efforts by the Human Cell Atlas (HCA)^20^, Human BioMolecular Atlas Program (HuBMAP)^21^, and others have identified more than 50 cell types in the paediatric human thymus^14-16^ and have been essential for profiling the cellular composition of the human thymus throughout the first two trimesters of prenatal development^3-6^. The results of these studies raise new questions about the fine-grained organisation of hematopoietic and stromal cells within the thymic lobules and the resulting niches and local chemokine microenvironments that are expected to drive the maturation and migration of thymocytes. These key points are difficult to address with conventional low-throughput spatial technologies^22^. However, the advent of robust methods for spatial genomics^23^ together with highly multiplexed RNA/protein imaging approaches^24-28^ provides the necessary level of resolution to investigate these questions in detail, offering a comprehensive and unbiased view of the cellular and molecular processes taking place in the different compartments of the human thymus.

Still, direct inter-sample comparisons, especially for human material, have been impeded by the variability in tissue morphology and sampling methodology. This represents a substantial hurdle for data alignment and integration between collaborating research groups and within larger consortia, and hampers comparisons across independent studies. In addition, the lack of a common coordinate framework (CCF)^29^, similar to the one established for the mouse brain (http://mouse.brain-map.org/static/atlas), has so far limited spatial annotations to discrete structures or morphological compartments. This hinders the exploration of inter-region variance or intrinsic gradients and restricts the ability to model thymocyte maturation in continuous space, especially at the interface between histological structures and cellular neighbourhoods. To overcome these obstacles, we set out to create a continuous CCF for the human thymus. This enabled us to establish consistent structural annotations that can be used as a reference for future thymus studies and to obtain a more holistic portrait of this critical immunological organ throughout early human development. Additionally, it serves as a platform to directly compare and integrate biological observations across modalities, replicates, unpaired samples, and different institutions.

To provide an in-depth cellular and spatial atlas covering pre-natal development to three years of age, which represents the early to peak stage of the organ function prior to the slow degeneration of the thymus, we integrated several types of thymic spatial data with the largest multi-modal single cell annotation reference of the human thymus to date, spanning more than 500K cells. By applying a newly developed computational tool set for reproducible image annotation (TissueTag) and analysis (ImageSpot), we have derived a mathematical model for a continuous scale- and rotation-invariant morphological CCF (OrganAxis) of the thymus lobules, the “cortico-medullary axis” (CMA). We show that this axis captures the major transcriptomic variance in pre- and post-natal tissue sections and across multiple spatial modalities in each sample individually and collectively.

Through this approach, we describe the organisation of the thymus at a resolution beyond the typically annotated morphological compartments. By comparing pre- and post-natal thymi, we show that broad cytokine/chemokine gradients and cellular sub-zonations are established and constant from as early as pcw 12. We identify a relatively homogeneous cellular distribution in the cortex as opposed to a highly structured organisation of the medulla with a gradient composed of different stromal cell types. Furthermore, we find specific marker gene expression profiles associated with HCs that are independent of the general layering of the thymic medulla. Finally, we show that thymocytes developing along the CD4 and CD8 lineages express different chemokine receptors and correspondingly show divergent timing of cortico-medullary migration.

Beyond these and other biological findings of our study, this combination of high content, multi-resolution data and computational analysis involving a substantial set of fetal and paediatric tissues provides a rich, spatially resolved atlas and CCF of the human thymus for future studies. Importantly, since our spatial tool set and modelling framework are based on tissue landmarks exclusively, it can be adapted to any tissue in 2D or 3D, brightfield or fluorescent images, multiplex antibody-based optical imaging and any spatial transcriptomics platform, thereby serving as a universal tool for continuous spatial annotation and analysis.

## Results

### Combining multimodal spatial and single cell data with novel computational tools to generate an integrated thymus atlas

To robustly map and characterise the cells in the thymus, we first expanded the current coverage of dissociated single cell data sets and refined their annotation, focussing on fetal samples from pcw 10 to 21 and post-natal samples from early infancy until 3 years of age (**Figure 1A**). To expand stromal cell representation, we integrated our previous single cell atlas^6^ with two recent studies^17,30^ that had specifically enriched for epithelial, endothelial, myeloid, and fibroblast lineages. Additionally, we added *de novo* generated data from 9 paediatric donors, resulting in a data set totalling over 500K high-quality cells, including TCR sequencing (scTCR-seq) for more than 100K cells as well as 120K cells with paired CITE-seq and scTCR-seq data, and 38K cells from CITE-seq experiments, for which only RNA could be detected (**Figure 1A, Extended Data Figure 1A-E, Supplementary Table 1**).

**Figure 1.**
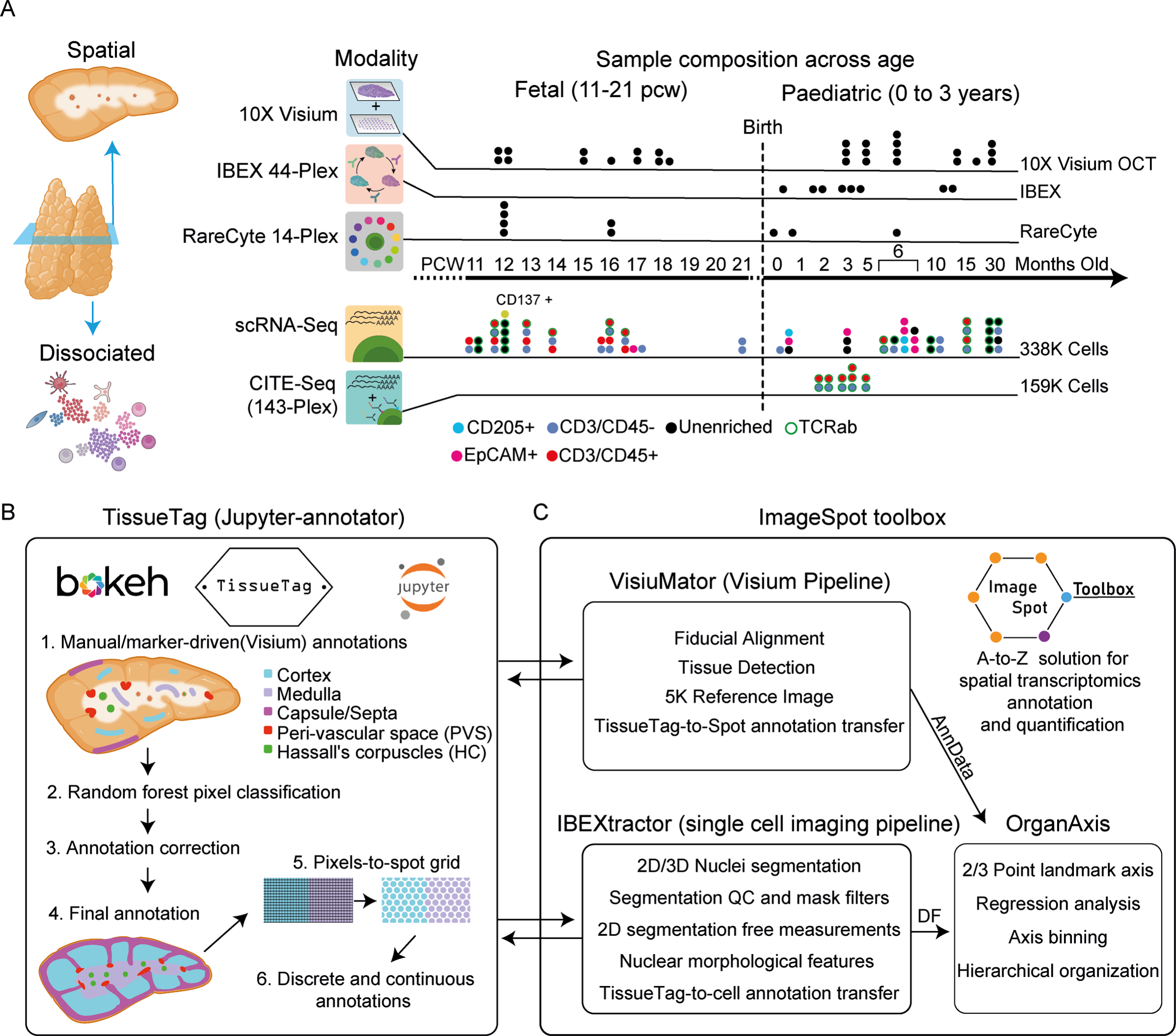
Spatial and single cell atlas data and methodology. **A.** Sample composition for dissociated and spatial datasets that span fetal (pcw 11 to 21) and early paediatric human life (0 to 3 years) from both newly generated data as well as data curated from recent studies. Top: Spatial datasets include IBEX 44-plex protein cyclic imaging, RareCyte single-cycle 14-plex protein imaging and 10x Genomics Visium spatial transcriptomics, along with 3 fetal Visium data sets we had published previously (Suo et al 2022). Bottom: Dissociated datasets are composed of scRNA-seq coupled with TCR-seq forTCRαβ, 143-plex surface protein CITE-seq coupled with TCR-seq for TCRαβ. Each dot represents a sample, stacked dots represent libraries from the same donor. Dot colour indicates enrichment strategy applied to cells prior to library generation. Dot border colour indicates whether TCR-seq was carried out. See Supplementary Table 1–2 for sample origin and metadata; also see Extended Data Figures 1, 2. B. TissueTag tool enables interactive (powered by bokeh), manual and semi-automated image annotation directly in a Jupyter environment allowing for unified annotation and analysis. TissueTag annotations are converted from pixel to spot grid space, where continuous distance measurements are possible. **C.** Components and basic workflow of the ImageSpot toolbox for complete image processing throughout this study. VisiuMator and IBEXtractor pipelines support TissueTag-derived annotations to be transferred to 10x Visium and high-resolution single-cell imaging data, respectively. Finally, OrganAxis allows generation of a landmark CCF for two or more landmarks combined and provides downstream analysis for continuous CCFs.

To link these single cell types with the landscape of the thymus, we implemented three complementary spatial technologies to generate rich data sets matching the two developmental windows that had been sampled in the dissociated single cell data. Specifically, we applied the 10x Genomics Visium spatial transcriptomics platform using tissues from paediatric and fetal thymus samples, and incorporated 3 fetal thymus Visium sections that were published in our previous study^31^, totalling 28 sections (fetal: 12 sections from 7 donors, paediatric: 16 sections from 6 donors)(**Supplementary Table 2**). Additionally, we generated high-resolution, single-cell thin-section protein imaging from post-natal thymi using the IBEX method^32^ with a 44-plex antibody panel curated specifically for TEC and stromal analysis (8 sections from 8 paediatric donors), as well as RareCyte 14-plex broad immunological protein staining from both fetal (6 sections, 2 donors) and paediatric tissue samples (3 sections, 3 donors)(**Figure 1A, Supplementary Table 2**).

To generate consistent image annotations from all spatial data sets, we developed TissueTag (https://github.com/nadavyayon/TissueTag), a Jupyter notebook-based interactive image annotation tool. TissueTag is powered by the Bokeh python library (https://docs.bokeh.org/en/latest/) and provides an annotation solution of tissue regions at subpixel resolution from various image types (brightfield, fluorescence, simulated, etc.) as well as for spatial transcriptomics (**Figure 1B, see Methods**). Annotation of the reference image overcomes resolution constraints (e.g., resulting from the Visium spot size (55 μm) and spacing (100 μm)), thus preserving the spatial sampling resolution across different samples by focusing on absolute scaling. TissueTag creates discrete annotations (“cortex”, “medulla”, etc.) as well as multi-objects (e.g., annotating individual lobules as “lobule 0,1,2…n”). To generate consistent analysis pipelines across samples and technologies based on TissueTag annotations, we developed ImageSpot-toolbox (https://github.com/Teichlab/thymus_spatial_atlas/ImageSpot) (**Figure 1C**), which consists of 2 components: (1) the custom processing pipelines “VisiuMator” for Visium data (**Extended Data Figure 2A-D**) and “IBEXtractor” for IBEX imaging data (**Extended Data Figure 3A-B**). These tools enable pre-processing for Visium and 2D/3D cell segmentations for IBEX data and build on TissueTag annotations to gain spatial insights, as well as (2) “OrganAxis”, a mathematical continuous spatial modelling approach, which is used to derive and analyse a landmark-based CCF.

To study both broad and specific morphological structures of the thymus, we generated image annotations at multiple levels. Major anatomical structures, such as capsule, cortex, and medulla (**Figure 2A),** were annotated as “level 0”, and additional, fine morphological structures, e.g., PVS and HC, were stored in “level 1” (**Figure 2B- D).** Additionally, we recorded discrete lobules with “object_annotator” (**Extended Data Figure 2B, 3A).** Finally, TissueTag also generates “continuous annotations” by calculating the position of each cell/spot according to its relative distance to a histological annotation in (e.g., capsule, cortex and medulla) with adjustable resolution (**Figure 2E, Methods**).

**Figure 2.**
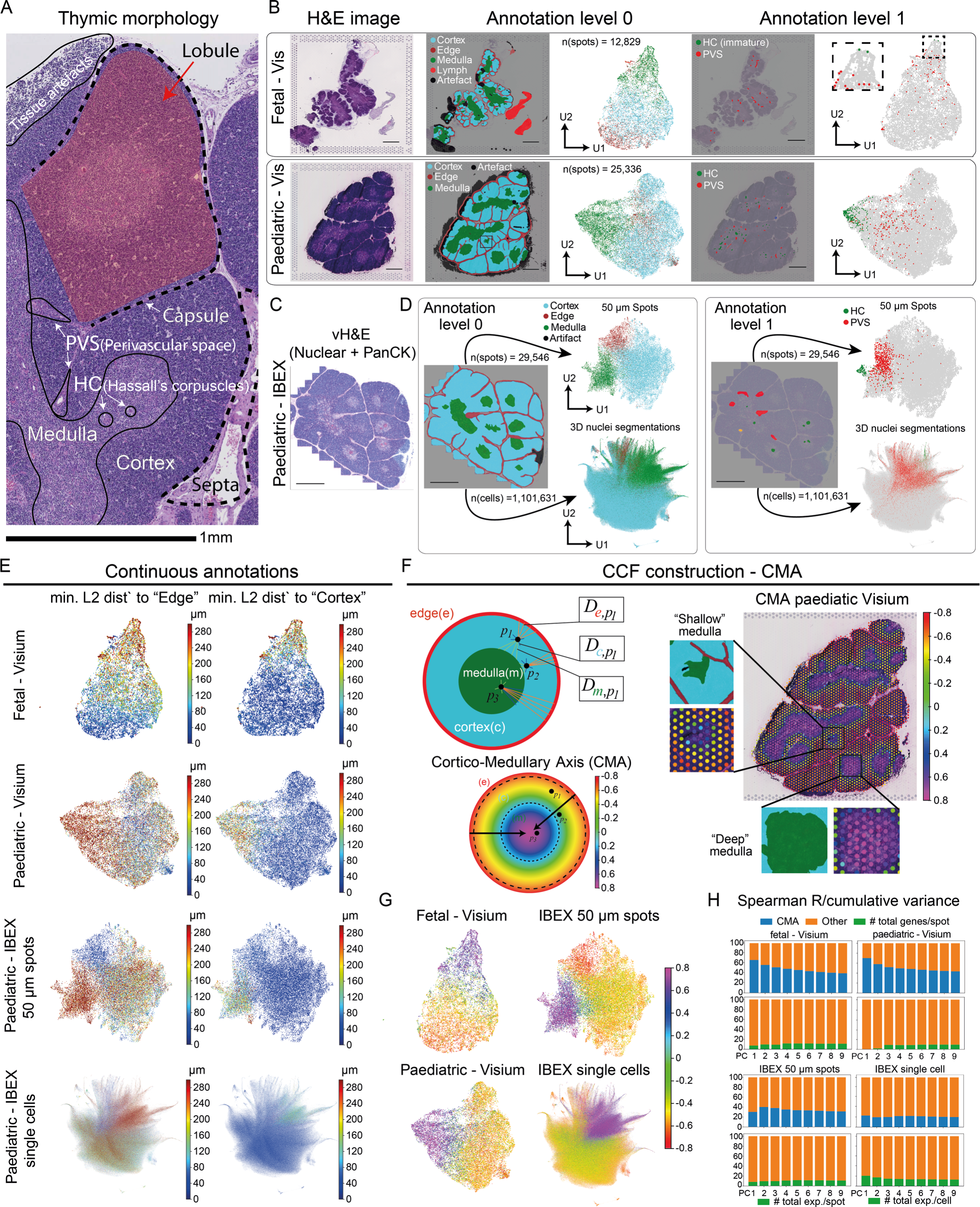
Relative distance-based construction of a continuous cortico-medullar morphological axis from image annotation landmarks. **A.** Representative H&E image of a section from a paediatric thymus showing the major anatomical subcompartments. **B.** Representative H&E images (left), discrete broad level annotations (level 0)(centre), and fine structure manual annotations (level 1)(right) of fetal (top) and paediatric (bottom) Visium samples paired with gene expression-corrected UMAP embeddings with corresponding annotations. Box highlights two Visium spots showing immature Hassall’s Corpuscles in fetal samples. PVS: perivascular space, HC: Hassal’s corpuscle. **C.** IBEX Virtual H&E image, constructed from nuclear Hoechst (blue) and Pan-Cytokeratin (magenta). **D.** Broad level annotations (left) and 44-protein measurement UMAP embedding on two resolution levels - 50 μm spot grid (top) and single nuclei 3D segmentations (bottom). Manual image annotations of PVS and HCs represented on UMAPs from both data types (right). All scale bars in A-D are 1 mm **E.** Continuous annotations of Visium and IBEX datasets plotted on batch-corrected UMAP space coloured by the minimal Euclidean distance to the nearest edge (capsule/septa, left) or to the cortex (right). **F**. Schematic illustration of the construction of the Cortico-Medullary Axis (CMA) CCF (left). Discrete annotations of edge (e), cortex (c) and medulla (m) (top). Function D is visually highlighted in (E). The combination of Euclidean distances between each high-resolution spot to the nearest anatomical structure as derived from the tissue annotations serves as a basis for generation of a continuous axis from the edge of the cortex to the centre of the medulla (bottom). Projection of the CMA to Visium spot space highlighting a differential representation of discrete (level 0) annotations from the CMA that accounts for the relative size of a structure (e.g., the medulla) to be inferred as 3D depth (right). **G.** UMAP representation of integrated data sets coloured by CMA values to highlight the consistency across age and spatial modality. **H.** Sum of the Spearman correlation coefficient R of individual PCA components with the CMA and corrected for explained PCA variance of individual PCs, highlighting the similarity of captured transcriptomic variance between fetal and paediatric data sets as opposed to the technical random factor (n_genes_by_spots).

To gain single cell resolution and overcome challenges with single cell segmentation in the dense thymic tissue, we extracted two levels of information from IBEX imaging data sets: (1) 3D nuclear segmentations from thin section (6-8 μm) Z stacks and (2) a 50 μm hexagonal grid mask to unbiasedly quantify the protein expression levels of structural protein markers and structural features that are not captured well using nuclear segmentation (e.g., large, branched cells such as thymic epithelial cells or vessels). This grid-based analysis also enables a more direct comparison to Visium data, which is restricted to a grid of hexagonal spots of 55 μm diameter (**Figure 2D**).

### Establishment of a CCF for the human thymus using OrganAxis

A major consideration for spatial tissue atlas construction is the establishment of a common reference framework that permits the proper integration of spatial data from diverse samples and experimental modalities, irrespective of biological variability between different donors or technical discrepancies in image acquisition. We aimed to construct such a continuous CCF for the thymus to map and compare the microenvironments that individual thymocytes encounter throughout their maturation and to ask whether the thymus exhibits internal organisation beyond broad morphological structures.

Due to the complex multi-lobular nature of the thymus, it is not possible to simply draw a line connecting thymus landmarks from the capsule across the CMJ to the middle of the medulla, or to use discrete mapping or assignment to “capsular”, “cortical”, or “medullary” locations without losing potentially important information about relative positioning within each micro-compartment. Moreover, using 2D sections limits the ability to identify core features, such as the centre of a medullary region, due to its irregular shape and depth. Finally, it is difficult to annotate discrete lobules as they vary dramatically in size and shape, and the sectioning plane will determine where one lobule ends and the next one starts, irrespective of the full 3D connectivity of these structures.

Consequently, we developed a spatial model that is “boundary-centred” and is computed using a sigmoid weight function that increases resolution close to the junction between two structures. In practice, the model uses TissueTag-derived continuous annotations to the three major histological structures in the thymus (capsule, cortex and medulla)(**Figure 2E**) and takes a weighted combination of two different normalised distances (capsule-to-cortex and cortex-to-medulla). This metric, which we termed the Cortico-Medullary Axis (CMA), uniquely defines the relative location of any cell or spot within the thymus macrostructure (**Figure 2F, Extended Data Figure 4, Methods**). The model assumptions are that the relative position within a structure will depend (sigmoidally) on the immediate proximity to a neighbouring structure/landmark, e.g., medullary regions that are relatively far from capsular/cortical areas would be ranked as “deeper” than medullary areas that are closer to capsular/cortical regions and will thus be ranked as “shallow” (**Figure 2F**). Through the assignment of relative positions, the CMA also resolves cases where Visium spots are located at the interface of two discrete structures and cannot unambiguously be assigned to one or the other. Since the CMA is calculated using relative distances to morphological landmarks, this permits a direct comparison of different samples with distinct spatial modalities, section sizes, donor ages or resolutions (**Figure 2G, Supplementary Figures 1-3**).

To estimate how well the CMA captures biological variance within samples, we performed Principal Component Analysis (PCA) on the batch corrected Visium and IBEX data sets and calculated the Spearman correlation of the first 9 principal components to the CMA as well as to the total number of counts (the most significant technical factor in Visium samples) or total marker expression levels in IBEX. To get a relative estimate of variance, we further normalised the regression coefficient by the cumulative explained variance. We found that the CMA was highly correlated to the first principal components and, more importantly, explained similar degrees of variance in both fetal and paediatric Visium data sets indicating a consistency in representation of transcriptomic diversity across both developmental stages regardless of their clear morphological difference. For IBEX data sets, the association with the CMA was higher for the 50 μm grid data compared to single nuclei segmentations, indicating that the former captures a higher degree of spatial variability. This suggests that single nuclei segmentations lack local environment information that is represented by the location along the axis (**Figure 2H**).

### The T cell developmental trajectory is largely conserved between fetal and paediatric thymus

Substantial differences in the architecture of the thymus are observed throughout development; for instance, the fetal thymus is much smaller in size and has segregated bud-like cortical lobules that are interconnected by the medullary space, whereas the paediatric thymus has much larger cortical regions that are fused together or separated by a thin capsule (**Figure 2B**). To obtain matching spatial annotations for fetal and paediatric tissue sections despite these macromorphological variations, we mapped all spots in the Visium data to the CMA to derive continuous spatial localisation values. Furthermore, for the axis to serve as a common reference and to aid with visualisation, comparison and enrichment score analysis within larger compartments, we defined subregions along the CMA corresponding to the major anatomical structures (capsule, sub-capsule, cortical CMJ, medullary CMJ) and then further subdivided the cortex and medulla into layers using specific axis values (cortical levels I-III, medullary levels I-II)(**Figure 3A, see Methods**).

**Figure 3.**
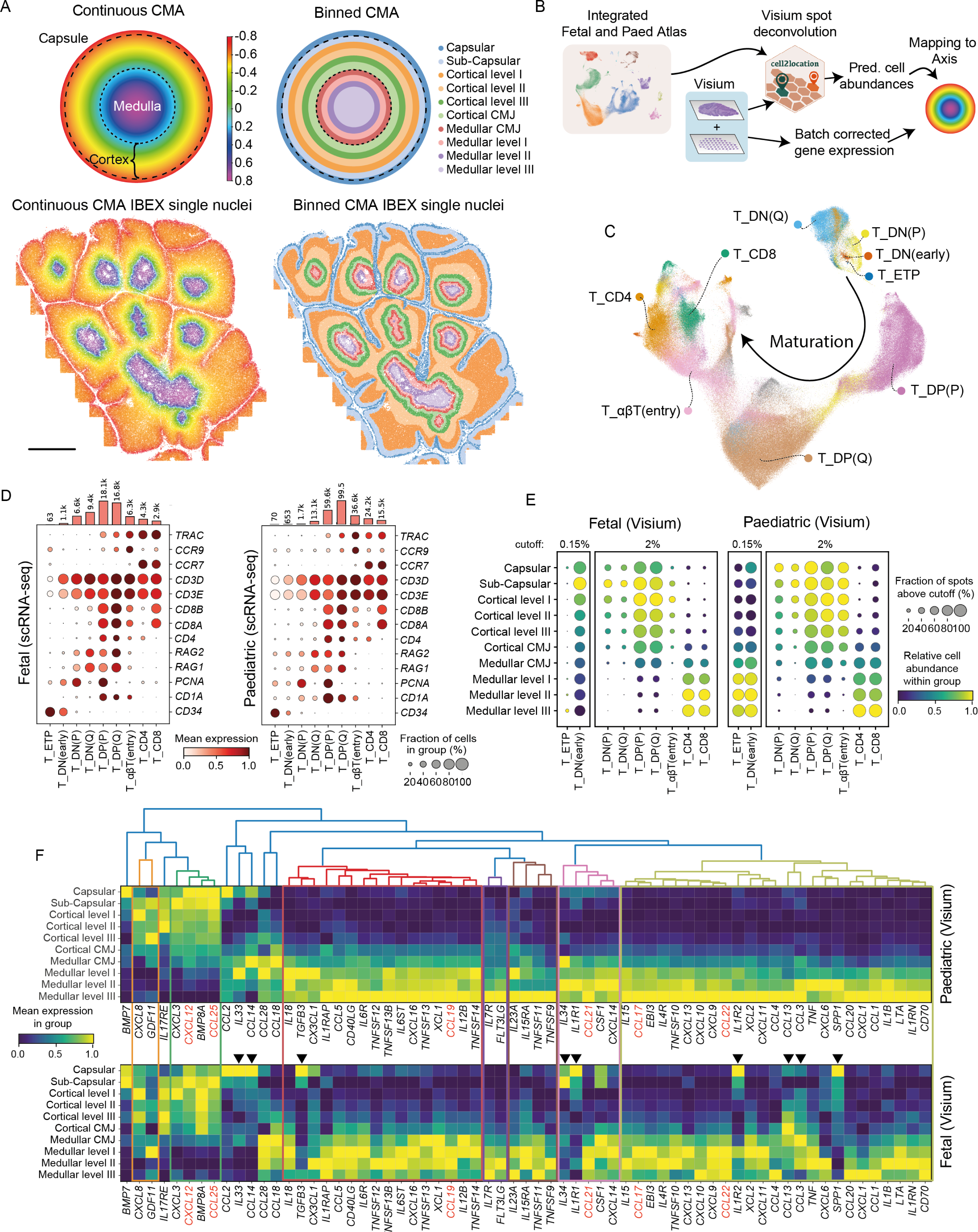
Mapping of fetal and paediatric thymocytes to the CMA reveals largely conserved trajectories for cells of the αβ T lineage. **A.** Illustration of the continuous (left) and binned (right) CMA as a schematic and overlaid on a representative IBEX section from paediatric thymus. **B.** Schematic illustration of the analysis workflow. scRNA-seq references are used for deconvolution of Visium spots via cell2location. The CMA is calculated for each cell/spot and predicted cell abundances, gene and protein expression per spot are interpreted and compared by fine-grained binning of the continuous CMA. **C.** UMAP embedding of integrated fetal and paediatric scRNA-seq data from T lineage cells and annotation of developmental stages. **D.** Expression of marker genes across the main differentiation stages of the T lineage in fetal and paediatric scRNA-seq data. Total number of captured cells per stage is indicated in the associated bar graph. **E.** CMA mapping of the main differentiation stages of the T lineage for fetal and paediatric Visium sections. ETP and DN_early stages were plotted separately with adjusted cutoff to aid visualisation of these rare subsets. **F.** Chemokine and cytokine transcript gradients derived from Visium spot data. Arrows highlight cytokines/chemokines with strong differences in their expression pattern between fetal and paediatric thymus. Cytokines are clustered by spatial pattern similarity across the CMA for the paediatric thymus. Selected cytokines/chemokines that are critical for thymocyte migration and maturation are highlighted in red.

To compare the same cell types and states from both fetal and paediatric T cell maturation trajectories and resident cell distributions, we integrated fetal and paediatric single cell data sets and generated common cell annotations based on marker genes identified in previous atlases (**Figure 3B-D, Extended Data Figure 5A- C**)^6,31^. We then mapped the local abundance of annotated cell subsets in Visium spatial transcriptomics data sets using cell type deconvolution with cell2location^33^. We found a remarkably conserved distribution of canonical αβ T lineage thymocytes along the CMA across development (**Figure 3E, Extended Data Figure 5D**), which complements our previous observation that the cellular outputs of the main αβ T cell lineage differentiation stages are generally constant from pcw 11 up until 3 years of age^3-6^. Relative cell abundances in both fetal and paediatric thymus were also consistent with biological events, with cell numbers increasing due to proliferation in the DP stage and decreasing following selection events at the αβT(entry) stage (**Figure 3E**).

The thymic entry of lymphoid progenitors has previously been described to occur in the medulla or at the CMJ^34-36^ and our Visium-derived CMA prediction did indeed pinpoint the Early T cell Progenitor (ETP) stage to the medulla in both fetal and paediatric tissue (**Figure 3E**). The subsequent DN(early) differentiation stage was predicted to predominantly localise to the subcapsular region of the fetal thymus, whereas in the paediatric tissue this thymocyte subset was still mostly medullary, although both stages showed substantial migratory activity as reflected by the wide distribution across all thymic layers (**Figure 3E**). Since the cortex notably increases in size and cellular density throughout development due to the substantial expansion of the cortical DP population, we speculate that the apparent differences in the DN(early) localisation are a consequence of the relative reduction in the proportion of these cells in the cortex due to the overwhelming number of cortical DP thymocyte in the paediatric thymus. We found that the subsequent maturation stages followed the conventional circular migration path^37,38^, highlighting the (sub-)capsular localisation of DN and proliferating DP(P) cells, followed by a more distributed cortical pattern in the more mature quiescent DP(Q) thymocytes that are in the process of TCR α-chain rearrangement. Cells undergoing positive selection αβT(entry) were still cortically located, but enrichment in the medulla was observed upon commitment to the CD4 or CD8 T lineage (**Figure 3E**). Hence, we show that by applying the CMA to spatial transcriptomics data and integrating this with a scRNA-seq reference, we are able to track the entire spatial trajectory of T lineage differentiation and demonstrate that it is already broadly established by pcw 12-18.

To elucidate possible driving forces governing this migration process, we explored cytokine/chemokine expression in our Visium data set. To this end, we extracted the distribution of the major cytokines/chemokines that were expressed in the thymus from the CellphoneDB database^39,40^ and calculated their relative distribution across the binned CMA. We then performed hierarchical clustering of these distribution profiles, detecting six co-expressed cytokine groups in the paediatric Visium data. Cytokine spatial organisation patterns were generally conserved between fetal and paediatric thymus samples, as was the expression of the major chemokines known to direct thymocyte migration, such as the cortical chemokines *CCL25* and *CXCL12* that serve as ligands for *CCR9* and *CXCR4* respectively, and the medullary chemokines *CCL19* and *CCL21* that act as ligands for *CCR7^22^* (**Figure 3F**). In contrast, several of the cytokines with high expression in the fetal capsule, e.g., *IL34*, and *SPP1*, were more enriched at the CMJ or in the medulla of paediatric samples, suggesting a differential role or source of these factors in pre-/post-natal thymus. Interestingly, these cytokines were predominantly expressed by endothelial cells, fibroblasts (*CCL14*, *IL33, IL34*, *TGFP3*), macrophages (*SPP1*, *CCL3* and *CCL13),* and paediatric TECs (*IL1R1* and *IL1R2*)(**Extended Data Figure 6A**). Importantly, the cytokine mapping was solely based on transcript abundance and does not take into account translation, secretion, and other means of controlling chemokine distribution^41^.

In summary, despite the pronounced differences in the overall structure of the fetal thymus, including rapid growth and the formation of vasculature, we have documented the early organisation of cytokine and chemokine gradients in the fetal thymus development.

### Distributions of resident stromal and immune cells vary between fetal and paediatric thymus

In addition to T cells, we annotated and spatially positioned resident stromal and immune cells that support and enable T cell maturation, including TECs, fibroblasts, endothelial cells, mural cells, B cells, and myeloid cells (**Extended Data Figure 7, 8**).

Thymic fibroblasts can broadly be grouped into capsular fibroblasts, which are a source of growth and signalling factors for TECs and T cells, and medullary fibroblasts, which form an interconnected network of conduits in the medulla that sequester cytokines such as CCL21^42^. Some of the medullary fibroblasts are also capable of presenting antigens and thus contribute to negative selection^43^. In our data set we annotated two sets of capsular fibroblasts, which we refer to as Interlobular (“InterloFb”) and Perilobular (“PeriloFb”) according to our previous atlas^6^(**Extended Data Figure 7A, B**). As expected, Visium-based spatial mapping indicated a clear capsular and subcapsular localisation for these two fibroblasts subtypes in both fetal and paediatric thymi (**Extended Data Figure 7C**). In line with the description of medullary fibroblasts subsets in recent studies^16,44^, we identified three subtypes of putative medullary fibroblasts (“medFb”), which displayed expression of the medullary cytokine *CCL19* as well as cytokine *IL33* and TNF receptor ligand *TNFSF10* (**Extended Data Figure 7B**). In the paediatric thymus, all medFB subsets were indeed predicted to reside in the medulla. In contrast, in the fetal tissue only medFB-MHCII^hi^ were mapped to this region, while the other two medFB subsets showed cortical enrichment, potentially indicating ongoing migration or maturation of certain fibroblast types in the fetal thymus (**Extended Data Figure 7C**). Finally, we identified a few small subsets of fetal-specific fibroblasts (“fetFB”) that could not be detected in the paediatric thymus (**Extended Data Figure 7A-B**).

The vascular compartment was represented in our data set by arterial (“Art”, *HEY1*+), venous (“Ven”, *PLVAP*+), and capillary cells (“Cap”, *RGCC*+) as well as mural cells, including pericytes (*RGS5+)* and smooth muscle cells (“SMC”, *ACTA2*+)(**Extended Data Figure 7D-I**). Most vascular cells, including arterial/capillary endothelial cells, elastin-expressing venous endothelial cells (“EC-Ven-ELN”), and most pericyte subtypes were predicted to be situated in the capsular/subcapsular regions of the fetal thymus (**Extended Data Figure 7F, I**). In contrast, only capillaries and lymphatic vessels showed cortical mapping in the paediatric thymus, while arterial cells and most pericytes were enriched at the CMJ or in the medulla. In both cases, venous cells were mostly detected in the medulla and at the CMJ, together with smooth muscle cells and *CCL19*-expressing pericytes (**Extended Data Figure 7F, I**).

Consistent with these predicted differences in the distribution of fibroblasts, endothelial cells and pericytes across development, RareCyte imaging confirmed that in the fetal thymus most fibroblasts (VIM+) and blood vessels (CD31+) are situated in the capsular and septal regions, whereas paediatric samples showed the largest vessel formations in the superficial medulla and CMJ (**Extended Data Figure 6B-C**).

In the myeloid compartment we distinguished subsets of conventional DCs (*CLEC9A+* cDC1, *CD1C*+ cDC2), including some *CCR7+ LAMP3*+ activated forms, and plasmacytoid DCs (pDCs, *CLEC4C+*)^3-6^. Proliferating subsets of cDCs and pDCs could be detected and mainly originated from fetal thymus (**Extended Data Fig 8A, B**). We also detected three subtypes of macrophages: fetal-specific *LYVE1+* macrophages, which have previously been described in other tissues^45,46^, a *SPIC1+* thymic subtype observed in mouse^47^ and a subtype of *APOC2+* macrophages. Finally, we noted subsets of developing monocytes and neutrophils mostly derived from the fetal thymus as described in Suo et al.^31^ (**Extended Data Fig 8D, E**). In accordance with their involvement in antigen presentation during negative selection, most DC subsets were predicted to reside in the medulla of the fetal and paediatric thymus, with the exception of DC1, which also showed some cortical localisation (**Extended Data Fig 8C**). *SPIC1+* and fetal *LYVE1+* macrophages were predicted to reside in the cortex, while *APOC2+* macrophages were mostly medullary in both fetal and paediatric thymus (**Extended Data Fig 8C**). Most of the monocyte and neutrophil subsets were enriched in capsular and/or medullary regions (**Extended Data Fig 8F**) pointing to their possible location within blood vessels in those areas. We also noted a range of developing *VPREB1*+ B cells and proliferating B cells in the fetal thymus^31^ as well as *TCLA1+* naive, memory and *XBP1+* plasma B cells (**Extended Data Fig 8G**). B cell subsets were mostly predicted to be located in the medulla, showing the strongest enrichment in deep medullary layers, with the exception of fetal pro-B cells, which were predominantly mapped to the cortex and capsule (**Extended Data Fig 8I**).

### Spatial mapping of thymic epithelial cells reveals the tissue niches of mcTEC progenitors

Different subsets of TECs play a key role in T cell differentiation and selection, with cTECs supporting the development of DP thymocytes with a diverse repertoire of functional TCRs (positive selection) and mTECs promoting the deletion of autoreactive T cells to promote self-tolerance (negative selection)^48^. Mouse studies have traditionally used the expression of MHCII genes to distinguish immature (MHCII^lo^) and mature (MHCII^hi^) subsets in both cTECs and mTECs^48^. While most mature mTECs express the transcription factor Autoimmune Regulator (AIRE), which allows the transcription of peripheral tissue antigens (PTAs) in random combinations to support negative selection^49-51^, there is also a group of mTECs that downregulate AIRE and express transcription factors and gene modules associated with a specific tissue. As a consequence, these mTECs resemble cell types from corresponding tissues, such as keratinocytes, microfold cells, enterocytes, and ciliated cells and were therefore termed “mimetic cells” 18,^52^. In postnatal development both mTECs and cTECs are thought to arise from a bipotent precursor^53-55^ that resides near the CMJ and hence these precursors are sometimes referred to as “junctional TEC”^56-5^8. While these major groups of cTECs and mTECs have been also described in the human thymus^6,16,17,30,59^, their spatial distribution and change over development are not well-defined. Moreover, the TEC progenitor state in human thymus and its niche had remained unclear and only very recent reports have provided some initial insights into this question^59,60^.

To profile human fetal and paediatric TECs, we carried out enrichment on our scRNA-seq samples by sorting for the mTEC marker EPCAM or the cTEC marker CD205 (encoded by the *LY75* gene) and integrated previously published data sets that had similarly undergone stromal enrichment^6,17,30,31^. This allowed us to identify three subsets of cTECs, multiple mTEC subtypes, and putative progenitor cells (“mcTECs”) as suggested in earlier studies^6,17,30,31^ (**Figure 4A-C**). To infer the spatial localisation of these cells in fetal and paediatric thymus and compare them, we leveraged our Visium data as described before for the T lineage (**Figure 4D**). Furthermore, we used an antibody panel tailored towards TEC detection for IBEX multiplex imaging on paediatric thymus sections (**Extended Data Figure 9A**). To obtain unbiased TEC annotations for IBEX nuclei that were consistent with those in the scRNA-seq data, we performed integration of segmented IBEX nuclei and the paediatric thymus single-cell dataset on the shared gene/protein space using a batch-balanced k-nearest neighbours (KNN) approach (**Figure 4E**). We then applied majority neighbour voting to annotate the IBEX nuclei, which resulted in robust identification of the majority of TEC subsets, although certain specialised mTEC subtypes were difficult to distinguish due to the lack of specific markers in the IBEX antibody panel (**Extended Data Figure 9B, C**, see Methods). Of note, due to international sample access and research regulatory constraints, IBEX imaging of fetal tissue was not possible.

**Figure 4.**
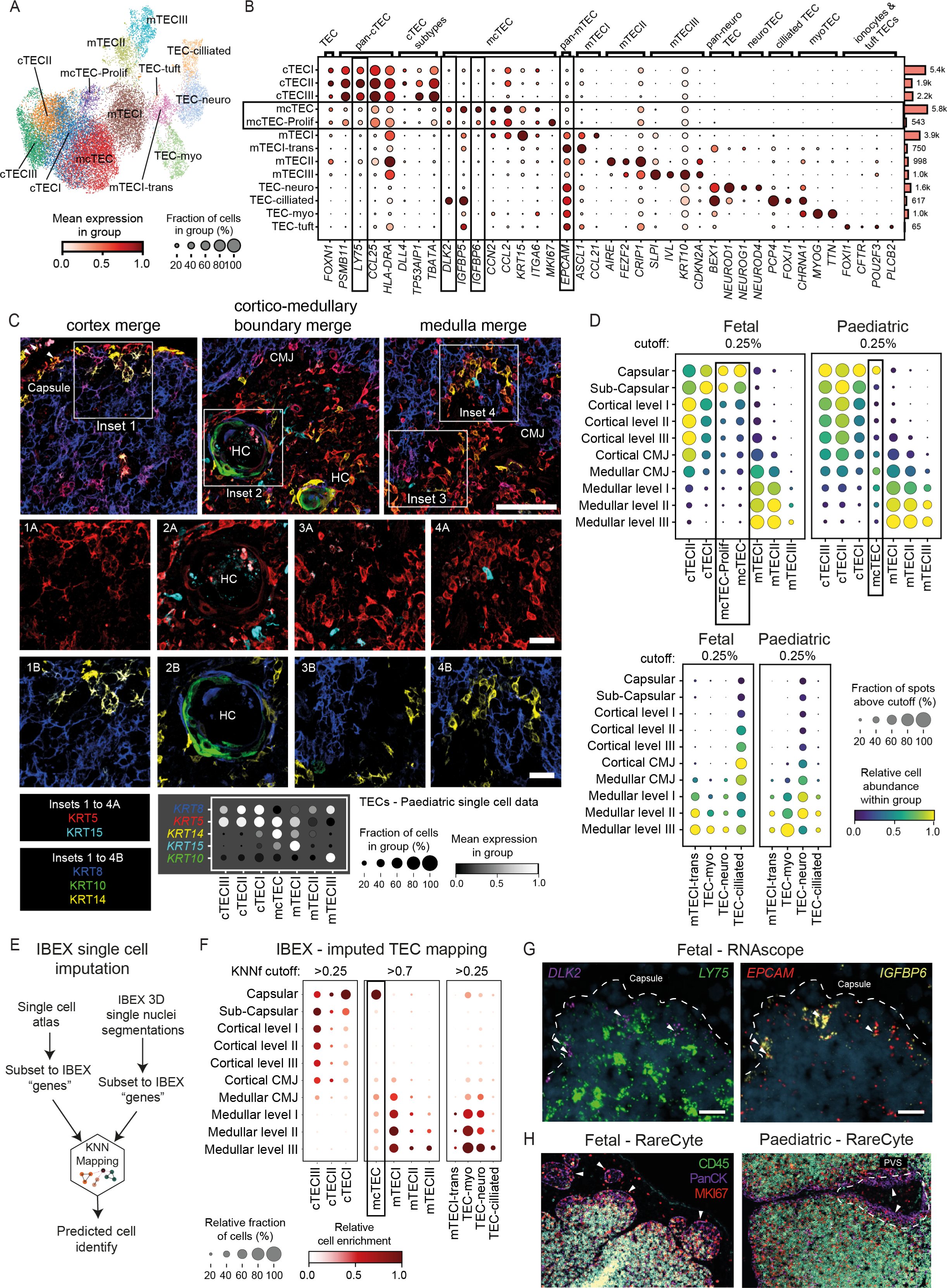
CMA mapping shows broad conservation in the distribution of the TECs in fetal and paediatric thymi. **A.** UMAP embedding of integrated fetal and paediatric scRNA-seq data for thymic epithelial cells (TECs) with cell type annotations. **B**. Dot plot visualisation of RNA expression levels of TEC lineage marker genes. Boxes highlight the mcTEC and mcTEC-proliferating cells and corresponding marker genes used for validation in G. **C.** IBEX confocal images from 7 day old male thymus (Sample_07) showing expression of five different keratins. Scale bar: 100 μm for large insets and 25 μm for small insets. Insets are approximate. Images are representative of 8 similar datasets. Dot plot (bottom) shows transcript levels of the corresponding keratin genes in the main TEC subtypes according to the paediatric scRNA-seq data set. **D.** Relative cell distribution and enrichment in CMA bins for TECs based on Visium spot deconvolution. Boxes highlight mcTECs. Note that proliferating mcTECs were only found in fetal thymus and cTECIII was exclusively detected in paediatric data. Cutoff levels indicate the threshold for the minimum abundance of a cell type in a Visium spot for the spot to be included. **E.** Schematic overview of the workflow of KNN mapping and gene imputation of IBEX single cell segmentations based on matching IBEX cells to the scRNA-seq reference. **F.** Spatial distribution patterns of TEC subsets in the IBEX data after inferring the annotation of segmented cells from the scRNA-seq reference. Dot size represents the relative abundance and colour depicts the local enrichments in the CMA bin. KNNf indicates the cutoff for the fraction of KNNs that corresponded to the eventually assigned majority cell type annotation and indicates mapping quality (see Methods). **G.** RNAscope probe-based staining of the transcripts of mcTEC-specific markers *DLK2* (purple) and *IGFBP6* (yellow) as well as mTEC marker *EPCAM* (red) and cTEC marker *LY75* (green). White arrows indicate the co-expression of all markers in the (sub-)capsular zones in the fetal thymus. Scale bar: 20 μm. **H.** RareCyte protein staining of fetal and paediatric thymus sections with MKI67 (red), Pan-Cytokeratin (PanCK, purple) and CD45 (green) antibodies. Arrows in the fetal image highlight a subcapsular niche in the fetal thymus with proliferating (MKI67+) non-lymphoid (CD45-) keratinized cells. Arrows in the paediatric image highlight keratinized cells in a lymphocyte-free region in the perivascular space (PVS), which do not show extensive proliferation. Scale bars: 200 μm.

Our scRNA-seq data revealed limited heterogeneity in the cTEC compartment with three major subtypes, cTECI-III (**Figure 4A, B**). These differed by expression of MHCII genes, *DLL4, TP53AIP1*, and *TBATA*, a gene associated with TEC proliferation^61^. *DLL4*^lo^ cTECIII cells could only be detected in the paediatric data and were absent in the fetal thymus, which agrees with mouse data showing a decrease in *DLL4* expression in cTECs of the post-natal thymus^62^. Visium mapping of cTEC subtypes predicted that in both fetal and paediatric samples cTECI was predominantly located in the capsular and subcapsular layers, whereas cTECII spanned from the capsule across all the cortical layers up to the CMJ (**Figure 4D**). Analysis of IBEX data confirmed this enrichment pattern for the paediatric thymus, and both IBEX and Visium data agreed on a broadly cortical and capsular distribution of the paediatric cTECIII population (**Figure 4D, F**). Notably, our scRNA-seq data showed that *KRT5*, which (together with KRT14) has often been used as a marker of mTECs, was also strongly transcribed in cTECs and KRT5 protein was clearly detected in both medulla and cortex (**Figure 4C**). This seemingly contradictory *KRT5* transcriptional profile was mirrored in cTECs in a recent human study, suggesting that expression of this gene/protein is not restricted to mTECs^59^.

In the mTEC compartment we could distinguish immature mTECI (*CCL21*+ *MHCII*^lo^ *AIRE*-) and mature *AIRE*-expressing mTECII, which were *MHCII*^hi^ (**Figure 4A, B**). In addition, we identified a set of *AIRE*-negative mature mTEC populations that resemble mimetic cells in the mouse and were described earlier in human^3-6^. These included mTECIII keratinocytes (*IVL*+), myoTEC (*MYOG*+), neuroTEC (*NEUROD1*+), ciliated TECs (*FOXJ1*+), and a TEC subset resembling ionocytes and tuft cells (*FOXI*+) (**Figure 4A, B**). mTECIII were marked by strong expression of *KRT10*, and IBEX imaging clearly placed KRT10+ cells at the HCs in accordance with the known association of mTECIII with these structures^63^ (**Figure 4C**). mTECIII were mostly absent in the fetal thymus except in a handful of small HCs in the oldest fetal sample (pcw 18), consistent with their earliest detection in previous reports^64^ (**Figure 4A, Extended Data Figure 10A**). Both Visium and IBEX mapping of mTEC subsets showed them to be organised in a gradient-like pattern with mTECIII almost exclusively located in the deep medulla and mTECI and mTECII more broadly distributed across all medullary layers (**Figure 4D, F**). Visium analysis predicted this pattern to be conserved between fetal and paediatric thymus. Similarly, a high degree of developmental conservation in their thymic distribution was observed for fetal and paediatric mTECI-trans and myoTECs, which were predominantly detected in the lower medullary layers. In contrast, ciliated TECs showed a notable discrepancy between the two developmental windows, with strong enrichment at the fetal CMJ but clear medullary enrichment in paediatric samples (**Figure 4D**).

In accordance with their putative progenitor nature, mcTECs expressed intermediate levels of cTEC and mTEC markers as well as *KRT15* and *ITGA6* (encoding CD49f), which have been suggested as stem cell markers in previous studies^30,54,65^ (**Figure 4B**). They were further characterised by the transcription of *DLK2*, as well as transcription of insulin-like growth factor binding proteins *IGFBP5*, *IGFBP6*, connective tissue growth factor *CCN2* and the cytokine *CCL2*. Finally, we noted strong expression of *KRT5* and *KRT14*, which aligns with the very recent description of this progenitor population as “polyKRT cells”^59^ (**Figure 4C**).

Our Visium and IBEX mapping of paediatric mcTECs predicted them to be predominantly found in the thymic capsule and around the medullary CMJ (**Figure 4D, F**). Inclusion of secondary structural annotations in the spatial mapping revealed that mcTECs in the vicinity of the CMJ were in fact mostly localised in the PVS region (**Extended Data Figure 10B**). This aligns well with the recent report from Ragazzini et al.^59^, which suggested the subcapsular region and PVS as the TEC progenitor niche. Based on the keratin expression profile of mcTECs in the scRNA-seq data (*KRT8*- *KRT5*+ *KRT14*+ *KRT15*+), we were also able to further validate putative mcTEC niches in the capsule and at the CMJ of the paediatric thymus (**Figure 4C**, inset 1 and inset 4), whereby KRT15 expression was not seen in the capsular TECs, suggesting potential differences between the capsular and CMJ mcTEC populations. Interestingly, Visium mapping of fetal mcTECs predicted them to be strongly enriched in the capsule or subcapsular regions and not at the CMJ, which was independent of their proliferative activity (**Figure 4D**). To further test this prediction, we carried out single molecule FISH (RNAscope), which confirmed a clear capsular localisation of cells expressing the mcTEC markers *DLK2* and *IGFBP6* in the fetal thymus (**Figure 4G, Extended Data Figure 10C**). Finally, multiplexed RareCyte protein imaging indicated the presence of proliferating epithelial cells (MKI67+ CD45- PanCK+) in very distinctive, early fetal (pcw 12) subcapsular zones, which may further indicate the location of the putative proliferating mcTEC niche. No similar cells were detected in the paediatric capsule, but some PVS regions in CMJ and medulla were found to contain non-proliferating epithelial cells (MKI67- CD45- PanCK+) cells (**Figure 4H**).

Overall, our data are consistent with a recent report identifying the PVS and subcapsular region as niches of TEC progenitor cells in the paediatric thymus^59^. In addition, we report exclusive (sub-)capsular localisation and increased proliferative activity of putative TEC progenitors, mcTECs, in the fetal thymus. This suggests a change in the mcTEC localisation and proliferative potential across early human development.

Beyond clarifying the spatial niches of various TEC subtypes, the reproducibility of our findings between two completely different data types - RNA sequencing-based Visium spot data and single cell resolution IBEX multiplex protein imaging - highlighted the utility of the CMA in comparing and quantifying cell/marker/gene distributions across spatial analysis platforms. Moreover, the overarching similarities in the distribution of developing T cells and supporting stroma, myeloid, and B cells across fetal and paediatric thymus, despite considerable morphological and transcriptional differences of the tissue, further emphasised the usefulness of the CMA for comparison of different biological samples. Together, this has revealed (sub-)capsular regions as a “cell type hub” of the fetal thymus, harbouring putative TEC progenitor cells, most fibroblasts and vascular cell types as well as a range of DN and DP thymocytes.

### Specific mTEC subsets show increased association with Hassall’s Corpuscles

HCs are medullary structures formed from keratin layers composed of post-AIRE mTECIII^63^ that arise in mid-gestation (**Figure 5A**). While the specific functions of HCs are unknown, previous studies have proposed that secretion of TSLP from corpuscular mTECs promotes CD80/CD86 expression on DCs, which in turn supports the development of Tregs and may also play a role in negative selection^9,66^. Expression of auto-antigens has been reported in HCs, with connections to various auto-immune diseases, such as rheumatoid arthritis, diabetes, myasthenia gravis and Hashimoto’s thyroiditis^67-70^. In addition, several studies have suggested association of HCs with cell types such as B cells, pDCs, and neutrophils in addition to Treg cells^71-73^. These findings point towards an involvement of HCs in the regulation of central tolerance and indicate the importance of a comprehensive, unbiased profiling of these structures to better understand their connection with other thymic cell types and their physiological function in the thymic context.

**Figure 5.**
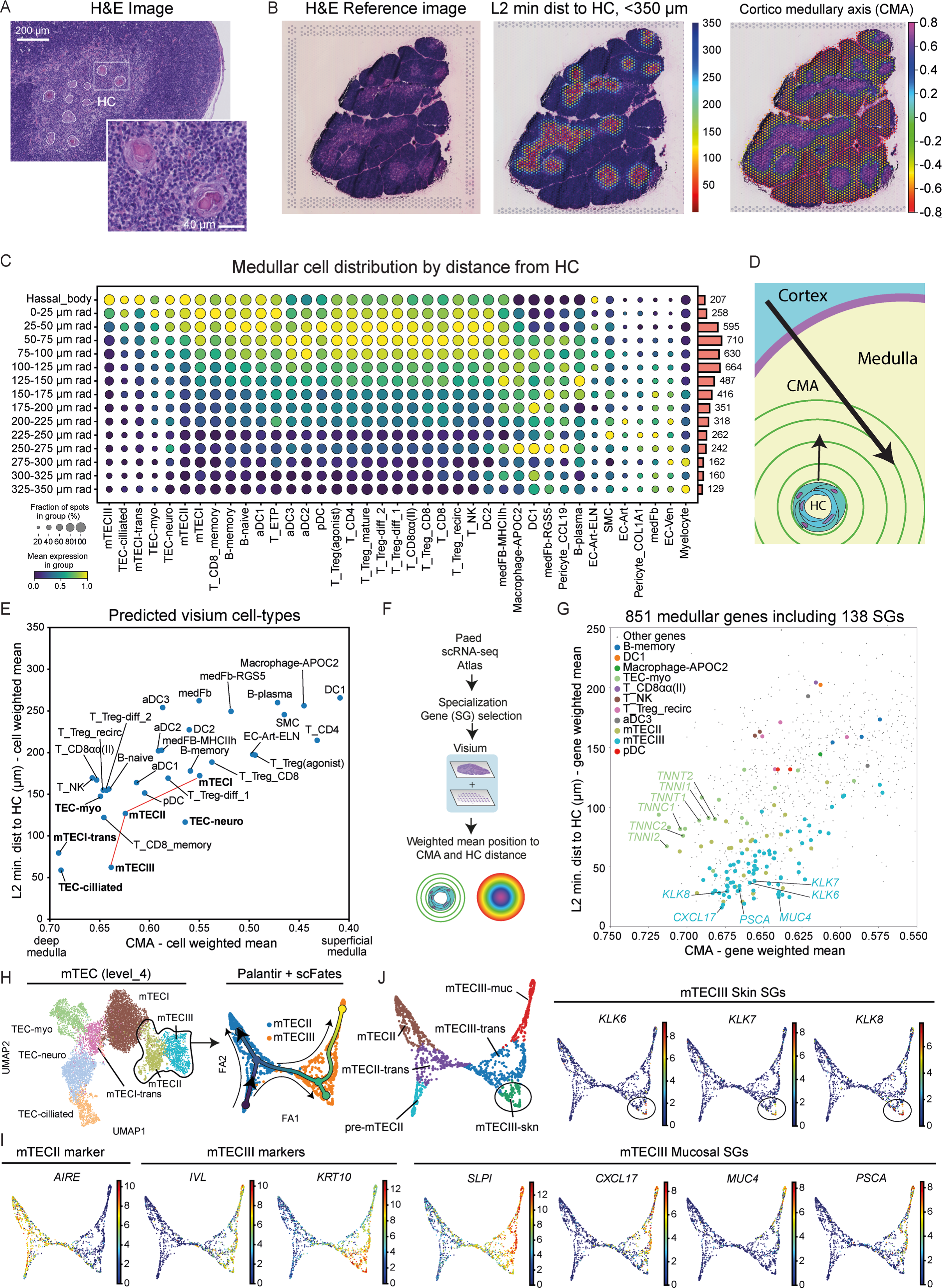
Specialised mTECs are organised around HCs in the paediatric medulla. **A.** Representative H&E Image of a thymus sample from a 7 day-old donor highlighting multiple HCs in the medulla. Scale bars full image: 200 μm, scale bar inset: 40 μm. **B.** Representative Visium H&E reference image (left), overlaid with minimal distance per spot to HC (plotted up to 350 μm for better visualisation, middle), and full CMA (right). **C.** Distribution of cells in the paediatric medulla from Visium deconvolution data. Distance to HC was split into bins of 25 μm. Absolute spot numbers are depicted on the right **D.** Illustration of the discrepancy between the distance to the HC as opposed to the CMA. While the CMA and medullary depth are parallel, the distance to the HC is dependent on the local spatial distribution of the nearest HC. **E.** Weighted mean position of medullary cells along the two axes based on deconvolved Visium data. Note that mTECIII are closest to the HC while not the deepest with respect to the CMA. **F.** Schematic of the workflow for identification of cell specialisation genes (SGs) and their spatial mapping based on Visium data. **G.** Distribution of all 851 medullary genes along the CMA and with respect to HC distance. SGs are represented by large dots, which are coloured according to the cell type they are uniquely expressed in. **H.** UMAP embedding for mTEC lineage with annotation of subtypes highlighting the mTECII/mTECIII branch (left). Palantir embedding and scFates trajectory analysis of the mTECII/III lineage (right). Colour indicates time. Small arrows were manually added for illustration of direction. **I**. Trajectory embeddings showing the expression of canonical markers of mTECII (*AIRE*) and mTECIII (*IVL, KRT10*). **J.** Annotation of differentiated mTECII/III states guided by skin and mucosal SGs, which are spatially enriched around HCs (see Supplementary Figure 5 for further information).

To explore this, we used TissueTag to annotate HCs from the high-resolution reference Visium images and to determine the distance from each spot to the edge of the nearest HC (**Figure 5B**). In contrast to the CMA, which is non-linearly anchored by three reference landmarks (capsule, cortex, medulla) and describes relative localisation, the HC distance is linearly dependent on a single landmark and is hence better suited for studying proximal associations with different cell types. Taking into account the average size of the medullary regions that contain HCs, we had to limit the range of distances examined to a maximum of 350 μm to include only medullary spots and we further decreased our range to 0-150 μm to consider biologically meaningful interactions, given the close proximity between individual HCs in the medulla (**Figure 5A, B, Supplementary Figure 4A**).

In our analyses, we observed that different mTECs, B cells, aDCs, mature T cells, and NK cells were distributed in proximity to the HCs (**Figure 5C**), but we also noted a general enrichment of these cells in medullary level III (**Figures 3E, 4C, Extended Data Figures 5C, 8C, 8I**). To better resolve associations of cells with the deep medulla vs. to HCs (**Figure 5D**), we calculated the mean weighted CMA position vs. mean weighted HC distance for all medullary cell types, based on the deconvolved paediatric Visium data set (see Methods). Similar to our earlier observations, mTECI showed the most dispersed and superficial localisation (**Figure 4C, 5E**). mTECII were found deeper along the CMA and relatively close to HCs, while mTECIII were the deepest and predominantly located in close proximity to HCs (**Figure 5E**), as expected based on previous literature^63^. In addition, the specialised TEC-ciliated, mTECI-trans and TEC-neuro were close to the HCs after accounting for medullary depth. Of all medullary TECs, TEC-myo were least associated with HCs. Other antigen presenting cell types displayed a more superficial medullary localisation and no direct association with HCs (**Figure 5E**).

We speculated that there might be a link between proximity to the HCs and the degree of specialisation of TECs with respect to their expression of peripheral tissue antigens (PTAs). To address this question, we investigated the medullary genes that were restricted in their expression to a single cell type in our paediatric scRNAseq data set, hereafter referred to as specialisation genes (SGs)(**Figure 5F, Supplementary Figure 4B, see Methods**). Out of 851 genes with medullary expression, 138 genes were identified as SGs. mTECII, which express AIRE and consequently a broad range of PTAs, were found to express a substantial (22) but, surprisingly, not the highest number of SGs. The highest number, 77 SGs, was determined to be expressed in mTECIII, while 18 SGs were found in myo-TECs and only 21 SGs were associated with the remaining medullary cell types (**Supplementary Figure 4B, Supplementary Figure 5A**). This highlights the canonical role of TECs as specialised antigen-presenting cells in the human thymus. Weighted spatial mapping of SGs to the CMA and HC distance revealed that mTECII and mTECIII SGs were highly enriched in the deep medulla but also close to the HC (**Figure 5G**). In contrast, myo-TEC SGs were mapped to the deep medulla but did not show a specific association with HCs. Finally, SGs from the other cell types were detected in more superficial medullary regions and further from HCs (**Figure 5G**). SGs in close proximity to HCs were *CXCL17* and *MUC4*, which may mark a subset of mTECIII expressing mucosal PTAs (**Supplementary Figure 5B**). In addition, a set of kallikrein-related peptidases (*KLK6*, *KLK7*, *KLK8*, *KLK10*) that are involved in skin desquamation^74^ were expressed in a second subset of mTECIII and in proximity to HCs (**Supplementary Figure 5B**). A subset of Epidermal Differentiation Complex (EDC) that are known to be responsible for terminal differentiation and cornification of keratinocytes^75^ were expressed in mTECIII and were among the SGs with highest HC association (**Supplementary Figure 5C**).

To further investigate the relationship between spatial proximity to the HC and mTECIII differentiation state, we performed an unbiased trajectory analysis using the scFates package on the combined mTECII and mTECIII subsets and only specified the root. This suggested the existence of mTECII and mTECIII transitioning states and two distinct branches in the mTECIII population (**Figure 5H, I**). One branch population that we termed mTECIII-muc which was characterised by the robust co-expression of SGs, such as *CXCL17* and *SLPI*, as well as *PSCA* and *MUC4* marking the end of the trajectory (**Figure 5J**), which are known to be expressed by the mucosal epithelia of the gut, lung, stomach, kidney and other organs according to the single-cell evidence from the Human Protein Atlas^77^. Of note, CXCL17 has been shown to act as a chemoattractant in the mucosal layers of the respiratory tract^78^. The second branch population, mTECIII-skn, stood out by expression of classical keratinocyte genes, such as *KRT10* and *KRTDAP* as well a number of kallikrein-related peptidases (*KLK6*, *KLK7*, *KLK8*, *KLK10*), which are involved in skin desquamation^75^ and were marking the most differentiated state of that branch (**Figure 5J**).

In summary, we find that multiple medullary cell types are found in proximity to HCs, as has been suggested previously. However, by accounting for the medullary depth and considering biologically meaningful distances, we show that mTECII and mTECIII are the most strongly associated with HCs whereas myo-TECs are further apart from HCs and more associated with medullary depth. We further find evidence that within the mTECIII subset, additional differentiated subtypes expressing mucosa- (mTECIII- muc) and skin-related PTAs (mTECIII-skn) can be identified, which also show strong association with HCs.

### CITE-seq permits fine grained annotation and high-resolution spatial mapping of the T lineage

Traditionally, studies of thymic T cell development have relied on surface markers for flow cytometric or imaging-based spatial analysis. While scRNA-seq provides detailed insights into continuous cell states, quantitative differences in abundance as well as delay between transcription and cell surface expression of marker genes impede relating antibody-derived findings to RNA-based cell annotations. To overcome this issue, we performed CITE-seq, which enables paired quantification of transcripts and surface protein levels at single cell resolution. We used a 143-plex customised antibody panel to cover a broad range of cell surface markers commonly used in thymocyte research (**Supplementary Table 7**) and further combined this with scTCR-seq to leverage insights into the *TRA* and *TRB* locus rearrangement status in T lineage cells. Through this approach we were able to annotate T cell developmental stages with higher resolution than in the scRNA-seq data set (**Extended Data Figure 11A, B**) and substantially increased the number of identified discrete differentiation stages compared to the few other studies that have recently applied CITE-seq to the human thymus^16,44^. To additionally gain spatial information about the identified cells, we mapped the annotated subsets to paediatric Visium data to determine their distribution along the CMA.

Based on the expression of well-known surface markers, such as CD1A, CD34, CD44 and CD4^8,76^, we annotated multiple differentiation stages of immature thymocytes, including cells before (“uncommitted”) and after T lineage commitment (“committed CD4neg”) and ISP cells (“committed CD4pos”). We identified a subset of quiescent DP thymocytes, which was characterised by a low frequency of *TRB* rearrangements and CD31 expression (**Extended Data Figure 11A-C**) and thus resembles a previously described CD31^hi^ CD4^+^ CD8a^+^ CD8β- stage, that has been associated with TCR β-selection^77^. All of these immature thymocyte subsets were determined to be predominantly resident in the capsule, subcapsular regions and the outer cortex of the paediatric thymus (**Extended Data Figure 11D**). In addition, uncommitted thymocytes also showed some enrichment in the upper layers of the medulla, which aligns with the mapping of ETPs to the medullary layers seen in our scRNA-seq based annotations (**Extended Data Figure 11D**).

We were further able to resolve cells that had undergone positive selection (CD69^hi^, *RAG1/2*^lo^), which was followed by a CD4^hi^CD8^lo^ stage, during which cells have already received the signalling cues instructing their lineage fate but do not yet exhibit a clear lineage-specific transcriptional programme (**Extended Data Figure 11B, E**)^78,79^. Both of these post-selection stages were still predicted to be localised in the cortex but already showed a shift in their spatial distribution towards the CMJ (**Extended Data Figure 11D**). Following the lineage bifurcation, CD4+ and CD8+ SP thymocytes could be distinguished in their maturation progress based on commonly used markers, e.g. CD1A, CD69, CD27, CD45RO and CD45RA^8^ (**Extended Data Figure 11B**). Spatial mapping indicated a change in localisation to the thymic medulla for all subsets except immature CD8SP thymocytes, which were still mainly detected in the cortex (**Extended Data Figure 11D**). The CITE-seq profiling was critical for the distinction of several Treg stages and subsets, such as CD103+ CD8+ Tregs^80^ as well as recirculating Tregs (CD39+ CD31-, “Treg_recirc”)(**Extended Data Figure 11A, 12A**), all of which were predicted to reside in the medulla (**Extended Data Figure 12B**). We were further able to annotate several unconventional T cell subsets, such as CD8αα(I) and CD8αα(II) cells, as well as differentiating and recirculating yδT cells, with immature cells mapping to the cortex/CM, and mature cells found in the medulla (**Extended Data Figure 11A, 12A, 12B**). Lastly, we detected several subtypes of NK cells that were predicted to reside in the thymic medulla as well as putative developing NKT cells (CD4+ CD8+ CD56+ CD161+, “NKT_dev”) that mapped closer to the CMJ in accordance with their DP phenotype and origin (**Extended Data Figure 12B**).

### CD4 and CD8 T cell lineages show diverging patterns of cortico-medullary migration

Overall, the spatial mapping of T lineage cells using the CITE-seq-derived annotations confirmed the previously reported trajectory, with most immature and mature stages predominantly localised in the medulla, in contrast to differentiation, proliferation, and selection of DN and DP thymocytes that appears to take place mostly in the cortex and capsule (**Figure 3E**). Through the distinction of immature, semi-mature, and mature stages for thymocytes of the CD4 and CD8 lineage, we were able to map cells at this final stretch of thymic differentiation with increased resolution. This suggested a difference in the localisation of immature CD8 vs. immature CD4 thymocytes (**Extended Data Figure 11D**), which prompted us to further leverage our multimodal data sets to investigate the cortico-medullary transition of thymocytes following their positive selection and lineage decision in more detail (**Figure 6A**). For this purpose, we carried out trajectory analysis on the weighted combined RNA and protein modalities in the CITE-seq data to predict the differentiation pseudotime of CD4 and CD8 lineage cells from initiation of positive selection to full maturity (**Figure 6A-C**, see Methods). Additionally, to obtain more continuous spatial mapping of the cells, we used high-resolution Leiden clustering to unbiasedly group cells according to their gene expression profiles. These clusters were mapped to the paediatric Visium data with cell2location to obtain mean CMA values of each cluster, which were then transferred back onto the cells comprising the different clusters (see Methods). This approach enabled us to locate cells in both space and developmental time, independent of discrete annotations, and thus explore the relationship between differentiation and migration.

**Figure 6.**
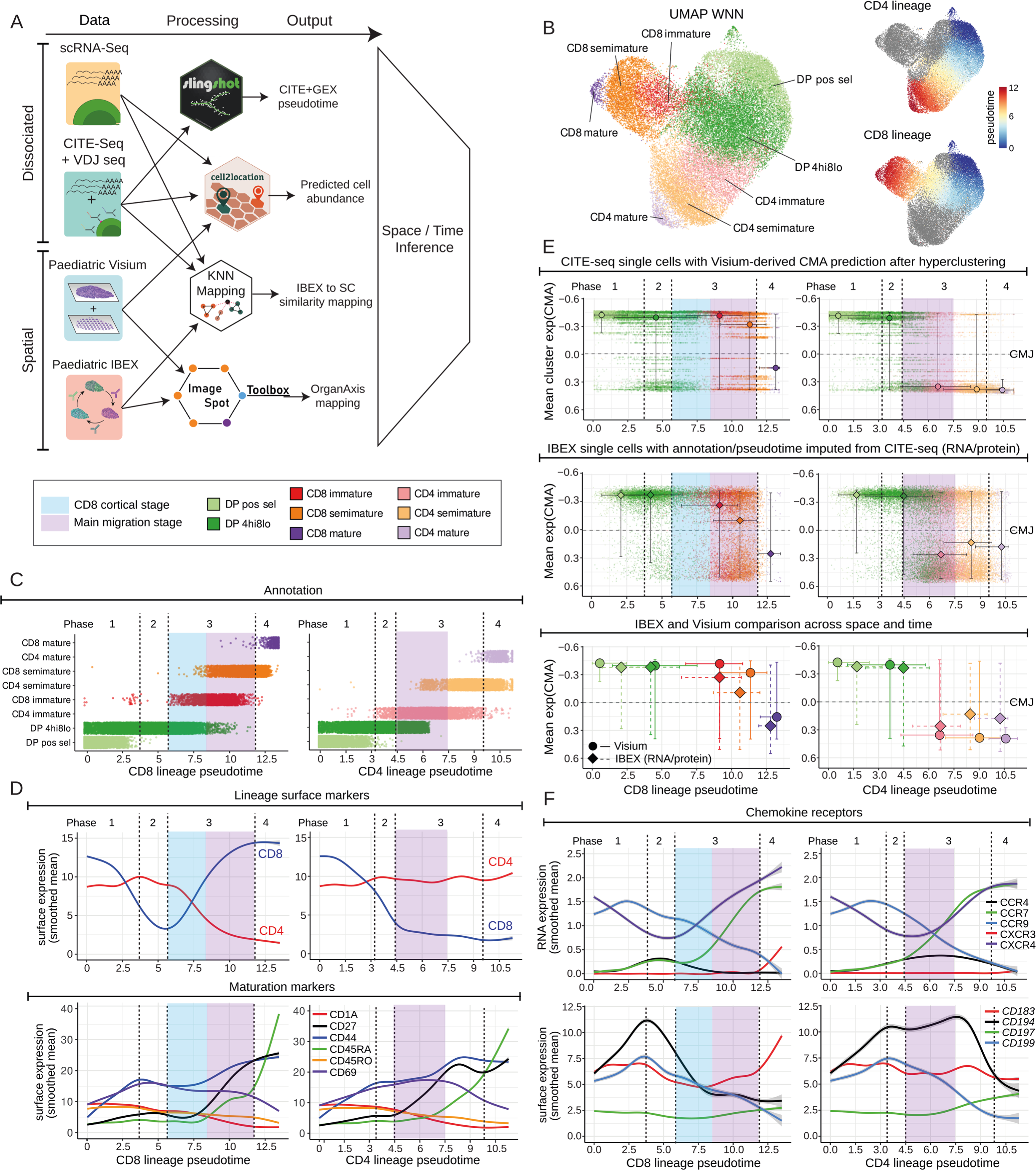
Multimodal high-resolution annotation and spatial mapping of CD8 and CD4 lineage cells reveals differences in the cortico-medullary migration. **A.** Workflow of feature extraction and integration of multimodal datasets. Denoised surface protein levels and RNA expression from CITE-seq together with TCR-seq data were used to call T lineage maturation stages in the paediatric thymus and carry out trajectory analysis for the CD4 and CD8 lineages using Slingshot. Integrated CITE-seq and scRNA-seq data were used for cell abundance prediction in Visium data using cell2location and as a single cell reference for KNN similarity mapping and annotation of IBEX nuclei segmentations. OrganAxis mapping on paediatric Visium and IBEX sections was carried out with ImageSpot. **B.** WNN UMAP representation of paediatric CITE-seq data based on RNA and cell surface protein expression for conventional αβ T lineage cells from positive selection to full maturity (left) and pseudotimes for the CD4 and CD8 lineage obtained with Slingshot (right). **C.** Cells ordered along CD4 (right) and CD8 lineage pseudotime (left) with colour indicating discrete annotations. Four pseudotime phases were derived from the cytokine receptor expression profiles in (D); cortical (blue) and migration stage (purple) were identified using spatial mapping in (E). **D.** Analysis of cell surface protein levels of lineage and maturation markers along predicted pseudotimes for CD4 (right) and CD8 lineage (left). Line plots represent smoothed means and standard error of surface protein levels in cells shown in (C). **E.** Localisation of CD4 (right) and CD8 lineage cells (left) within the thymus lobule as predicted by integration of spatial and CITE-seq data. Top panel shows individual cells in the dissociated dataset with inferred CMA from Visium data after hyperclustering. Medians and 0.05/0.95 quantiles for CMA value and pseudotime of the discrete annotated substages are also shown. Middle panel shows individual cells in the IBEX data after CMA calculation and inference of cell annotation/pseudotime through KNN similarity mapping with the CITE-seq protein/RNA data serving as reference. KNNf = 0.5 was used as a cutoff for cells to be included. Medians and 0.05/0.95 quantiles for the discrete annotated substages are also depicted. Bottom panel provides a direct comparison of the medians/quantiles shown in the top and middle panel for Visium (circle) and IBEX mapping (diamond). **F.** Analysis of RNA and protein levels of chemokine receptors. Line plots represent smoothed means and standard error of surface protein or RNA levels in cells shown in (C).

For easier visual interpretation and comparison, we segmented the differentiation pseudotimes of each lineage into four matching phases based on the surface expression of CD4, CD8, and various maturation markers and beginning with the earliest stage seen upon initiation of positive selection: (1) CD4/8 inversion and increase in CD69 and CD44; (2) continuous CD8 downregulation and plateau in maturation marker expression; (3) adoption of the lineage-specific CD4/CD8 profile and increase in CD27 and CD44; and (4) strong increase in CD45RA and drop in CD69 (**Figure 6D**). We found that cells in phase 1 that had just passed positive selection were still clearly located in the cortex. In the subsequent CD4^hi^ CD8^lo^ stage some cells had already transitioned to the thymic medulla, although the majority of cells at this lineage bifurcation stage were still located cortically, as indicated by the median CMA value (**Figure 6E**, top panel). CD4SP immature thymocytes were enriched in the medulla, as were CD4 semi-mature and CD4 mature cells, indicating that for this lineage the cortico-medullary migration predominantly occurs in early phase 3 and thus in parallel with the induction of the lineage-specific transcriptional and phenotypic programme (**Figure 6E**, top right panel, purple shading). In contrast, the majority of immature CD8 cells were still predicted to remain in the cortex in early phase 3, while they underwent co-receptor reversal to establish their complete CD8+ CD4- phenotype (**Figure 6D, E**, top left panel, blue shading). Similarly, semi-mature CD8 cells in late phase 3 were still found in the cortex in substantial numbers and only mature CD8 cells in phase 4 were finally primarily detected in the medulla (**Figure 6E**, top left panel).

To further validate this relative delay in the cortico-medullary transition of CD8 lineage cells, we sought to identify the localisation of the corresponding cells in the IBEX imaging data. To this end we used our previously applied KNN matching approach to annotate IBEX T cells and infer developmental pseudotimes using the CITE-seq data as a reference. Since a delay occurs between changes in RNA and protein expression, which can be particularly noticeable in differentiating cells that continuously change their transcriptional and phenotypic profiles, we leveraged the fact that 19 protein targets were covered by both the CITE-seq and IBEX antibody panels. This allowed us to directly compare protein expression for these markers, while the corresponding transcript levels were used for the remaining IBEX targets, yielding an RNA/protein hybrid reference data set to ensure accurate cell annotation of IBEX single cells (**Figure 6A, Supplementary Table 4**, see Methods). Remarkably, this cell type inference and pseudotime imputation resulted in a predicted spatial distribution of CD4 and CD8 lineage cells that was highly similar to that previously obtained from Visium data (**Figure 6E**, middle panel). CD4^hi^ CD8^lo^ cells were predominantly found in the thymic cortex, whereas CD4 lineage cells seemed to transition to the medulla at the immature stage in early phase 3. This was determined to happen considerably later for CD8 lineage cells at the semi-mature stage in late phase 3, as predicted by the Visium-based mapping (**Figure 6E**, middle panel, purple shading).

Next, we aimed to explore which chemokines may be involved in this lineage-specific migration pattern. In line with previous studies demonstrating a role of CXCR4 in cortical retention and transient downregulation of the chemokine receptor (CR) after positive selection^81^, we observed an initial drop in the RNA levels of this receptor in phase 1-2 in both lineages, followed by continuous upregulation from phase 3 onwards (**Figure 6F**). Similarly, *CCR9* (encoding CD199) peaked in phase 1/2 and was subsequently downregulated in both CD4 and CD8 lineage cells, as described previously^82^. Notably, the corresponding chemokines for these two receptors, *CXCL12* and *CCL25*, were strongly expressed in the capsule and cortex but not in the medulla, which suggests contributions to localisation of DP thymocytes to the cortex (**Extended Data Figure 12C**). *CCR7* (encoding CD197), which mediates responses to CCL19 and CCL21, is essential for medullary homing and negative selection^81,83,84^. This CR was upregulated in early phase 3 for CD4 lineage cells but only in late phase 3 for the CD8 lineage, coinciding with the main medullary migration window of each lineage (**Figure 6F**). This delay in *CCR7/*CD197 expression during CD8 differentiation matches observations in a recent mouse CITE-seq study^85^ and points to a potential role of this receptor in the timing of the cortico-medullary transition.

Expression of *CCR4*/CD194 increased throughout phase 1 in CD4^hi^ CD8^lo^ cells, but while cells of the CD4 lineage remained CD194-high until the end of their migration window, CD8 lineage thymocytes exhibited a swift reduction in the receptor RNA and protein levels in phase 2-3 (**Figure 6F**). Based on the known function of *CCR4*/CD194 in the cortico-medullary transition^81,83^ the diverging expression patterns in CD4 and CD8 lineage thymocytes suggest that CD194 levels and the resulting responsiveness to medullary CCL17 and CCL22 may also be a driving force behind the earlier medullary migration of CD4+ thymocytes. These observations are matched by those in a recent report on mouse thymocytes, which describes the staggered upregulation of CCR4 and CCR7, resulting in the migration of immature and mature SP thymocytes, respectively. In line with our results, the authors further showed a temporal delay in the differentiation of CD8 lineage cells after the TCR stimulus, downregulation of CCR4 upon co-receptor reversal, and delayed medullary accumulation compared to CD4 lineage thymocytes^83^.

Finally, *CXCR3* (encoding CD183) was not detected in CD4 lineage cells but was upregulated in the CD8 lineage in phase 4, which could indicate a role in the late stages of medullary entry or even intra-medullary migration for this lineage (**Figure 6F**). Alternatively, CD183 expression on mature CD8 T cells could be related to gut homing, since a population of CXCR3+ CD45RA^hi^ CDαβ+ human thymocytes with properties of CD8αα+ gut intraepithelial lymphocytes (IEL) has been reported previously^86-88^.

In summary, by using two highly different spatial technologies in conjunction with a multimodal single cell reference atlas we could reproducibly detect a substantial difference in the timing of thymocyte migration from cortex to medulla for the CD4 vs. CD8 lineage cells that matched differing expression profiles for the medullary homing receptors *CCR4*/CD194 and *CCR7*/CD197.

## Discussion

Large scale efforts of international consortia like the HCA and HuBMAP have emphasised the need for constructing extensive tissue atlases to offer a basis for deeper understanding of cell and human biology. Initially consisting of analyses of dissociated cells, next generation spatial atlases now provide crucial insights into cell interactions and structural underpinnings of complex tissues and organs. However, harmonisation of spatial datasets remains a challenge and represents a major hurdle for multi-group consortia. Here, we offer a new integrative computational approach to spatial transcriptomics analysis and implement these novel tools to both spatial transcriptomics and multiplex protein imaging methods. Combined with newly generated multimodal single-cell data and previously published data sets, we have created a thymus spatial atlas spanning fetal and early paediatric life that was curated specifically to cover developing T cells as well as thymic hematopoietic and stromal cell subsets and states at higher resolution and diversity than previously achieved.

By conducting CITE-seq and scTCR-seq we have gathered whole transcriptome readouts, protein expression of 143 cell surface markers, and TCR rearrangement status for over 30 stages of conventional and unconventional T cell development in the human paediatric thymus. This trimodal data set can serve as a critical bridge between classic methodological approaches and recently developed methods, e.g., antibody-based applications, such as flow cytometry or immunofluorescence imaging, and nucleic acid-based approaches, like scRNA-seq or spatial transcriptomics, through imputation of the missing modality. We further integrated a large cohort of fetal and early paediatric Visium spatial transcriptomics data and RareCyte multiplex imaging with an extensive paediatric IBEX 44-plex imaging data set based on a custom TEC- and T cell-focussed antibody panel to chart both micro- and macroscopic tissue architecture and spatial cellular compositions. These data can be used to directly compare protein and RNA for cell types in their local environment as well as larger structural anatomical features. Beyond these reference datasets, we have developed an extensive computational tool set for annotation, processing, and analysis of imaging and spatial transcriptomics data (TissueTag and ImageSpot). All of these data sets and computational analysis pipelines will be made available to the broader community for further application, exploration and analysis.

The thymus is a dynamic organ, in which the majority of cells are constantly moving, while the organ itself can undergo rapid macro-morphological changes within the course of days, e.g., during infection, as well as more gradually during ageing. Changes to the relative size of various classically defined compartments within the organ as well as the migration patterns of the constituent cells make it difficult to determine the preferred spatial location of cell types and distinguish random from consistent spatial associations. Here, we establish a new spatial approach, OrganAxis, which leans on a combination of distance functions derived from tissue landmarks to generate a rotation- and scale-invariant, continuous CCF for the thymus, the Cortico-Medullary Axis (CMA). The CMA maps the relative position of a given cell or spot within the thymus and thus permits biological comparisons across distinct spatial data sets obtained from different donors, experimental approaches and institutes. Since the CMA is expression-independent and solely based on morphological landmarks, it does not require cell annotation for the analysis of transcriptional gradients and niches in spatial data. For this reason, our tool set for continuous spatial annotations can be adapted to any tissue that has consistent landmarks and applied to a broad variety of spatial technologies. We show the ability of this axis to bridge technical, sample, technique and developmental gaps while preserving biological diversity. By comparing cell/gene distributions along the CMA between IBEX and Visium data we were able to conduct highly useful cross-platform validation of our observations. At the same time, it is important to note that despite the ability of the CMA to regularise comparisons across samples that vary in size and overall cellularity, our study of the earliest stages of T cell development points to a need for additional refinement of the approach to more fully account for issues such as the relative proportion vs. absolute cell count in an anatomical region.

Because it is well established that the thymus undergoes dramatic changes over an individual’s lifetime, culminating in the onset of adipose involution around adolescence, we focussed our analyses on fetal and early paediatric periods, during which the organ is functioning at peak capacity in terms of T cell generation. We found that, despite large-scale macrostructural deviations between the fetal and paediatric thymus, the spatial maturation trajectories and major TEC distributions are established as early as pcw 12 and maintained over the first year of post-natal life. This ontologic conservation of trajectories was also seen in transcriptional patterns of key chemokine genes, whose products guide the thymocytes within and between compartments. Future studies using many of the tools we introduce here and the baseline spatial atlas emerging from our work will be essential to chart the changes during later life and the mechanisms controlling the involution process.

While the patterns of cell movement were largely preserved across age, our detailed investigation revealed an unexpected difference in the timing of cortico-medullary migration of maturing conventional CD4+ vs. CD8+ αβ T cells after positive selection. We show that the pattern of downregulation of *CCR4*/CD194 and upregulation of *CCR7*/CD197 differs as cells mature along the two opposing developmental trajectories, whereby a CD8 lineage-specific CD194-CD197- phase substantially retarded the migration of developing CD8SP thymocytes from the cortex to the medulla. These differences were seen in both Visium and IBEX data, demonstrating how multimodal data sets can be employed to derive cross-validated interpretations regarding a cell’s maturation state, its local environment, and its position in a tissue. Our findings directly align with two recent studies that suggest a slower maturation of CD8 lineage cells^83,85^. Moreover, our observation that co-receptor reversal in CD8 lineage cells coincides with the cortical retention window further indicates that CD8SP thymocytes may require more time to initiate the lineage-specific differentiation programme, which in turn could impact the timing of intercompartmental cell migration. Additionally, the cortex may provide the specific microenvironment and signalling cues that CD8 lineage cells need to initiate their maturation. Finally, in accord with Li et al.^83^ we found that ligands for CD194 and CD197 were predominantly expressed on hematopoietic antigen-presenting cells and mTECs, respectively, which implies that the observed differences in CD194 and CD197 expression between the two T cell lineages may influence which medullary cell types attract and interact with CD4SP vs. CD8SP thymocytes.

The maturation and selection of T cells in the thymus is critically supported by TECs and the origin and location of TEC progenitors needs to be identified to understand thymus growth or to potentially leverage this information for therapeutic applications. Our data revealed the fetal capsule/subcapsular region as a niche that houses TEC progenitor proliferation, organ growth, and vascularisation. Postnatally, many of these cell types and processes seem to be shifted to the PVS and medulla, suggesting the possibility that the specialised fetal subcapsular region is remodelled to form the paediatric capsule and PVS during late stages of fetal development. However, because our data does not cover the 3rd trimester of fetal development, further studies are needed to carefully explore this concept.

Interestingly, the fetal deep cortex or “cortical layer ll/III” was distinct from that in the paediatric thymus in cTECII distribution as well as late DP cells, CD8αα(entry) and yδ T cells. In addition, fetal cTECII show the strongest expression of MHCII, which resembles perinatal cTECs in mouse that have upregulated MHCII and LY51 and were recently shown to localise in deep cortical regions and mediate positive selection^89^. Perinatal cTECs decrease with age, which may explain the disappearance of this deep cortical niche in the post-natal human thymus. In contrast to these changes between fetal and paediatric samples in terms of the cortex and more in line with our observations on thymocyte maturation, there was a conserved level of internal organisation of the thymic medulla in both the fetal and the paediatric thymus.

The medulla is the site where T cells are screened for their potential to recognise auto-antigens through numerous interactions with PTA-presenting cells and our analyses suggest that the various hematopoietic and stromal cells involved in this task show a distinct structural organisation. Several studies have noted an enrichment of various antigen-presenting cells in proximity to HCs and the latter have been implicated in the induction of tolerance. This led us to investigate whether the medullary organisation and HCs are linked. By distinguishing between medullary depth and proximity to HCs, we found that specific gene families, which are expressed by subsets of *AIRE+* mTECII and post-AIRE mTECIII, were highly associated with the HCs. We further found two mTECIII differentiated keratinocyte sub-lineages representing distinct body boundaries: mTECIII-skn (*KLK*+, *KRT10*^hi^), which show similarities to terminally differentiated skin keratinocytes in line with the previously suggested role of HC cells, and mTECIII-muc (*MUC4*+, *CXCL17*+, *PSCA*+), which is defined by mucosa-related genes. Both specialised mTECIII subsets were found in very close proximity to HCs. This level of medullary organisation might hint at how negative selection occurs in an efficient manner capable of promoting central tolerance by rare mimetic cells and/or rarely expressed PTAs under AIRE control.

Overall, we report here a large collection of reference data for the fetal and paediatric human thymus and demonstrate that these data both reconcile prior findings while also leading to a series of novel biological insights in the areas of stromal cell organisation and T cell maturation and migration. We expect that beyond the observations we detail here, there is still much to uncover using these data sets. We have further illustrated the usefulness and applicability of the CMA and hope that this CCF can support future thymus studies by offering a common language of reference for the community, especially when discussing hard to define structures like the CMJ and subcapsular region. Finally, the computational tools and specifically OrganAxis translates well to other organ systems in 2D, 3D and even 4D (time), enabling accurate spatial modelling and CCF construction, hopefully providing valuable resources to the broad scientific community.

## Acknowledgements

First and foremost, we would like to thank the donors and their families for their consent and contribution for this study. The supporting teams of the Teichlab wetlab, CellGenIT, CellGen wetlab and Spatial Genomics Platform teams (SGP) have been central for this study. We thank Dr. Ziv Yaniv for computational support for IBEX image alignment. We gratefully acknowledge the Sanger Flow Cytometry Facility for support with sample sorting. We thank Ellen De Meester (NXTGNT, Ghent University) for help with single cell sample preparation and sequencing. We also thank Prof. Bart Vandekerckhove (Ghent University) for providing thymus samples through the hematopoietic cell biobank.

Finally, we would like to dedicate this publication to our beloved friend and colleague, Daniele Muraro, who tragically passed away last year.

This study was primarily funded by the Chan Zuckerberg Foundation by grant number 2019-002445 and 2022-249170 (doi: 10.37921/644286wvygde). The Wellcome Sanger Institute is supported by core funding from the Wellcome Trust (220540/Z/20/A). Work in the Taghon lab was supported by the Fund for Scientific Research Flanders (FWO, grant G075421N and fellowship 12D9523N to L.B.) and the Concerted Research Action from the Ghent University Research Fund (GOA, BOF18- GOA-024). Work at the NIH was supported, in part, by the Intramural Research Program of the NIH, National Institute of Allergy and Infectious Diseases (NIAID) and National Cancer Institute (NCI). All research at GOSH is supported by the UK National Institute of Health Research and Great Ormond Street Biomedical Research Centre, AYK Wellcome Trust (222096/Z/20/Z) EGD GOSH Children’s Charity.

## Author Contributions

N.Y. wrote the paper, conceived the study, performed analysis, generated single cell data, generated spatial data, conceived spatial modelling, conceived methodology, contributed software, and collected biological samples. V.R.K. wrote the paper, conceived the study, performed analysis, generated single cell data, generated spatial data, conceived methodology, provided key biological interpretation, and collected biological samples. L.B. wrote the paper, conceived the study, performed analysis, generated single cell data, conceived methodology, provided key biological interpretation, and collected biological samples. C.S. performed analysis, generated single cell data, conceived methodology, and provided key biological interpretation. B.W. and R.T.B. generated spatial data, provided key biological interpretation, and collected biological samples. O.A. conceived methodology, contributed software, and provided computational support. K.P. performed analysis and provided computational support. S.K. performed analysis, conceived methodology, and contributed software. E.T. generated spatial data. E.D. conceived methodology and provided computational support. J.V.H. performed analysis and generated single cell data. S.P. generated spatial data. T.P. performed analysis. A.P. curated published data and provided computational support. M.D. generated spatial data. L.R. generated single cell data. C.T. generated spatial data. A.Y.K. provided key biological interpretation and collected biological samples. J.E. collected biological samples. E.S. collected biological samples. V.K. conceived methodology and provided computational support. F.D.R. collected biological samples. M.B. provided key biological interpretation and collected biological samples. F.P. provided key biological interpretation and collected biological samples. E.P. generated single cell data. N.-J.C. generated spatial data. M.P. provided computational support. K.T. contributed software. L.F. contributed software. R.A.B. and X.H. provided key biological interpretation and collected biological samples. D.C. collected biological samples. F.V.N. generated single cell data. M.P. generated spatial data. G.D. collected biological samples. M.A.H. collected biological samples. L.D.N. provided key biological interpretation and collected biological samples. V.U. conceived spatial modelling and conceived methodology. R.N.G. wrote the paper, conceived the study, conceived methodology, provided key biological interpretation, provided computational support, and supervised/managed the project. A.J.R. wrote the paper, conceived the study, performed analysis, generated spatial data, conceived spatial modelling, conceived methodology, provided key biological interpretation, and supervised/managed the project. J.C.M. wrote the paper, conceived the study, conceived spatial modelling, conceived methodology, provided key biological interpretation, provided computational support, and supervised/managed the project. T.T. wrote the paper, conceived the study, conceived methodology, provided key biological interpretation, collected biological samples, and supervised/managed the project. S.A.T. wrote the paper, conceived the study, conceived spatial modelling, conceived methodology, provided key biological interpretation, and supervised/managed the project.

## Data availability

The annotated fetal and paediatric integrated atlas and Visium objects for this study will be available to the public following publication.

Sequencing data for the newly generated libraries as well as human thymus scRNA-seq that was remapped from publicly available repositories will be deposited to Array Express and accession codes will be available before publication. Publicly available data sets were downloaded from the following sources: Park et al.6: ArrayExpress, accession ID E-MTAB- 8581; Campinoti et al.^30^: GEO, accession ID GSE159745; Bautista et al.^17^: GEO, accession ID GSE147520. Prior to publication data will be shared upon specific request under collaborative agreement.

### Code availability

All code scripts and notebooks used in the study are open and available to the public. Manuscript - https://github.com/Teichlab/thymusspatialatlas and TissueTag - https://github.com/nadavyayon/TissueTag with one Visium sample as an example code.

## Materials and Methods

### Data generation by institute

Metadata about single-cell RNA-seq samples can be found in **Supplementary Table 1**. Information about spatial data, including Visium, IBEX, RareCyte and RNAscope can be found in **Supplementary Table 2**. Briefly, all CITE-seq data was generated at Ghent University and mapped to the human genome (GRCh38) at the Wellcome Sanger Institute (WSI). All non-CITE-seq original single-cell and Visium data sets were generated and mapped to the human genome (GRCh38) at WSI. All IBEX imaging was performed at the National Institute of Allergy and Infectious Diseases (NIAID), NIH. All non-IBEX imaging data sets were generated at WSI. No fetal work was performed at the NIH and at Ghent University.

Human fetal and paediatric data from several previous studies^6,17,30,31^ was included and reanalysed; please check the respective publications for details on sample processing, ethics and funding. Paediatric samples were provided by Newcastle University collected under REC approved study 18/EM/0314 and Great Ormond Street Hospital under REC approved study 07/Q0508/43. The human embryonic and fetal material was provided by the joint MRC & Wellcome Trust (Grant # MR/006237/1) Human Developmental Biology Resource (http://www.hdbr.org). Samples processed at Ghent University were obtained according to and used with the approval of the Medical Ethical Commission of Ghent University Hospital, Belgium (EC/2019-0826) through the hematopoietic cell biobank (EC-Bio/1-2018). Samples processed at the NIH were obtained (under NIAID MTA 2016-250) from the pathology department of the Children’s National Medical Center in Washington, DC, following cardiothoracic surgery from children with congenital heart disease. Use of these thymus samples for this study was determined to be exempt from review by the NIH Institutional Review Board in accordance with the guidelines issued by the Office of Human Research Protections. Informed consent was obtained from all donors or their legal guardians.

#### Sample processing and library preparation for 10x sc/snRNA-seq

The following protocols apply to sample processing at WSI. Human paediatric samples were obtained from cardiac corrective surgeries. Removed thymi were directly moved to HypoThermosol(R)(Sigma-adrich H4416-100ML), shipped by courier with ice packs and processed in under 24 h post surgery. Upon arrival, samples were separated for embedding in Optimal Cutting Temperature (OCT) compound, formalin fixation and paraffin embedding (FFPE), and single cell dissociation as described in the following protocol: https://www.protocols.io/view/human-thymus-single-cell-dissociation-protocol-tei-bx8sprwe

**Single-cell processing.** Briefly, tissue was finely minced and cell dissociation was performed using a mixture of liberase TH (Roche, 05401135001) and DNase-I (Roche, 4716728001) for ∼30 min in two rounds. Digested tissue was filtered through a 70 δm strainer and digestion was stopped with 2% FBS in RPMI media. Next, red blood cell lysis was performed on a cell pellet using RBC lysis buffer (eBioscience, 00-4333-57), after which cells were washed and counted. Afterwards, magnetic, FACS sorting or both were performed to enrich stromal populations. Magnetic sorting to enrich for EPCAM+ or to deplete CD45+ or CD3- cells was performed for U09, U48 and Z11 samples (Supplementary Table 1) using a magnetic sorting kit from Miltenyi Biotec including LS Columns (130-042-401) and the following bead-tagged antibodies: CD45 MicroBeads, human (130-045-801), CD326 (EpCAM) MicroBeads, human (130-061-101), CD3 MicroBeads, human (130-050-101). Some samples underwent FACS sorting to enrich CD45- stromal cells (Z11 and U48) and total TEC cells (U48) (Supplementary Table 1). To perform FACS sorting, cells were resuspended in the FACS buffer (0.5% FBS and 2mM EDTA in PBS), underwent blocking in TruStainFcX (Biolegend, 422302) for 10 min and were stained with a mixture of EPCAM (anti-CD326 PE, clone 9C4, Biolegend #324206), CD45 (anti-CD45 BV785, clone HI30 (mouse), Biolegend #304048), DEC205 (anti-CD205 (DEC-205) APC, Biolegend #342207) and CD3 (anti-CD3 FITC, clone OKT3 (mouse), Biolegend #317306) antibodies and DAPI for 30 min. Upon staining, cells were washed and analysed using the Sony SH800 or Sony MA900 sorters with a 130 μm nozzle. CD45- cells were sorted as a total stroma, while EPCAM+ or CD205+ were sorted to obtain the total TEC fraction including cortical and medullary epithelial cells after standard QC, including debris and dead cell removal (**Supplementary Figure 6A**). Importantly, multiplet filtering was not performed to ensure large thymic epithelial cells (thymic nurse cells) can be included. After enrichment, cells were resuspended to the recommended concentration (10^A^6/1 ml) and loaded on a 10x Genomics chromium controller to generate an emulsion of cells in droplets using Chromium Next GEM Single Cell 5’ Kit v2 (#1000263). GEX and V(D)J libraries were further prepared using the Library Construction Kit (#1000190) and Chromium Single Cell Human TCR Amplification Kit (#1000252) according to the manufacturer’s instructions. Sequencing was performed with a Novaseq6000 sequencer (Illumina). See sample metadata for further details (**Supplementary Table 1, 2**).

### Curation and processing of published scRNA-seq and scTCR-seq data sets

Both newly generated and published data sets were remapped from fastq files and processed as follows: For public data sets deposited on ArrayExpress, paired-end fastq files were downloaded from ENA, and the sdhf file was used to determine the type of experiment (3’/5’ and the version of 10× Chromium kit). For public data sets deposited on GEO, if the data were deposited as a Cell Ranger bam file, URLs for the bam files were obtained using srapath v2.11.0. The bam files were downloaded and converted to fastq files using 10x bamtofastq v1.3.2. On the other hand, if GEO data were deposited as paired-end fastq files, sra files were located using the search utility from NCBI entrez-direct v15.6, downloaded, and converted to fastq files using fastq-dump v2.11.0. Sample metadata was curated from the abstracts deposited on GEO. Finally, samples generated at WSI were downloaded from iRODs v4.2.7 in the form of cram files, and converted to fastq files using samtools v1.12 using the command “samtools collate -O -u -@16 $CRAM $TAG.tmp | samtools fastq -N -F 0×900 -@16 - 1 $TAG.R1 .fastq.gz -2 $TAG.R2.fastq.gz -”. Sample metadata was obtained using the imeta command from iRODs. Following the fastq file generation, 10x Genomics scRNA-seq experiments were processed using the STARsolo pipeline detailed in the github repository https://github.com/cellgeni/STARsolo. A STAR reference matching Cell Ranger 2020-A for human was prepared as described here: https://support.10xgenomics.com/single-cell-gene-expression/software/release-notes/build#header. Using STAR v2.7.9a and the previously collected data about sample type (3’/5’, 10× Genomics kit version), we applied the STARsolo command to specifying UMI collapsing, barcode collapsing, and read clipping algorithms to generate results maximally similar to Cell Ranger v6: “--soloUMIdedup 1MM_CR -- soloCBmatchWLtype 1MM_multi_Nbase_pseudocounts --soloUMIfiltering MultiGeneUMI_CR --clipAdapterType CellRanger4 --outFilterScoreMin 30”. For cell filtering, the EmptyDrops algorithm employed in Cell Ranger v4 and above was invoked using “--soloCellFilter EmptyDrops_CR”. Option “--soloFeatures Gene GeneFull Velocyto” was used to generate both exon-only and full length (pre-mRNA) gene counts, as well as RNA velocity output matrices. For TCR-seq experiment processing, Cell Ranger v6.1.1 with VDJ reference 5.0.0 (https://support.10xgenomics.com/single-cell-vdj/software/downloads/latest) were used to quantify the single cell amplicon profiling results. Default settings of reference-based “cellranger vdj” command were used. Fastq files were converted to <Sample>_S1_L001_R1_001.fastq.gz format to be compatible with Cell Ranger.

### sc/snRNA-seq quality control, data integration, and annotation

All of the jupyter notebooks used for data QC, preprocessing, integration, and annotations are available at the following repository: https://github.com/Teichlab/thymus_spatial_atlas/blob/main/manuscript/preprocessing/single_cell/ Mapped libraries were subjected to computational removal of ambient RNAwith CellBender version 0.1.0^90^. Next, all data sets underwent the quality control filtering using metrics calculated from scanpy.pp.calculate_qc_metrics: number of genes per cell (more than 400, but less than 6500); percentage of mitochondrial counts (less than 6%), percentage of ribosomal counts (less than 5%). Doublets were annotated using Scrublet^91^ and cells with scrublet_score above 0.4 were removed.

Next, cells were integrated with scVI while removing mitochondrial, TCR, and cell cycle genes^92^ and annotated into major lineages (T, TEC, B, Blood, DC, FB, EC, SMC). Individual cell lineages were then separated and integrated with scVI to perform fine-grained annotation and remove some of the remaining doublets, which were picked up by manual annotation. Cell annotations were assigned based on three strategies: (1) annotations taken from the original data set and were transferred to unlabelled cells using KNN graph, as most cells were integrated from past studies; (2) automatic label transfer using CellTypist^93^ using Developing_Human_Thymus or Pan_Fetal_Human models; (3) by marker genes and literature search. After doublet removal and fine-grained annotations, individual lineage objects were combined together to produce a final object encompassing fetal and paediatric thymus. All annotations were separated into 5 levels of hierarchy from the finest (“cell_type_level_4”) to the broadest (“cell_type_level_0”).

scTCR-seq data was processed with Dandelion^94^ and a detailed notebook can be found on the github repository indicated above. Briefly, a dandelion class object with n_obs was constructed from all combined TCR libraries. Cells were then further subset to only include DP, and SP subtypes which contained all 4 V and J TCR chains: ‘v_call_abT_VDJ_main,j_call_abT_VDJ_main,v_call_abT_VJ_main’,j_call_abT_VJ_ main’. Next, milo neighbourhoods were constructed based on scVI neighbourhood graphs(n_neighbors = 100) and VDJ genes and frequency feature space was calculated based on all chains depicted above. VDJ pseudotimes were calculated with Palantir to determine the VDJ pseudotime (‘pseudotime_nhood_vdj’), probability of a cell to commit to the CD8 lineage (‘prob_T_CD8_nhood_vdj’), and the inverse to CD4 lineage (‘prob_T_CD4_nhood_vdj’). All pseudotimes were migrated back to the single cell object.

### Identification of cell type-specific specialisation genes

To identify genes exclusively expressed in a specific cell type or subset thereof we developed custom python functions. Briefly, starting from raw read count, gene expression was scaled with scipy.stats.zscore() for each tested gene. Cells that showed expression below a cutoff of 0.05 were excluded from further steps. Next, a quantile threshold (>95%) was selected to isolate the cells, which showed the highest expression level of a specific gene. A chi-squared test (scipy chi2_contingency) was performed per gene to identify if a specific cell type was overrepresented compared to all annotated cell types, indicating the gene to be a marker gene. We further focused on genes that were predicted to be expressed only in a single cell type. The core function and parameters used were the following: rare_cell_marker_detection(adata_paed, genes=space_map_hc_filt_exp, quantile=0.95, annotation=’cell_type_level_4’, cutoff=0.05, alpha=1e-50, plot=False,scale=True).

### mTEC trajectory analysis

The scFates package^95^ was used to run trajectory analysis on the combined mTECII and mTECIII cells. First, mTECII and mTECIII were integrated and batch-corrected using scVI and major clusters were re-annotated, including small doublet clusters, which were removed. Next, the object was pre-processed according to recommendations from Palantir^96^ and scVI latent embedding was used as an input to “palantir.utils.run_diffusion_maps”. Tree learning was performed using “scf.tl.tree” by using multiscale diffusion space from Palantir as recommended by scFates. Finally, the “mTECII_early” state, which was characterised by expression of some mTECI markers (*ASCL1*, *CCL21*) together with mTECII markers (*AIRE*, *FEZF2*), was used as a root to compute pseudotime using scf.tl.pseudotime.

### Formulation of a cortico-medullary axis with OrganAxis

Biological tissues are classically subdivided by discrete compartmental niches pre-defined through careful examination by histologists. However, the boundaries and distributions of cells within these discrete structures are determined by orchestrated signalling events during development, differentiation, or migration, following autocrine, paracrine and endocrine signalling driven by molecular diffusion. The OrganAxis approach establishes a continuous axis that accounts for local distances within and between structures, without compromising discrete compartmentalisation. To achieve this, we derive the relative distances of each point from the boundary between structures using the following steps: We first define a spatial sampling frequency and a geometric point space by superimposing a non-overlapping hexagonal grid on each image (**Extended Data Figure 4A**, tissue_tag.generate_hires_grid()). Any discrete tissue annotations are then translated to hexagonal grid units, which is preferable to a pixel-based representation since resolution varies dramatically within and between data sets and would impose computational constraints, which are unnecessary given that the most informative spatial variance of the thymus typically spans ∼1 mm in scale. Nevertheless, the user can control this scale parameter and generate a finer grid via TissueTag. In this study, we generated all grids with a diameter of 15 μm, resulting in no interpoint spacing. Next, we calculated the L2 distance of all points in the grid to the closest corresponding points in each annotation by number of KNN points (tissue_tag.dist2cluster_fast(), Equation 1).

Definition:
1. Let p be any point on an hexagonal grid with spacing — r, p ∈ R^2^
2. Let S be an assembly of p points inside an anatomical structure, S ∈ {Medulla, Cortex, Capsule….}
3. dS^(p)^ is the Euclidean distance between point p, and all points that belong to structure S
4. D_s,p_^[i]^ the sorted (by minimal value) series of dS^(p)^ where i is the index, D_s,p_^[i]^≤D_s,p_^[i+1]^,∀i

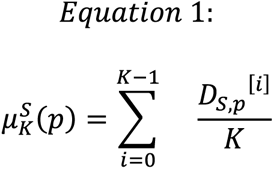

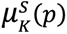 is the mean minimal distance, where K is the number of nearest neighbours

Increasing the KNN parameter increases robustness by ignoring outlier annotation points but can also hamper spatial resolution. However, both the spot grid diameter r and K should remain constant across the study for a given annotation, preventing inconsistencies of distances and axis values between samples. Next, to construct an axis anchored to the interface between two structures, we derived Equation 2, which fits a normalised sigmoidal curve to the relative distances of each point μ(p) and places point p in a relative position in respect to these structures:

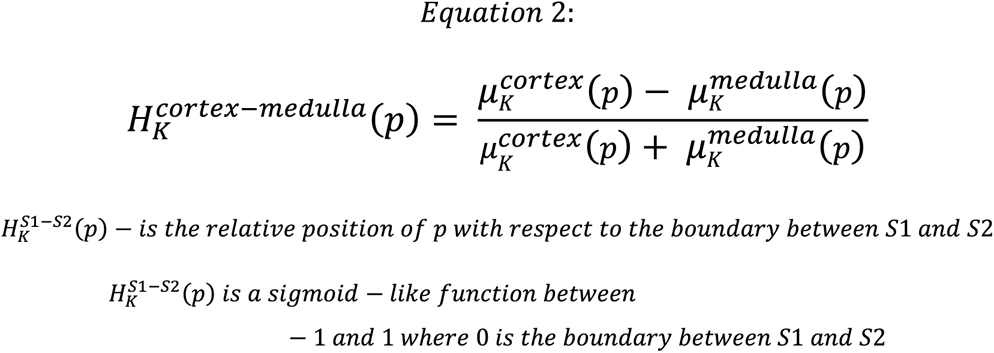

Finally, because the axis aims to span three major thymic compartments (capsule, cortex, and medulla) we linearly combined the two axis functions (H) as detailed in Equation 3:

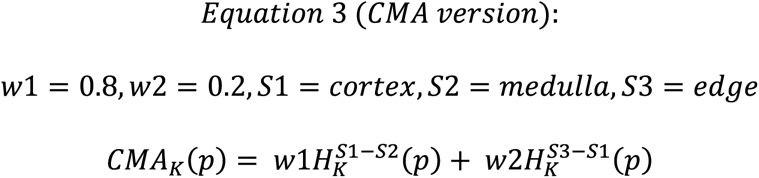

Of note, we also derived a general form of the axis as it could potentially account for more than three histological structures as depicted in Equation 4:

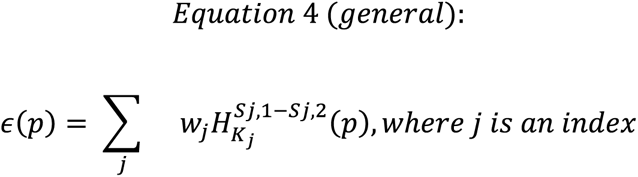

To derive the two-structure axis from Equation 2, users can apply imagespot.calculate_axis_2p(); for the three-point axis in Equation 3 use imagespot.calculate_axis_3p(). All axis related functions are part of the imagespot toolbox pipeline: https://github.com/Teichlab/thymus_spatial_atlas/blob/main/ImageSpot/ To associate defined morphological compartments with the continuous axis and provide a common reference, while facilitating improved visualisation and interpretation, we binned the axis to discrete annotation layers according to the following rules, derived by imagespot.bin_axis().

| Annotations | Assignment rule |
| --- | --- |
| Capsular | $(-0.75 \leq \text{cma})$ and manually annotated from “edge” |
| Subcapsular | $(-0.75 \leq \text{cma}) \ \& \ (-0.62 > \text{cma})$ |
| Cortical level I | $(-0.62 \leq \text{cma}) \ \& \ (-0.55 > \text{cma})$ |
| Cortical level II | $(-0.55 \leq \text{cma}) \ \& \ (-0.45 > \text{cma})$ |
| Cortical level III | $(-0.45 \leq \text{cma}) \ \& \ (-0.3 > \text{cma})$ |
| Cortical CMJ | $(-0.3 \leq \text{cma}) \ \& \ (0.1 > \text{cma})$ |
| Medullar CMJ | $(0.1 \leq \text{cma}) \ \& \ (0.55 > \text{cma})$ |
| Medullar level I | $(0.55 \leq \text{cma}) \ \& \ (0.65 > \text{cma})$ |
| Medullar level II | $(0.65 \leq \text{cma}) \ \& \ (0.7 > \text{cma})$ |
| Medullar level III | $(0.7 \leq \text{cma})$ |

## 10x Visium spatial transcriptomics sample processing and sequencing

Fresh thymus tissue was first transferred to PBS and then placed on ice for a few minutes to clear away any excess media and preservation liquid (e.g. hypothermosol). Next, as much liquid as possible was removed from the sample and, if necessary, the tissue was trimmed to fit into a cryomold. The sample was placed in a cryomold (Tissue-Tek AGG4581) filled with OCT (Leica biosystems 14020108926) and positioned according to the desired orientation using thin forceps. The cryomold was then placed in isopentane that had been equilibrated to -60°C and left to fully freeze for 2 min. The sample was then rested on dry ice to allow draining of the isopentane. Finally, the cryomolds were wrapped in foil and stored at -80°C. On the day of sectioning, samples were removed 1h before sectioning and placed in the cryostat (Leica biosystems) at -18°C to equilibrate. Tissue was sectioned at 7-10 μm thickness and sections were placed on Visium slides according to the manufacturer’s protocol. Sections were stained with Haematoxylin and Eosin and imaged at 20X magnification (hamamatsu Nanozoomer 2.0 HT). Libraries were further processed according to the manufacturer’s protocol (Visium Spatial Gene Expression Slide & Reagent Kit, 10x Genomics, PN-1000184)(see **Supplementary Table 2** for permeabilisation times). Samples were sequenced on a Novaseq6000 sequencer (Illumina). Libraries were mapped with SpaceRanger (10x Genomics, see **Supplementary Table 2** for version).

### VisiuMator pre-processing and image registration

To process the Visium histology image data in higher resolution than the SpaceRanger defaults, which are either 600 (“lowres”) or 2000 (“hires”) pixels along longest image dimension, we built a custom pipeline to extract an additional layer of image resolution at 5000 pixels (“hires5K”), which we found to be more suitable for morphological analysis. We also developed our own fiducial image registration pipeline for increased accuracy with cellpose (https://www.cellpose.org/) and RANSAC affine registration (https://scikit-image.org/) of reference fiducial frame (provided by 10x Genomics). Lastly, for flexible tissue detection we used Otsu thresholding with adjustable threshold (see **Extended Data Figure 2A**). The output of our pre-processing pipeline is compatible with the SpaceRanger output format and was applied to all the Visium sections included in this study. A full example of the Visium processing pipeline and annotation is provided on github. https://github.com/Teichlab/thymus_spatial_atlas/blob/main/ImageSpot/visiumator/

### TissueTag Visium annotation

All Visium samples were annotated with TissueTag by semi-automatic mode to consistently call the border between cortex and medulla. Cortex and medulla pixels were predicted with a random forest pixel classifier (sklearn.ensemble.RandomForestClassifier) by generating training annotations based on spots that showed the highest gene expression of *AIRE* (for medulla) and *ARPP21* (for cortex). These initial annotations were corrected with the TissueTag manual annotation tool “annotator” and the “capsule/edge” annotations were added manually based on H&E. Tissue regions that showed substantial freezing or sectioning artefacts were also labelled manually. The annotation categories for “annotation_level_0” were cortex, medulla, edge and artefact, which were later used for calculations of the cortico-medullary axis. In addition, finer tissue structures such as Hassall’s Corpuscle (HC), Perivascular space (PVS), and adipocytes in paediatric samples or lymph aggregates in fetal samples were manually delineated and stored as “annotation_level_1”. Finally, we used TissueTag’s “poly_annotator” to mark individual lobules, which were saved as “annotation_lobules_0”. All annotations were saved as TissueTag output format, which logs the annotation resolution, the pixels per μm (ppm) as well as pixel value interpretation of annotation names (e.g., 1 = “Medulla”) and colours (e.g., “Medulla”: “magenta”). To robustly and efficiently migrate TissueTag annotations to the Visium objects, we first transferred TissueTag annotations from pixel space to a high-resolution hexagonal grid space (15 μm spot diameter and 15 μm point-to-point centre distance with no gap between spots) based on the median pixel value of the centre of the spot (radius/4) in the annotated image. Next, to generate continuous annotations for Visium data we measured for each spot in the hexagonal high-resolution grid the mean Euclidean distance to the 10 nearest points from each annotated structure in annotation_level_0 as well as the distance from the closest point for structures in annotation_level_1. All annotations were mapped to the Visium spots by proximity of the spot annotation grid to the nearest corresponding spot in the Visium array.

### Spatial Visium mapping with cell2location

For cell2location mapping, all Visium objects (keeping fetal and paediatric separate) were concatenated into a single object as described in the cell2location tutorial. The mapping pipeline was executed as described at https://github.com/Teichlab/thymus_spatial_atlas/blob/main/manuscript/ Briefly, we subsetted the data according to either fetal or paediatric stage and removed sparse cell annotations that were predominantly found in one of these stages: fetal (following cells were removed) - to_remove = [’T_CD8_memory’,’T_Treg_CD8’,’fetFB-NKX2-5’,’fetFB-CCL21’,’fetFB- RSPO2’,’T_DP(Q)-HSPH1’,’T_SP8or4’,’T_SP-HSP’,’T_DN(Q)-stress_1’,’T_DN(Q)- stress_2’,’T_DN(Q)-intermediate’,’T_Treg-intermediate’, ‘B-plasma’, ‘large_pre_B’, ‘pDC-Prolif’]. Paed (following cells were removed) - prolif_cells = [’DC1-Prolif’, ‘DC2- Prolif’, ‘pDC-Prolif’, ‘PeriloFb-Prolif’, ‘ProlifPericyte’,’mcTEC-Prolif’, ‘B-Prolif’]. fet_specific = [’pro_B’, ‘late_pro_B’, ‘large_pre_B’, ‘small_pre_B’, ‘CMP’, ‘GMP’, ‘InterloFb-COL9A3’, ‘fetFB-NKX2-5’, ‘fetFB-CCL21’,’fetFB-RSPO2’, ‘Mesothelium’, ‘mcTEC-Prolif’, ‘T_NK_fetal’,]. unclear = [’T_DP(Q)-HSPH1’,’T_SP8or4’,’T_SP- HSP’,’T_DN(Q)-stress_1’,’T_DN(Q)-stress_2’,’T_DN(Q)-intermediate’,’T_Treg-intermediate’].

We further subsetted the cells by limiting the maximum number of cells from an annotation adata_ref.obs[’cell_type_level_4’].value_counts().loc[adata_ref.obs[’cell_type_level_ 4’].value_counts()<=40], preventing over-representation of abundant cell types in the single-cell data. We then removed cell cycle (**Supplementary Table 8**), mitochondrial (adata.var_names.str.startswith(’MT-’))), and TCR genes (re.search(’^A^TR[AB][VDJ]|^A^IG[HKL][VDJC]’). We used the following parameters to determine the highly variable genes: sc.pp.highly_variable_genes(adata_ref, n_top_genes=18000,subset=True,layer=’counts’,flavor=“seurat_v3”, batch_key=’sample’, span=1) and metadata information to correct for batch effects in the cell2location model - cell2location.models.RegressionModel.setup_anndata(adata=adata_ref, batch_key=‘sample’, labels_key=‘annotation_level_3’, categorical_covariate_keys=[’chemistry_simple’,’age_group’,’study’,’donor’], continuous_covariate_keys=[’age_numeric’]). We ran the cell2location pipeline on a computing cluster with NVIDIA A100 GPUs and 80GB GPU RAM.

### Quality control and batch correction of Visium data

All Visium data after cell2location deconvolution were subjected to filtering based on read coverage and predicted cell abundance. Spots with fewer than 800 genes per spot or fewer than 30 predicted cells were omitted. In addition, annotated tissue artefacts and areas not assigned to a specific structure were removed. Next, in order to batch correct the samples we performed scVI integration after removing cell cycle, mitochondrial and TCR genes from the highly variable gene selection: sc.pp.highly_variable_genes(adata_tmp,n_top_genes=4000,subset=True,flavor=“seurat_v3”, batch_key=’SampleID’,span=1). Training was performed with the following parameters:

scvi.model.SCVI.setup_anndata(adata_tmp,batch_key=’SampleID’,categorical_covar iate_keys=[’SlideID’,’Spaceranger’,’section_thickness(um)’],continuous_covariate_ke ys=[’Age(numeric)’,’n_genes_by_counts’]). vae = scvi.model.SCVI(adata_tmp,n_layers=2, n_latent=30), vae.train(max_epochs=300, batch_size=2000).

Before performing any association analysis with the CMA, we removed lobule data (based on “annotations_lobules_0”) that had fewer than 3 medullar or cortical spots, since we expected our CMA model to be less accurate in cases without cortex or medulla.

### PCA cumulative contribution of CMA

To estimate the dependency of the axis on spot gene variance across samples, we first performed log normalisation and combat regression to regress out the batch effect of individual samples on gene counts as follows:sc.pp.normalize_total(adata_paed_cma,target_sum=2500), sc.pp.log1p(adata_paed_cma), sc.pp.combat(adata_paed_cma, key=’SampleID’, inplace=True). We then computed Spearman correlation coefficients between the CMA values and the n_gene_by_counts in the transcriptomes of the Visium data sets. This technical measure, in our hands, was mostly influenced by inconsistent permeabilisation during Visium library preparation and constituted as the largest technical source of variance we found in both fetal and paediatric Visium samples. To estimate the cumulative contribution of the CMA amd n_gene_by_counts among the 9 first PCs we calculated the absolute value of Spearman’s R with the cumulative explained variance of the first PCs with either the CMA or n_gene_by_counts.

### Cytokine clustering by expression along the binned axis in Visium data

To analyse cytokine gradients based on spatial distribution across CMA bins, we first selected a group of 65 cytokines from the CellphoneDB database (v4.1.0, bat categories, [’Cytokine’, ‘Growthfactor I Cytokine’, ‘cytokine’, ‘Cytokine | Hormone’]). We excluded cytokines that were expressed in less than 5% of the spots in all CMA bins. We then performed hierarchical clustering of the standardised mean expression of genes across bins using the Ward linkage method with the “linkage” function from the scipy.cluster.hierarchy module. A heat map was generated using the “matrixplot” function from the scanpy package^97^.

### Cell and gene weighted location on Visium axis

To estimate the average position of a cell or gene distribution along either CMA and HC axes (L2 distance to the nearest HC), spots with low gene expression were filtered out by using appropriate thresholds (0.2 for scVI corrected gene expression and 0.5 for predicted cell abundances). Then, the position of a gene or cell was calculated according to the following formula: For every gene/cell for an array of “spot_counts” and spot axis positions (“axis_value”), the weighted mean (gene/cell) = (numpy.dot(spot_counts,axis_value)/numpy.sum(spot_counts)).

### IBEX clinical cohort details and sample preparation

Human thymus samples were obtained from the pathology department of the Children’s National Medical Center in Washington, DC following cardiothoracic surgery from children with congenital heart disease, as the thymic tissue is routinely removed and discarded to gain adequate exposure of the retrosternal operative field. Use of these thymus samples for this study was determined to be exempt from review by the NIH Institutional Review Board in accordance with the guidelines issued by the Office of Human Research Protections. There were no genetic concerns for the patients in this cohort. Details about the cohort can be found in **Supplementary Table 2**. Human thymuses were placed in PBS upon receipt and processed within 24 h post-surgery. Excess fat and connective tissue were trimmed and sectioned into < 5 mm cubes. For IBEX imaging, human thymi were fixed with BD CytoFix/CytoPerm (BD Biosciences) diluted in PBS (1:4) for 2 days. Following fixation, all tissues were washed briefly (5 min per wash) in PBS and incubated in 30% sucrose for 2 days before embedding in OCT compound (Tissue-Tek) as described previously^26,32^.

### IBEX sample imaging preparation

IBEX imaging was performed on fixed frozen sections as described previously^26,32^. Briefly, 20 μm sections were cut on a CM1950 cryostat (Leica) and adhered to 2-well chambered cover glasses (Lab-tek) coated with 15 μl of chrome alum gelatin (Newcomer Supply) per well. Frozen sections were permeabilised, blocked, and stained in PBS containing 0.3% Triton X-100 (Sigma-Aldrich), 1% bovine serum albumin (Sigma-Aldrich), and 1% human Fc block (BD Biosciences). Immunolabeling was performed with the PELCO BioWave Pro 36500-230 microwave equipped with a PELCO SteadyTemp Pro 50062 Thermoelectric Recirculating Chiller (Ted Pella) using a 2-1-2-1-2-1-2-1-2 program. A complete list of antibodies and an IBEX thymus antibody panel can be found in **Supplementary Table 3**. Custom antibodies were purchased from BioLegend or conjugated in house using labelling kits from (# 1230) for Lumican and Thermo Fisher Scientific (# A20182) for LYVE-1. A biotin avidin kit (Abcam, # ab64212) was used to block endogenous avidin, biotin and biotin-binding proteins prior to streptavidin application. Cell nuclei were visualised with Hoechst 33342 (Biotium) and sections were mounted using Fluoromount G (Southern Biotech). Mounting media was thoroughly removed by washing with PBS after image acquisition and before chemical bleaching of fluorophores. After each staining and imaging cycle, samples were treated with two 15-min treatments of 1 mg/mL of LiBH4 (STREM Chemicals) prepared in diH2O to bleach all fluorophores except Hoechst.

### IBEX image acquisition and alignment

Representative sections from different tissues were acquired using an inverted Leica TCS SP8 X confocal microscope with a 40X objectives (NA 1.3), 4 HyD and 1 PMT detectors, a white light laser that produces a continuous spectral output between 470 and 670 nm as well as 405, 685, and 730 nm lasers. Panels consisted of antibodies conjugated to the following fluorophores and dyes: Hoechst, Alexa Fluor (AF)488, FITC, AF532, phycoerythrin (PE), eF570, AF555, iFluor (iF)594, AF647, eF660, AF680, and AF700. All images were captured at an 8-bit depth, with a line average of 3, and 1024×1024 format with the following pixel dimensions: x (0.284 μm), y (0.284 μm), and z (1 μm). Images were tiled and merged using the LAS X Navigator software (LAS X 3.5.7.23225). For IBEX tissue imaging, multiple tissue sections were examined before selecting a representative tissue section that contained several distinct lobules with multiple functional units, often resulting in an unusually shaped region of interest. Fluorophore emission was collected on separate detectors with sequential laser excitation of compatible fluorophores (3-4 per sequential) used to minimise spectral spillover. The Channel Dye Separation module within the LAS X 3.5.7.23225 (Leica) was then used to correct for any residual spillover. For publication quality images, gaussian filters, brightness/contrast adjustments, and channel masks were applied uniformly to all images. Image alignment of all IBEX panels was performed as described previously using SimpleITK^98,99^. Additional details on antibodies, protocols, and software can be found on the IBEX Knowledge-Base (10.5281/zenodo.7693279).

### IBEXtractor analysis pipeline

To overcome the challenges related to the high cellular density of thymic tissue and accurately capture single cell data from cells of diverse shapes and sizes, we took two distinct approaches for IBEX image analysis: 3D single nuclei segmentation and segmentation-free grid analysis to measure protein expression. To enable comparison to Visium, we set the spot size of the grid to 50 μm in diameter (Extended Data Figure 3). Samples were converted from .ims (IMARIS Oxford Instruments) to either 2D single-plane images or 3D stacks (TIFF) by individual channels with FIJI (bio-formats, https://github.com/Teichlab/thymus_s-patial_atlas/tree/main/ImageSpot/ibextractor).

## 3D#single nuclei segmentation with cellpose

We developed a custom pipeline for 3D single nuclei segmentation with cellpose, focusing on the following steps:

1. Image Preparation: Individual TIFF images were separated by z-plane and channel and the sample channel metadata were extracted as a .csv file. The nuclear staining channel (Hoechst) was located, and a set of tiled 3D image arrays generated.
2. Segmentation: The image arrays were sequentially segmented with cellpose using specific parameters, e.g., diameter, resampling, anisotropy, thresholds, batch size. We used tiles to overcome restrictions on GPU and RAM resources.
3. Re-Stitching and Formatting: After segmentation, tiles were stitched back together, and all segmentation masks were stored in a uint32-bit image format to hold more than 65K mask labels, since a typical sample produces 200-700K cell masks.
4. Handling High Nuclear Density: The high nuclear density, especially in the thymic cortex, produced scenarios where segmentation masks were often bordered by neighbouring cells, resulting in significant inter-cell signal bleed. To overcome this issue, we excluded pixels in the interface between two cells such that the mask boundary was preserved if no neighbouring cell was present.
5. Mask filtering: We removed small cell masks to avoid noise and cell fragments using skimage.morphology.remove_small_objects.
6. Signal Intensity Extraction: For each cell mask post-filtering, we extracted the mean and maximum signal intensity for each channel in the IBEX image. This produced a cellxprotein .csv file for each sample.
7. Data Merging: Once all sample single-cell segmentations were collected, we merged all samples and stored metadata, channel, and spatial information in a unified AnnData object, totalling more than 1.1M cells.

### Segmentation-free 50 μm grid measurements

To capture the channel intensity across the acquisition regions of IBEX samples, we constructed a hexagonal grid of 50 μm diameter and spot-to-spot centre distance, providing the basis of the segmentation-free channel measurement approach. We extracted image intensity per hexagon by maximum z-projection of images for each channel using Fiji. All channel measurements were stored as .csv files and further combined into a unified AnnData object with SpotXProtein measurements.

### TissueTag IBEX annotations

All IBEX samples were annotated with TissueTag manually with the TissueTag manual annotation tool “annotator” based on a virtual H&E image that was generated based on nuclear (Hoechst) and broad epithelial (PanCK) markers by an implementation adapted from^100^. The remaining steps for annotation were identical to the steps described in TissueTag Visium annotation. All annotations were migrated to the IBEX object by proximity of the spot annotation grid to the nearest corresponding spot/cell in the IBEX data See full IBEXtractor example here - https://github.com/Teichlab/thymus_spatial_atlas/blob/main/ImageSpot/ibextractor/

### Label transfer of IBEX cells from scRNA-seq and CITE-seq reference atlas

We utilised a KNN (K-Nearest Neighbours) algorithm to compare the annotated cell types in our scRNA-seq reference atlas with the IBEX single cell segmentations. For this purpose, the protein expression in the IBEX query cells was matched with the RNA expression of the corresponding genes in the scRNA-seq reference. Protein-gene names were converted according to **Supplementary Table 4**. Batch effects were removed from IBEX with scanpy.pp.combat(ibex_gene, key=’sample’, inplace=True). Next, we subsetted each IBEX sample and ran the KNN prediction algorithm per sample with k=30, including the following steps:

1. The shared feature space between the two objects was identified, log-normalised and z-scored on a per-object basis.
2. The K nearest neighbours of the low-dimensional observations (IBEX) in the high-dimensional space (GEX) were identified. The counts of absent features were imputed as the mean of the high-dimensional neighbours.
3. Based on majority voting, the most frequent cell annotation in the GEX reference was assigned to the IBEX query cell. The proportion of KNNs that contributed to the majority voting was recorded as the evidence fraction (KNNf). For example, if out of the 30 nearest neighbours, 13 are labelled as “A”, 10 as “B”, and 7 as “C” in GEX, then the IBEX cell is labelled as “A”. The “fraction” in this case refers to the proportion of the 30 nearest neighbours that contributed to the majority label, which would be KNNf = 13/30 = 0.43 for label “A”.

For visualisation purposes, to generate a symmetrical representation of the axis without subsetting to the axis bins, we further transformed the axis with the following function numpy.exp2(adata.obs[’cma_v2’]-0.16)-1.

The code for the KNN mapping algorithm can be found here - https://github.com/Teichlab/iss patcher/tree/main

The full example for KNN mapping using IBEX can be found here - https://github.com/Teichlab/thymus_spatial_atlas/tree/main/manuscript/figures/figure_4

To annotate T cell types in the IBEX data with high accuracy we applied the KNN algorithm to the CITE-seq T lineage data. We first applied the KNN to IBEX data using the scRNA-seq reference atlas to identify and subsetted IBEX T lineage cells. We then used the CITE-seq data as reference to repeat the KNN-based annotation on these selected cells. This KNN-based reannotation was performed on a hybrid RNA and protein reference, which included protein measurements for the 19 markers assayed in both CITE-seq and IBEX in addition to RNA measurements for the remaining 23 genes as detailed in **Supplementary Table 4**. We used the same KNN implementation as described above but with k=5, while also imputing CD4 and CD8 pseudotime.

### Human thymus FFPE processing

Human paediatric samples were obtained from cardiac corrective surgeries. Removed thymi were directly moved to HypoThermosol(R)(Sigma-adrich H4416-100ML), shipped with a courier with ice packs and processed in under 24 h post surgery. Upon arrival, for FFPE processing, samples were cut to approx 1 cm^A^2 pieces with sharp scissors in 1X DPBS. Tissue pieces were then rinsed in clean DPBS to remove any excess hypothermazol, patted dry with wipes (Kimtech) and placed into 10 % formalin for 16-24 h at room temperature. On the following day tissues were dehydrated and embedded in wax then kept at 4°C.

### RareCyte Immunostaining and single staining 14-plex imaging

Multiplex immunofluorescence and single round imaging was performed as described here^101^. All steps were performed at room temperature unless stated otherwise. Briefly, FFPE blocks were sectioned using a microtome (Leica RM2235) at 3.5-5 μm thickness and placed on a superfrost slide (Fisher scientific 12312148). Slides were dried at 60°C for 60 min to ensure tissue sections had adhered to the slides. After deparaffinisation, tissue sections were subjected to antigen retrieval using the BioGenex EZ-Retriever system (95°C for 5 min followed by 107°C 5 min). To remove autofluorescence, slides were bleached with AF Quench Buffer which consists of 4.5% H2O2 / 24 mM NaOH in PBS. Slides were quenched for 60 min using the HIGH setting with a strong white light exposure followed by further quenching for 30 min using 365 nm HIGH setting using a UV transilluminator. Slides were rinsed with 1X PBS and incubated in 300 μl of Image-iT™ FX Signal Enhancer (Thermo Fisher, # I36933) for 15 min. Slides were rinsed and 300 μl of labelled primary antibody staining cocktail was added to the tissue, which subsequently was incubated for 120 min in the dark within a humidity tray. All antibodies were pre-diluted according to company recommendations and were not adjusted further. Details about antibodies used can be found in **Supplementary Table 5**. Slides were washed with a surfactant wash buffer and 300 μl of nuclear staining in goat diluent was added to the slide. Slides were then incubated in the dark for 30 min in a humidity tray. Slides were then washed and placed in 1X PBS. Finally the slides were coverslipped using ArgoFluor mount media and left in the dark at room temperature overnight to dry. Slides were imaged on the following day using a RareCyte Orion microscope with a 20X objective. Scans were performed using Imager and relevant acquisition settings were applied using the software Artemis. Slides were subsequently transferred to -20°C for extended storage.

### RNAscope processing and imaging

Sections were cut from the fresh frozen OCT-embedded (OCT, Leica) samples at a thickness of 10 μm using a cryostat (Leica CM3050S) and placed onto SuperFrost Plus slides (VWR). Sections were stored at -80 °C until stained. Sections were removed from the -80 °C storage and submerged in chilled (4 °C) 4% PFA for 15 min, then acclimatised to room temperature 4% PFA over 120 min. Sections were then briefly washed in 1X PBS to remove any remaining OCT. Sections were dehydrated in a series of 50%, 70%, 100% and 100% ethanol (5 min each) and air-dried before performing automated 4-plex RNAScope.

Using the automated Leica BOND RX, RNAscope staining was performed on the fresh frozen sections using the RNAscope LS multiplex fluorescent Reagent Kit v2 Assay and RNAscope LS 4-Plex Ancillary Kit for LS Multiplex Fluorescent (Advanced Cell Diagnostics (ACD), bio-techne) as per manufacturer’s instructions. All sections were subjected to 15 min of protease III treatment before staining protocols were performed. Before running RNAscope probe panels, the RNA quality of fresh frozen samples was assessed using multiplex positive (RNAscope LS 2.5 4-plex Positive Control Probe, ACD Biotechne, 321808) and negative (RNAscope® 4-plex LS Multiplex Negative Control Probe, ACD Biotechne, 321838) controls.

The probes were labelled using Opal 520, 570 and 650 fluorophores (Akoya Biosciences, diluted 1:1000) and one probe channel was labelled using Atto 425- Streptavidin fluorophore (Sigma, diluted 1:500), which was first incubated with TSA-biotin (Akoya Biosciences, diluted 1:400). Details on RNAscope probes, procedure, controls and imaging can be found in **Supplementary Table 6**. All nuclei were DAPI-stained (Life Technologies, D1306).

Confocal imaging was performed on a Perkin Elmer Operetta CLS High Content Analysis System using a 20X (NA 0.16, 0.299 μm/pixel) water-immersion objective with 9-11 z-stacks with 2μm-step. Channels: DAPI (excitation 355-385 nm, emission 430-500 nm), Atto 425 (ex. 435-460 nm, em. 470-515 nm), Opal 520 (ex. 460-490 nm, em. 500-550 nm), Opal 570 (ex. 530-560 nm, em. 570-620 nm), Opal 650 (ex. 615-645 nm, em. 655-760 nm). Confocal image stacks were stitched as individual z-stacks using proprietary Acapella scripts provided by Perkin Elmer, and visualised using OMERO Plus (Glencoe Software).

Contrast used for Figure 4G: *DLK2* (magenta 150-500), *IGFBP6* (yellow 200-1500), *LY75* (green 200-4000), *EPCAM* (red 300-2500).

### CITE-seq tissue processing

Paediatric thymus samples from children undergoing cardiac surgery were obtained according to and used with the approval of the Medical Ethical Commission of Ghent University Hospital, Belgium (EC/2019-0826) through the hematopoietic cell biobank (EC-Bio/1-2018). Thymus tissue was cut into small pieces using scalpels and digested with 1.6 mg/ml collagenase (Gibco, 17104-019) in IMDM medium for 30 min at 37°C to generate a single cell suspension. The reaction was quenched with 10% FBS and the thymocyte suspension was passed through a 70 μm strainer to remove undigested tissue. Cells were frozen in FBS containing 10% DMSO and stored in liquid nitrogen until needed.

### CITE-seq antibody preparation

The TotalSeq-C Human Universal Cocktail 1.0 (Biolegend, 399905) was resuspended according to the manufacturer’s instructions. In brief, the lyophilised cocktail was equilibrated to RT for 5 min and then centrifuged at 10.000g for 30s. 27.5 μl Cell Staining Buffer (Biolegend, 420201) was added and the tube was vortexed for 10s, incubated for 5 min at RT, and vortexed again. The resuspended cocktail was centrifuged for 30 s at RT at 10.000g and the entire volume was transferred to a low-bind tube. Finally, the tube was centrifuged again at 14.000g for 10 min at 4°C and 25 μl of the supernatant was used per sample (2×10^A^6 cells in 200 pl).

13 TotalSeq-C antibodies (Biolegend) were titrated individually via flow cytometry using PE-conjugated versions of the same antibody clone as recommended by Biolegend. After choosing a suitable concentration for each antibody, a master mix was prepared for cell staining. For this, all antibodies were initially diluted in the Cell Staining Buffer to obtain a concentration 100-fold higher than the desired final staining concentration. 2 μl of each diluted antibody substock were then combined in a master mix, which was added to the cells for labelling in a total volume of 200 μl. Details on TotalSeq-C antibodies can be found in **Supplementary Table 7**.

### CITE-seq sample preparation

Cells were thawed slowly by gradually adding 15 volumes of pre-warmed IMDM media and pelleted at 1700 rpm for 6 min at 4°C. After resuspending in PBS, cells were passed through a 70 μm strainer to remove clumps. Enrichment for viable cells was achieved using a magnetic bead-based dead cell removal kit (Miltenyi, 130-090-101). For this, cells were pelleted as before, washed with 1X Binding Buffer (part of kit, prepared with sterile distilled water) and resuspended in Dead Cell Removal MicroBeads (part of kit) at a concentration of 10^A^7 total cells/100 μl beads. After incubation at RT for 15 min, cells were applied to an LS column (Miltenyi, 130-122-729), which was pre-rinsed with 3ml 1X Binding Buffer. The column was washed 4x with 3 ml Binding Buffer and the flow through containing viable cells was collected. Cells in the flow through were pelleted and viability was confirmed using trypan blue. 2×10A6 viable cells were used for TotalSeq-C and anti-CD3-PE antibody staining. For this purpose, cells were washed with Cell Staining Buffer (Biolegend, 420201), pelleted at 600 g for 10 min at 4°C, and resuspended in 90 μl Cell Staining Buffer. 10 μl Human TruStain FcX Blocking solution (Biolegend, 422301) was added and cells were incubated for 10 min at 4°C. The TotalSeq-C Human Universal Cocktail 1.0 (Biolegend, 399905) was resuspended as described above, centrifuged at 14.000 g for 10 min at 4°C and 25 μl of the supernatant was added to the blocked cells. Individual TotalSeq-C antibodies were prepared as described above and 26 μl of the master mix was added to each sample. To facilitate enrichment of immature and mature thymocytes via FACS, 10 μl anti-CD3-PE (clone SK7, Biolegend, 344805) was added and samples were topped up with 40 μl Cell Staining Buffer resulting in a total staining volume of 200 μl. Samples were incubated for 30 min at 4°C in the dark. To wash off unbound antibody Cell Staining Buffer was added to the samples, and cells were pelleted for 10 min at 600 g at 4°C. All supernatant was removed, cells were resuspended in Cell Staining Buffer, transferred to a new tube and pelleted as before. Cells were again resuspended in Cell Staining Buffer and pelleted and this wash step was repeated once more before cells were resuspended in 200 μl MACS buffer (PBS + 2% FCS + 2mM EDTA) in preparation for sorting. 1 μl PI was added for detection of dead cells and samples were sorted on a BD FACSAria III or BD FACSAria Fusion cell sorter using a 100 μm nozzle and a maximum flow rate of 4. Cells were gated using forward/side scatter to remove doublets and debris, then dead cells were excluded based on PI staining. CD3- and CD3+ cells were collected separately in cooled IMDM + 50% FCS (Supplementary Figure 6B). Upon completion of the sort, collection tubes were topped up with DPBS and cells were pelleted at 400 g for 10 min at 4°C. Supernatant was removed and cells were resuspended at an estimated concentration of 1500 cells/μl in PBS + 0.04% BSA (Miltenyi, 130-091-376), of which 16.5 μl was used in the GEM generation step. The Next GEM Single Cell 5’ Kit v2 (10x Genomics, 1000265) and the manufacturer’s protocol CG000330 Rev A were used to prepare the reaction master mix, and load cells, gel beads and partitioning oil on a Chip K (10x Genomics, 1000286). GEMs were generated using a Chromium Controller (10x Genomics), transferred to a tube strip and the reverse transcription was carried out in a BioRad C1000 Touch Thermal Cycler according to the protocol. The samples were stored at 4°C overnight and the library preparation was carried out on the following day.

### CITE-seq library preparation and sequencing

Feature barcode (FB), gene expression (GEX) and TCR libraries were prepared according to protocol CG000330 Rev A (10x Genomics) using the Chromium Next GEM Single Cell 5’ GEM Kit v2 (10x Genomics, 1000244), Library Construction Kit (10x Genomics, 1000190), 5’ Feature Barcode Kit (10x Genomics, 1000256), Human TCR Amplification Kit (10x Genomics, 1000252), Dual Index Kit TT set A (10x Genomics, 1000215) and Dual Index Kit TN set A (10x Genomics, 1000250). The protocol version for >6000 cells was followed and libraries were amplified for 13 cycles (cDNA), 14 cycles (GEX), 8 cycles (FB) or 12+10 cycles (TCR libraries). Library quality and quantity were checked on a Bioanalyzer instrument (Agilent) using a High Sensitivity DNA Assay. Libraries were pooled and sequenced on a Novaseq6000 instrument (Illumina) to a minimum of 25.000 reads/cell for GEX, 10.000 reads/cell for FB, and 5000 reads/cell for TCR libraries.

### CITE-seq QC and denoising

Fastq files from FB libraries were mapped with cellranger 7.0.0. GEX libraries were mapped with Starsolo as described above for scRNA-seq data. CITE-seq data was first subjected to quality control processing based on RNA properties as described above. For the retained high-quality cells, ADT data was then denoised using dsb v1.0.3^102^. For this purpose, empty droplets were identified in the unfiltered mapping output and used as reference for estimation of noise and antibody background levels. Approximately 1.4 million droplets were selected based on RNA count < 240 reads, ADT count between 120 and 240 reads, and < 5% mitochondrial reads to ensure that damaged cells were not included in the subset. During denoising and normalisation with “DSBNormalizeProtein”, ADT data from droplets was used for background correction and the 7 isotype control antibodies included in the TotalSeq-C Human Universal Cocktail were used to determine technical variation. Data was scaled based on the background by subtracting the mean and then dividing by the standard deviation of the empty droplets. Negative values after denoising, which correspond to very low expression, were reset to zero to improve interpretation and visualisation.

### CITE-seq annotation

Annotation of the CITE-seq data was carried out on integrated RNA and denoised ADT modalities. For this purpose, both data modalities were log-normalised and the top 2000 highly variable genes (HVGs) were identified, followed by PCA on the scaled HVGs using the standard functions in the Seurat package^103^. MNN correction was applied to the PCA loadings matrix via the “reducedMNN” function from the Batchelor package^104^ to reduce batch effects between samples. To integrate both modalities, multimodal neighbours and modality weights were identified and a weighted neared neighbours (WNN) graph ^103^ was constructed using “FindMultimodalNeighbors” based on the PCA. The number of PCs to be used was determined based on the difference in variation of consecutive PCs being higher than 0.1%. A UMAP visualisation was generated based on the WNN graph to represent the weighted combination of both modalities. In addition, a supervised PCA (sPCA) was carried out on transcriptome data using the “RunSPCA” function of the Seurat package, to obtain a dimensionality reduction for the RNA modality that represents the structure of the WNN graph^103^.

To identify cell types and developmental stages we performed Leiden clustering at a low resolution based on the sPCA using the Seurat functions “FindNeighbors” and “FindClusters”. The obtained clusters were then subsetted and analysed individually starting with the most immature cluster, which was identified based on high expression of the marker CD34. For each subset, normalisation and scaling for RNA and ADT data was repeated as described above and a new WNN UMAP and sPCA were constructed. A combination of Leiden clustering on the sPCA and thresholding of protein levels was used to identify cell types. In addition, to identify proliferating cells, cell cycle scoring was carried out using the G2M and S phase markers supplied in the Seurat package. The “FindMarkers” function (Seurat) and the package singleCellHaystack^105^ were used to identify differentially expressed genes and surface markers in a cluster-based and cluster-independent manner, respectively. Distinction of CD4 and CD8 lineage maturation stages was based on CD45RA, CD45RO and CD1A and identical expression cutoffs were used for both lineages to ensure that subsets would be directly comparable.

### CITE-seq pseudotime analysis

To carry out trajectory inference for αβT lineage cells, CITE-seq data was subsetted to contain DP_pos_sel, DP_4hi8lo, SP_CD4_immature / semi-mature / mature, SP_CD8_immature / semi-mature / mature, CD8αα(l)_immature / mature, CD8αα(ll)_immature / mature, Tregjmmature / mature, Treg_PD1, and Treg_CD8 subsets. A new WNN UMAP was constructed based on surface protein and RNA. Slingshot was used to construct a minimum spanning tree on the WNN UMAP using the “getLineages” function based on mutual nearest neighbour-based distance with DP_pos_sel set as start point and SP_CD4_mature, SP_CD8_mature, SP_Treg_mature, and CD8ααll_mature specified as end points. Smooth lineages were obtained using the “getCurves” function and the derived pseudotime orderings were used to assess transcript and surface marker expression throughout differentiation.

### CITE-seq cell2location mapping, integration and processing

For the focused multimodal analysis we subsetted the AnndData object (HSTA_v17.h5ad) to include only post-natal data. We further subsetted the T cell lineage to cells which had CITE-seq protein information in the new object (HTSA_paed_multi_v17.h5ad). This data set includes 3 modalities: RNA expression (either 5’ or 3’), surface protein expression, and TCR sequencing. Cell2location deconvolutes spots by unique expression profiles from a single-cell reference, based on discrete cell annotations. Here, In order to investigate the continuous spatio-temporal nature of CD4/CD8 cell lineages at increased resolution we performed high-resolution Leiden clustering on the CITE-seq data via scanpy.tl.leiden(gex,resolution=35,key_added=’hyper_leiden’), resulting in 624 cell clusters. These cell clusters were then mapped to Visium data with cell2location as described above and further detailed here To measure the position of each cell cluster across the CMA, we selected Visium spots with the highest cluster label abundance (above percentile 95%) and calculated the weighted mean CMA values for these spots. This values was then assigned this to the cells comprising the associated cluster in the single cell AnnData object. Further details and notebooks can be found here -

> https://github.com/Teichlab/thymus_spatial_atlas/blob/main/manuscript/figures/figure_6/

---

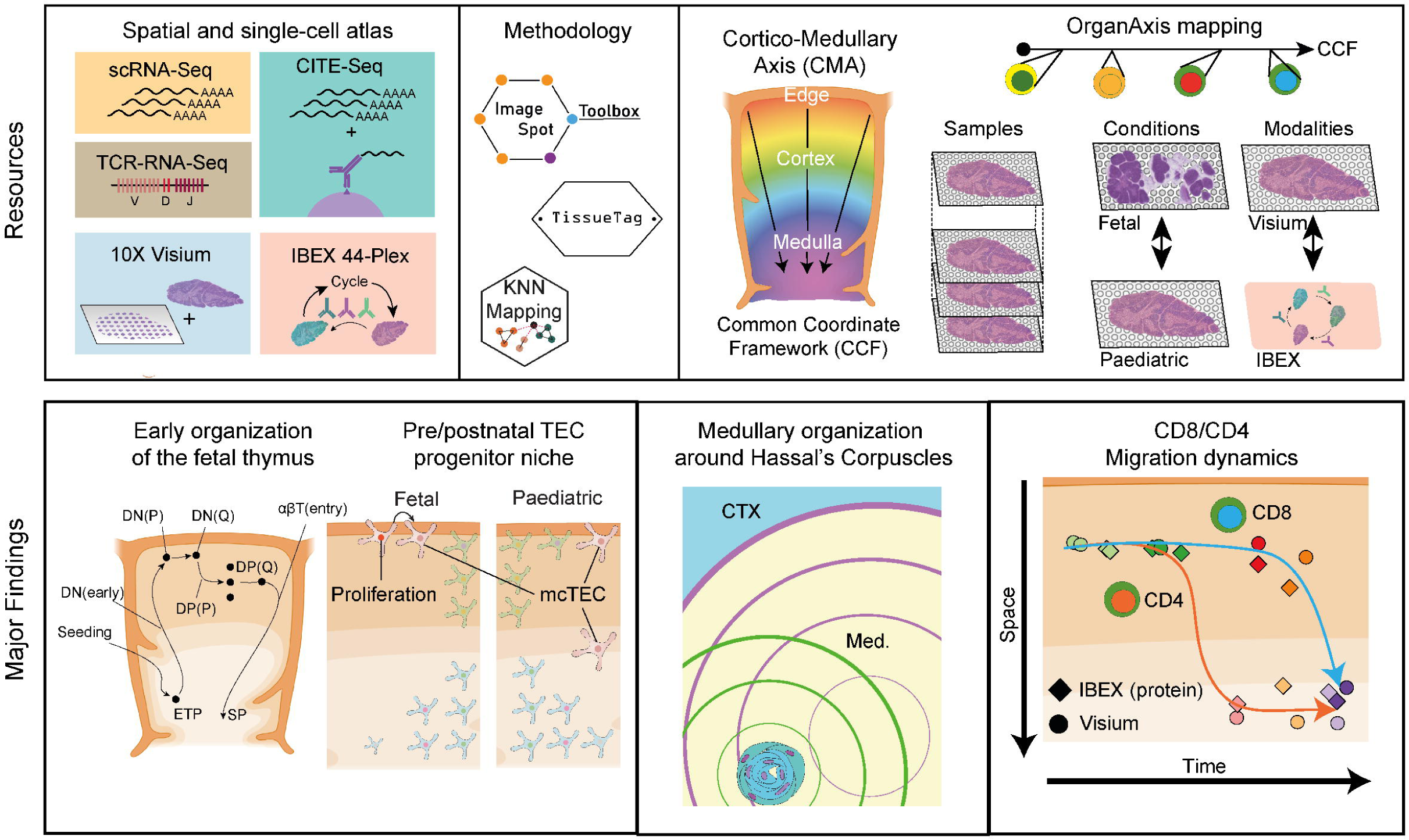

