## Extended data figures for "A spatial human thymus cell atlas mapped to a continuous tissue axis"

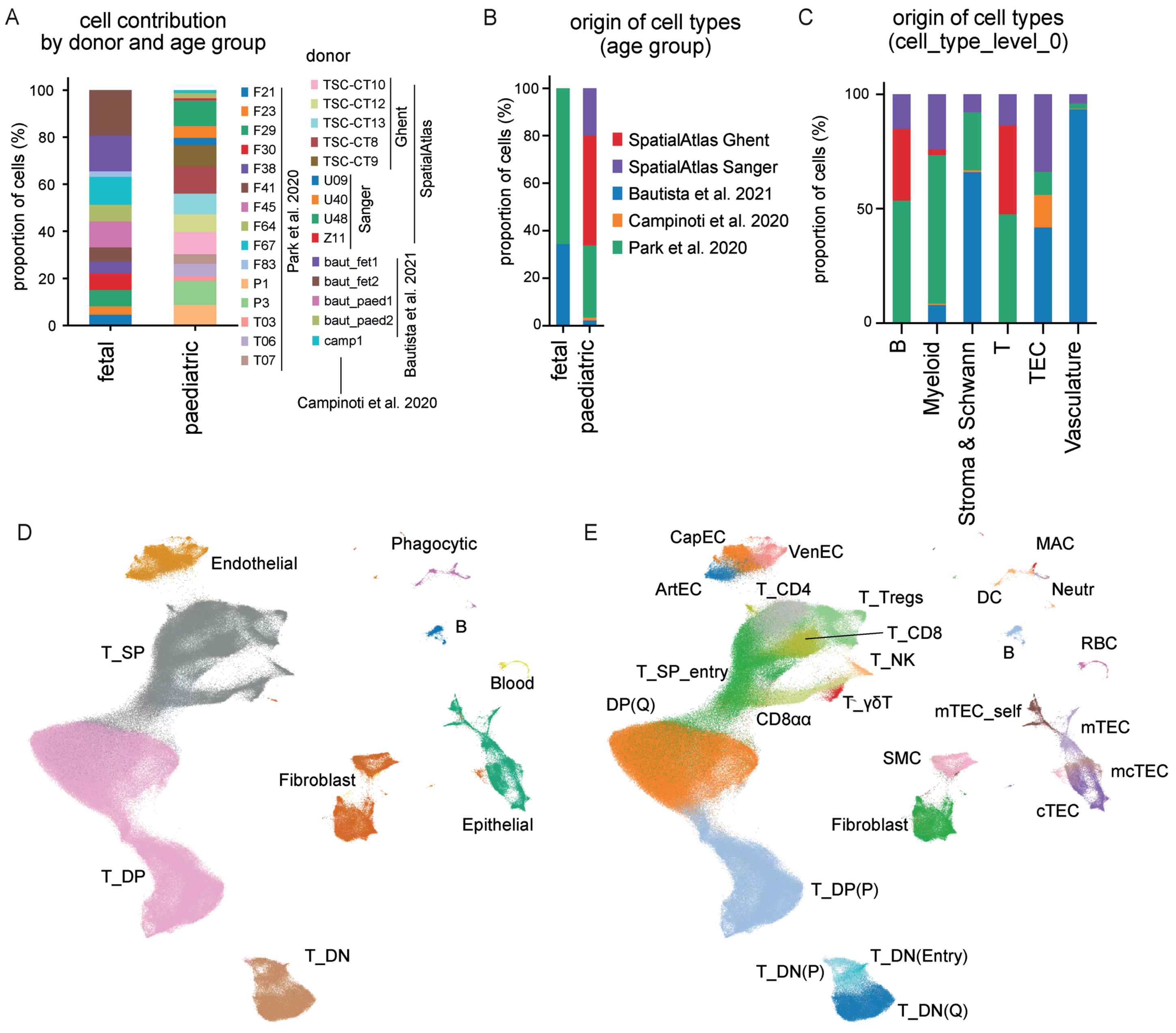
A
cell contribution by donor and age group
proportion of cells (%)
0
20
40
60
80
100
fetal
paediatric
F21
F23
F29
F30
F38
F41
F45
F64
F67
F83
P1
P3
T03
T06
T07
Park et al. 2020
donor
TSC-CT10
TSC-CT12
TSC-CT13
TSC-CT8
TSC-CT9
Ghent
U09
U40
U48
Z11
Sanger
SpatialAtlas
baut_fet1
baut_fet2
baut_paed1
baut_paed2
camp1
Bautista et al. 2021
Campinoti et al. 2020
B
origin of cell types (age group)
proportion of cells (%)
0
20
40
60
80
100
fetal
paediatric
SpatialAtlas Ghent
SpatialAtlas Sanger
Bautista et al. 2021
Campinoti et al. 2020
Park et al. 2020
C
origin of cell types (cell_type_level_0)
proportion of cells (%)
0
50
100
B
Myeloid
Stroma & Schwann
T
TEC
Vasculature
D
Endothelial
Phagocytic
B
T_SP
Blood
Fibroblast
Epithelial
T_DP
T_DN
E
CapEC
VenEC
ArtEC
T_CD4
T_Tregs
T_CD8
T_NK
T_SP_entry
DP(Q)
CD8αα
T_γδT
MAC
DC
Neutr
B
RBC
mTEC_self
mTEC
mcTEC
cTEC
SMC
Fibroblast
T_DP(P)
T_DN(Entry)
T_DN(P)
T_DN(Q)

**Extended Data Figure 1: Composition of fetal and paediatric scRNA-seq data. A.** Relative cell contribution per donor, split by age. Infant was defined as younger than 12 months. Sample origin is indicated for all donors. **B.** Relative contribution of published and newly generated scRNA-seq data sets by donor age. Infant was defined as younger than 12 months. **C.** Relative contribution of published and newly generated scRNA-seq data sets by cell lineage (annotation level 1). **D.** UMAP embedding of the full, integrated scRNA-seq data set with annotations of the major cell lineages (annotation level 2). **E.** UMAP embedding of the full, integrated scRNA-seq data set with more detailed lineage annotations (annotation level 3).

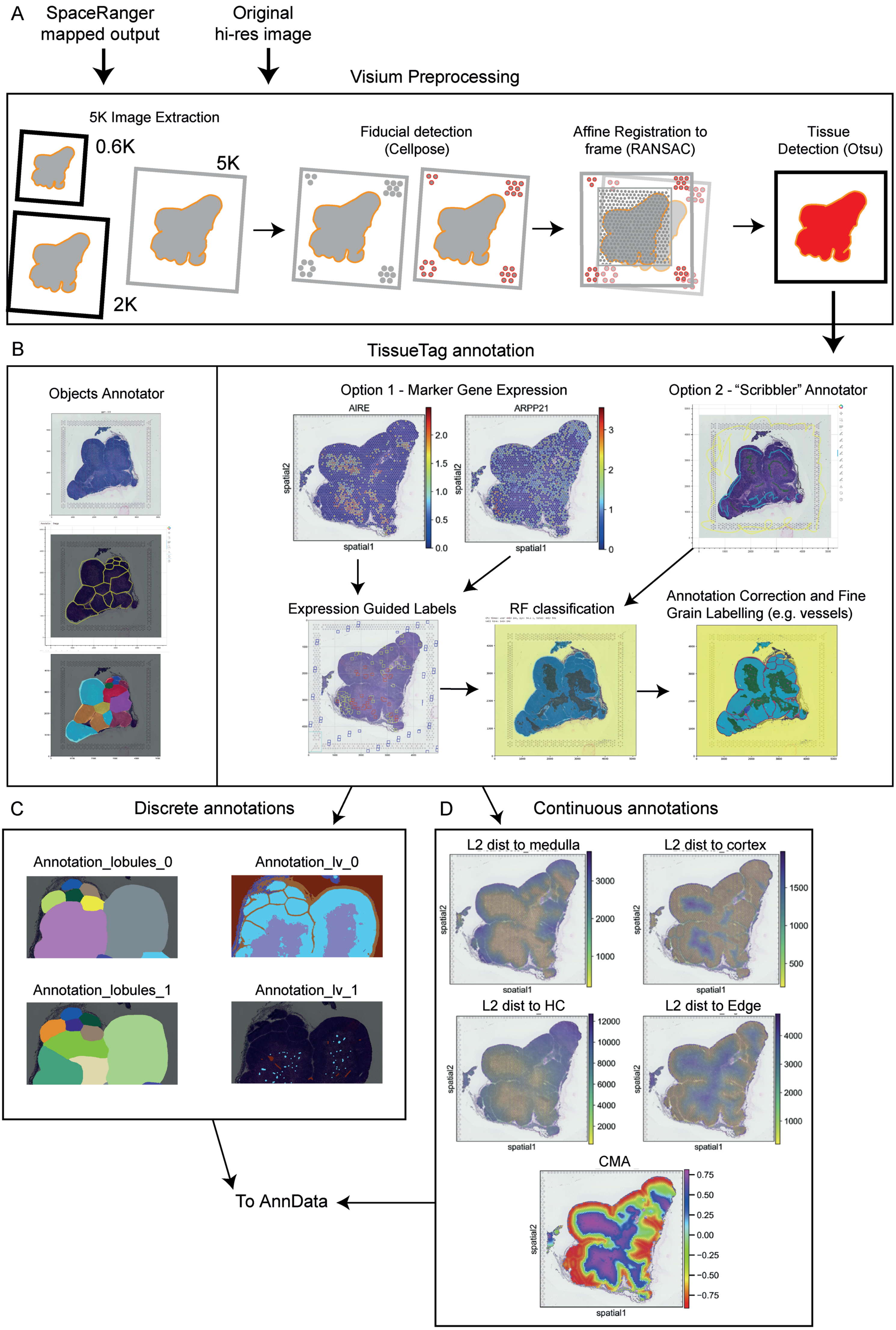
A
SpaceRanger mapped output
Original hi-res image
Visium Preprocessing
5K Image Extraction
0.6K
5K
2K
Fiducial detection (Cellpose)
Affine Registration to frame (RANSAC)
Tissue Detection (Otsu)
B
TissueTag annotation
Objects Annotator
Option 1 - Marker Gene Expression
AIRE
ARPP21
spatial1
spatial2
Option 2 - "Scribbler" Annotator
Expression Guided Labels
RF classification
Annotation Correction and Fine Grain Labelling (e.g. vessels)
C
Discrete annotations
Annotation_lobules_0
Annotation_lv_0
Annotation_lobules_1
Annotation_lv_1
D
Continuous annotations
L2 dist to medulla
L2 dist to cortex
L2 dist to HC
L2 dist to Edge
CMA
To AnnData

**Extended Data Figure 2. General processing steps for Visium data sets. A.** Visium output files from SpaceRanger and the original reference image (optional) are fed into the pre-processing pipeline that first generates a 5k high-resolution reference image, then aligns the image to the fiducial frame (cellpose), and lastly detects the tissue region with an adjustable threshold. **B.** To annotate individual structures, the object annotator tool can count discrete objects, here thymus lobules (left). Schematic of the TissueTag annotation process through either guided RNA marker genes (option 1) or manually scribbling initial tissue annotations (option 2)(right). Initial annotations are then used to guide a random forest classifier and the final step includes manual correction of the predicted annotations. **C.** Discrete tissue annotations produced with TissueTag. **D.** Continuous tissue annotations with TissueTag are used as a reference to construct the cortico-medullary axis (CMA). All annotations are mapped back to the original Visium object for downstream analysis.

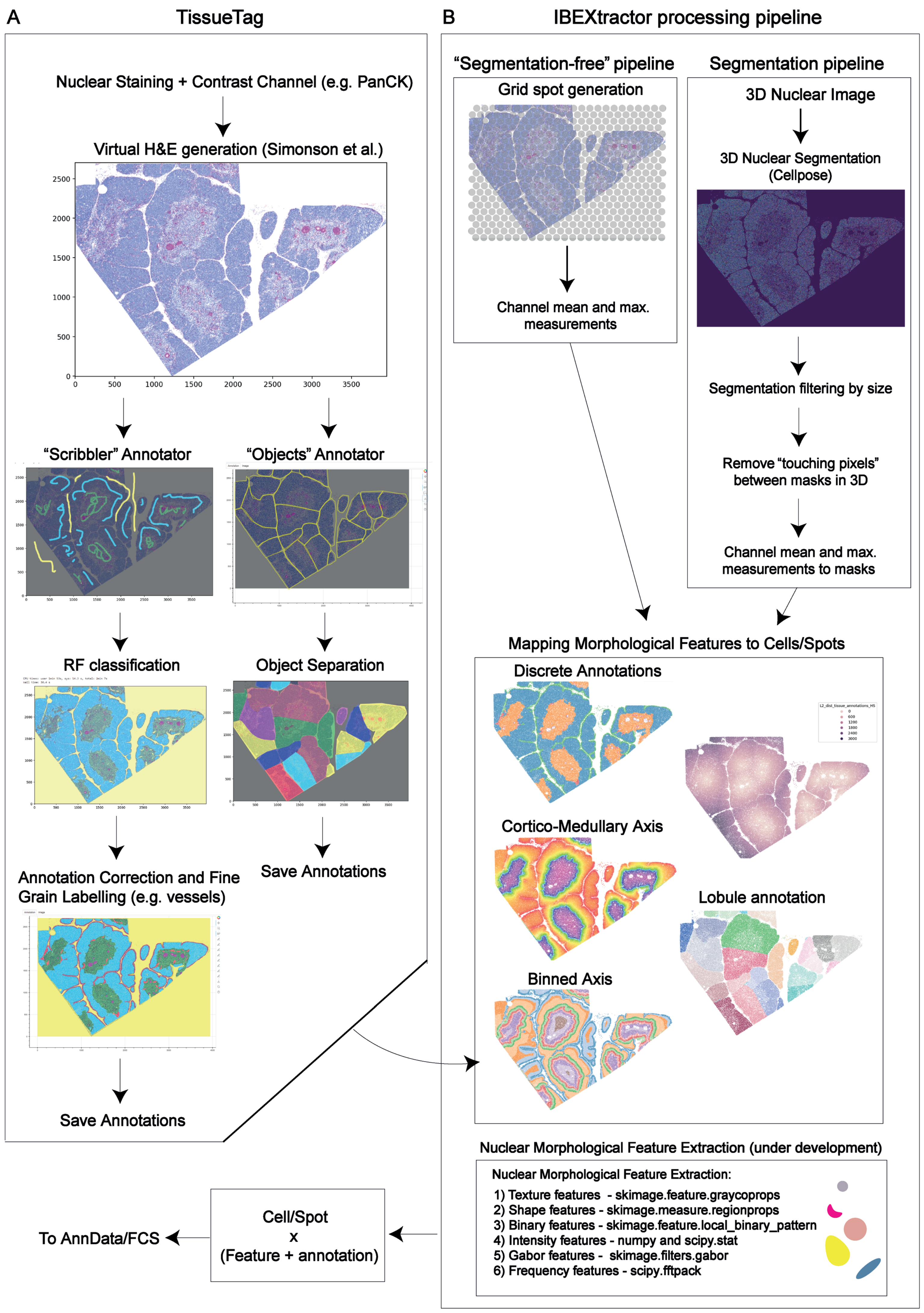
A
TissueTag
Nuclear Staining + Contrast Channel (e.g. PanCK)
Virtual H&E generation (Simonson et al.)
2500
2000
1500
1000
500
0
0
500
1000
1500
2000
2500
3000
3500
"Scribbler" Annotator
"Objects" Annotator
RF classification
Object Separation
Annotation Correction and Fine Grain Labelling (e.g. vessels)
Save Annotations
Save Annotations
B
IBEXtractor processing pipeline
"Segmentation-free" pipeline
Grid spot generation
Channel mean and max. measurements
Segmentation pipeline
3D Nuclear Image
3D Nuclear Segmentation (Cellpose)
Segmentation filtering by size
Remove "touching pixels" between masks in 3D
Channel mean and max. measurements to masks
Mapping Morphological Features to Cells/Spots
Discrete Annotations
L2_dist_tissue_annotations_HS
0
600
1200
1800
2400
3000
Cortico-Medullary Axis
Lobule annotation
Binned Axis
Nuclear Morphological Feature Extraction (under development)
Nuclear Morphological Feature Extraction:
1) Texture features - skimage.feature.graycoprops
2) Shape features - skimage.measure.regionprops
3) Binary features - skimage.feature.local_binary_pattern
4) Intensity features - numpy and scipy.stat
5) Gabor features - skimage.filters.gabor
6) Frequency features - scipy.fftpack
Cell/Spot
x
(Feature + annotation)
To AnnData/FCS

**Extended Data Figure 3. General processing steps for IBEX data sets. A.** Schematic of the TissueTag annotation process. First, a virtual H&E image is generated based on two image channels. Manually scribbled initial tissue annotations are used to guide a random forest classifier and the final step includes manual correction of the predicted annotations (left). To annotate individual structures, the object annotator tool can count discrete objects, here thymus lobules (right). **B.** The IBEXtractor pipeline has two processing options. For the segmentation-free approach (left) a 50 µm grid mask is drawn on the image and used to measure general marker composition and distribution. 3D nuclear segmentations are produced with cellpose and tiled image (right). Then, masks that are too small or large are removed. Pixels that are at the boundary between cells are removed and the mean and maximum values of each cell are recorded. TissueTag annotations are migrated to the image space including discrete and continuous annotations, which serve as the basis to derive the CMA. Nuclear morphology examination is still under development but allows the extraction of distinct morphological features. The IBEXtractor pipeline produces an AnnData spatial object with all meta data and tissue annotations.

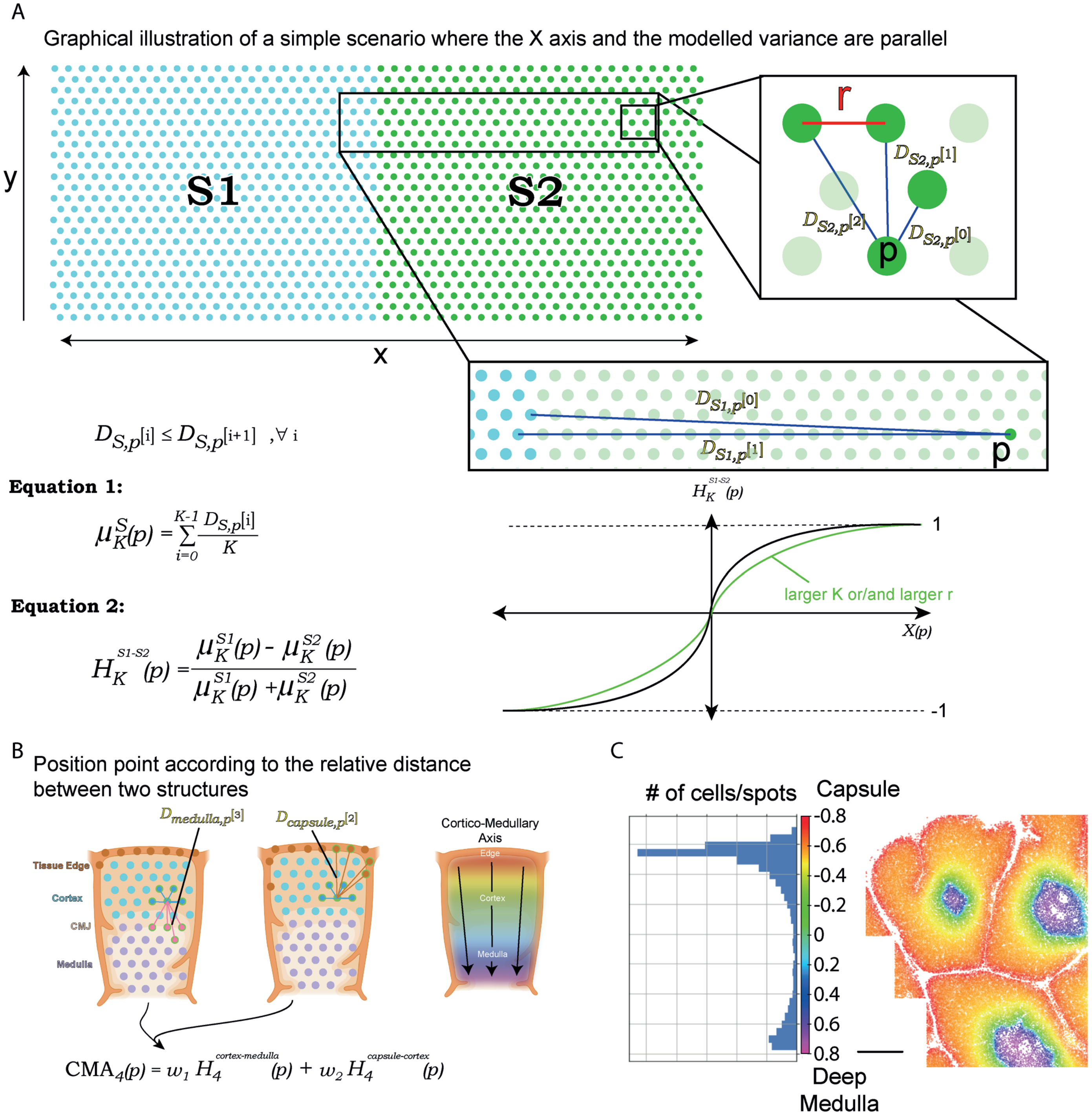
A
Graphical illustration of a simple scenario where the X axis and the modelled variance are parallel
y
x
S1
S2
r
$D_{S2,p}[1]$
$D_{S2,p}[2]$
$D_{S2,p}[0]$
p
$D_{S1,p}[0]$
$D_{S1,p}[1]$
$D_{S,p}[i] \leq D_{S,p}[i+1] \quad ,\forall\, i$
Equation 1:
$\mu_K^S(p) = \sum_{i=0}^{K-1} \frac{D_{S,p}[i]}{K}$
Equation 2:
$H_K^{S1\text{-}S2}(p) = \frac{\mu_K^{S1}(p) - \mu_K^{S2}(p)}{\mu_K^{S1}(p) + \mu_K^{S2}(p)}$
$H_K^{S1\text{-}S2}(p)$
1
-1
larger K or/and larger r
$X(p)$
B
Position point according to the relative distance between two structures
$D_{medulla,p}[3]$
$D_{capsule,p}[2]$
Tissue Edge
Cortex
CMJ
Medulla
Cortico-Medullary Axis
Edge
Cortex
Medulla
$\mathrm{CMA}_4(p) = w_1 H_4^{cortex\text{-}medulla}(p) + w_2 H_4^{capsule\text{-}cortex}(p)$
C
### of cells/spots
Capsule
-0.8
-0.6
-0.4
-0.2
0
0.2
0.4
0.6
0.8
Deep Medulla

**Extended Data Figure 4. Axis graphical intuition. A.** A simplified illustration of how the axis is calculated and "behaves" under the assumption that we are looking to position a cell along the x-axis. We start by measuring the minimal Euclidean distance *D* from k points from each structure S (Equation 1). H is the basic axis function and produces a normalised sigmoid in space where the slope is determined by the grid resolution and number of KNNs to consider for the calculation of equation 1. **B.** The combination of two *H* functions generates the full CMA. **C.** Real life representation and relative cell density along the CMA of an IBEX sample. Notice that the representation of space is far from uniform where regions around the junction get a larger range as opposed to the section in the depth of the cortex that gets a more limited range.

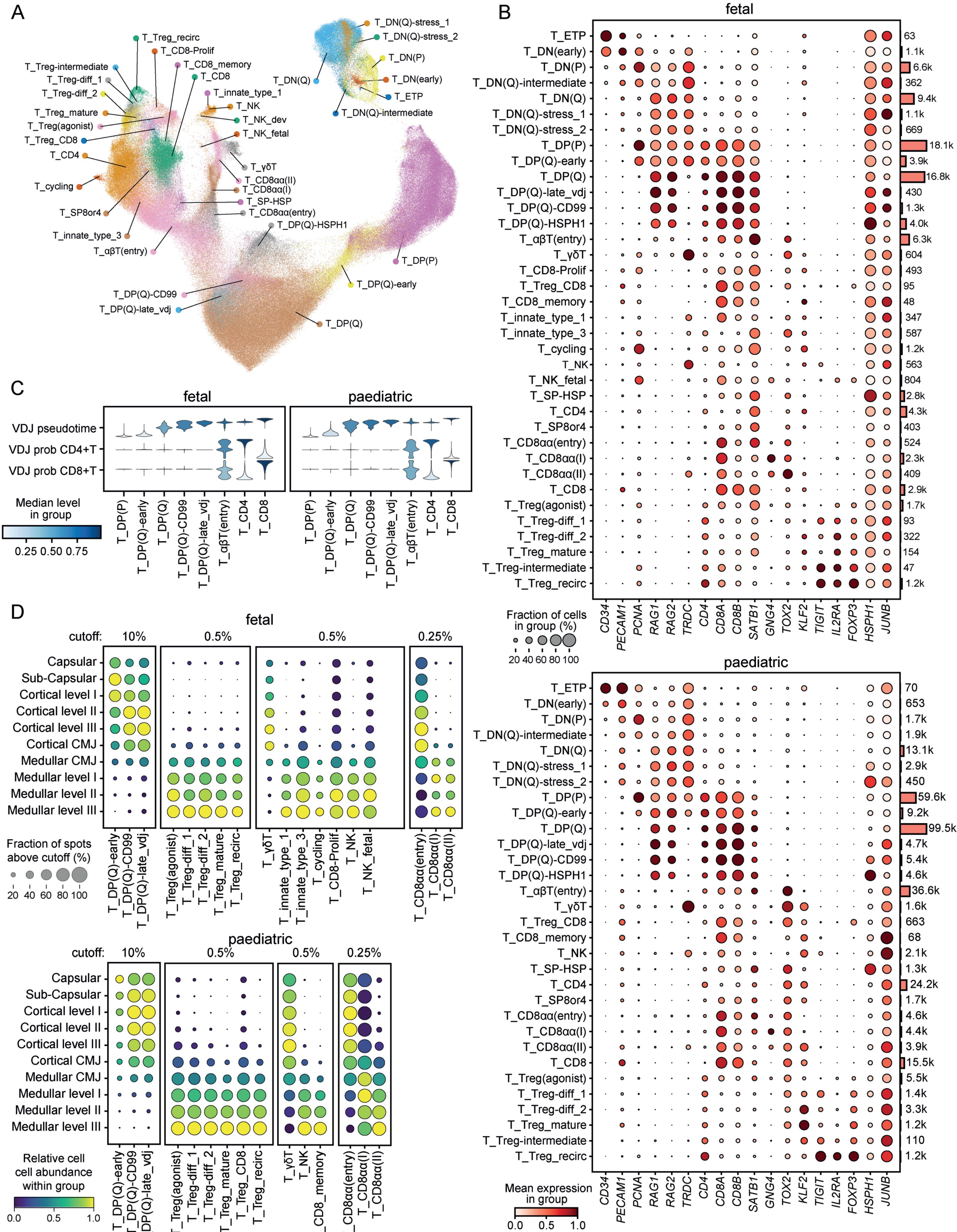
A
T_Treg_recirc
T_CD8-Prolif
T_Treg-intermediate
T_CD8_memory
T_Treg-diff_1
T_CD8
T_Treg-diff_2
T_innate_type_1
T_Treg_mature
T_NK
T_Treg(agonist)
T_NK_dev
T_Treg_CD8
T_NK_fetal
T_CD4
T_γδT
T_cycling
T_CD8αα(II)
T_CD8αα(I)
T_SP-HSP
T_SP8or4
T_CD8αα(entry)
T_innate_type_3
T_DP(Q)-HSPH1
T_αβT(entry)
T_DP(P)
T_DP(Q)-early
T_DP(Q)-CD99
T_DP(Q)-late_vdj
T_DP(Q)
T_DN(Q)-stress_1
T_DN(Q)-stress_2
T_DN(P)
T_DN(Q)
T_DN(early)
T_ETP
T_DN(Q)-intermediate
B
fetal
T_ETP 63
T_DN(early) 1.1k
T_DN(P) 6.6k
T_DN(Q)-intermediate 362
T_DN(Q) 9.4k
T_DN(Q)-stress_1 1.1k
T_DN(Q)-stress_2 669
T_DP(P) 18.1k
T_DP(Q)-early 3.9k
T_DP(Q) 16.8k
T_DP(Q)-late_vdj 430
T_DP(Q)-CD99 1.3k
T_DP(Q)-HSPH1 4.0k
T_αβT(entry) 6.3k
T_γδT 604
T_CD8-Prolif 493
T_Treg_CD8 95
T_CD8_memory 48
T_innate_type_1 347
T_innate_type_3 587
T_cycling 1.2k
T_NK 563
T_NK_fetal 804
T_SP-HSP 2.8k
T_CD4 4.3k
T_SP8or4 403
T_CD8αα(entry) 524
T_CD8αα(I) 2.3k
T_CD8αα(II) 409
T_CD8 2.9k
T_Treg(agonist) 1.7k
T_Treg-diff_1 93
T_Treg-diff_2 322
T_Treg_mature 154
T_Treg-intermediate 47
T_Treg_recirc 1.2k
CD34 PECAM1 PCNA RAG1 RAG2 TRDC CD4 CD8A CD8B SATB1 GNG4 TOX2 KLF2 TIGIT IL2RA FOXP3 HSPH1 JUNB
Fraction of cells in group (%)
20 40 60 80 100
paediatric
T_ETP 70
T_DN(early) 653
T_DN(P) 1.7k
T_DN(Q)-intermediate 1.9k
T_DN(Q) 13.1k
T_DN(Q)-stress_1 2.9k
T_DN(Q)-stress_2 450
T_DP(P) 59.6k
T_DP(Q)-early 9.2k
T_DP(Q) 99.5k
T_DP(Q)-late_vdj 4.7k
T_DP(Q)-CD99 5.4k
T_DP(Q)-HSPH1 4.6k
T_αβT(entry) 36.6k
T_γδT 1.6k
T_Treg_CD8 663
T_CD8_memory 68
T_NK 2.1k
T_SP-HSP 1.3k
T_CD4 24.2k
T_SP8or4 1.7k
T_CD8αα(entry) 4.6k
T_CD8αα(I) 4.4k
T_CD8αα(II) 3.9k
T_CD8 15.5k
T_Treg(agonist) 5.5k
T_Treg-diff_1 1.4k
T_Treg-diff_2 3.3k
T_Treg_mature 1.2k
T_Treg-intermediate 110
T_Treg_recirc 1.2k
CD34 PECAM1 PCNA RAG1 RAG2 TRDC CD4 CD8A CD8B SATB1 GNG4 TOX2 KLF2 TIGIT IL2RA FOXP3 HSPH1 JUNB
Mean expression in group
0.0 0.5 1.0
C
fetal
paediatric
VDJ pseudotime
VDJ prob CD4+T
VDJ prob CD8+T
T_DP(P) T_DP(Q)-early T_DP(Q) T_DP(Q)-CD99 T_DP(Q)-late_vdj T_αβT(entry) T_CD4 T_CD8
Median level in group
0.25 0.50 0.75
D
fetal
cutoff: 10% 0.5% 0.5% 0.25%
Capsular
Sub-Capsular
Cortical level I
Cortical level II
Cortical level III
Cortical CMJ
Medullar CMJ
Medullar level I
Medullar level II
Medullar level III
T_DP(Q)-early T_DP(Q)-CD99 T_DP(Q)-late_vdj
T_Treg(agonist) T_Treg-diff_1 T_Treg-diff_2 T_Treg_mature T_Treg_recirc
T_γδT T_innate_type_1 T_innate_type_3 T_cycling T_CD8-Prolif T_NK T_NK_fetal
T_CD8αα(entry) T_CD8αα(I) T_CD8αα(II)
Fraction of spots above cutoff (%)
20 40 60 80 100
paediatric
cutoff: 10% 0.5% 0.5% 0.25%
T_DP(Q)-early T_DP(Q)-CD99 T_DP(Q)-late_vdj
T_Treg(agonist) T_Treg-diff_1 T_Treg-diff_2 T_Treg_mature T_Treg_CD8 T_Treg_recirc
T_γδT T_NK T_CD8_memory
T_CD8αα(entry) T_CD8αα(I) T_CD8αα(II)
Relative cell abundance within group
0.0 0.5 1.0

**Extended Data Figure 5. T lineage annotations in scRNA-seq data and spatial mapping via deconvolution of Visium datasets. A.** UMAP embedding of integrated fetal and paediatric scRNA-seq data spanning the entire T lineage with detailed annotations of differentiation stages (“cell_type_level_4”). **B.** Dot plot showing expression of marker genes across the entire T lineage for fetal (top) and paediatric data (bottom). Total cell numbers per cell type are indicated by bar graph. **C.** VDJ pseudotime and CD4/8 linear probability as predicted by the Dandelion pipeline for DP and SP subtypes illustrating the annotation of VDJ-late DP(Q) thymocytes. **D.** Spatial mapping of T lineage cells through calculation of the CMA for Visium data and deconvolution based on scRNA-reference shown in (A). Cell subsets shown in Figure 3 are not included here. Cutoff indicates the minimum proportion of the respective cell type in a Visium spot for the spot to be included. Dot size represents the proportion of spots meeting the cutoff and colour indicates the relative cell abundance.

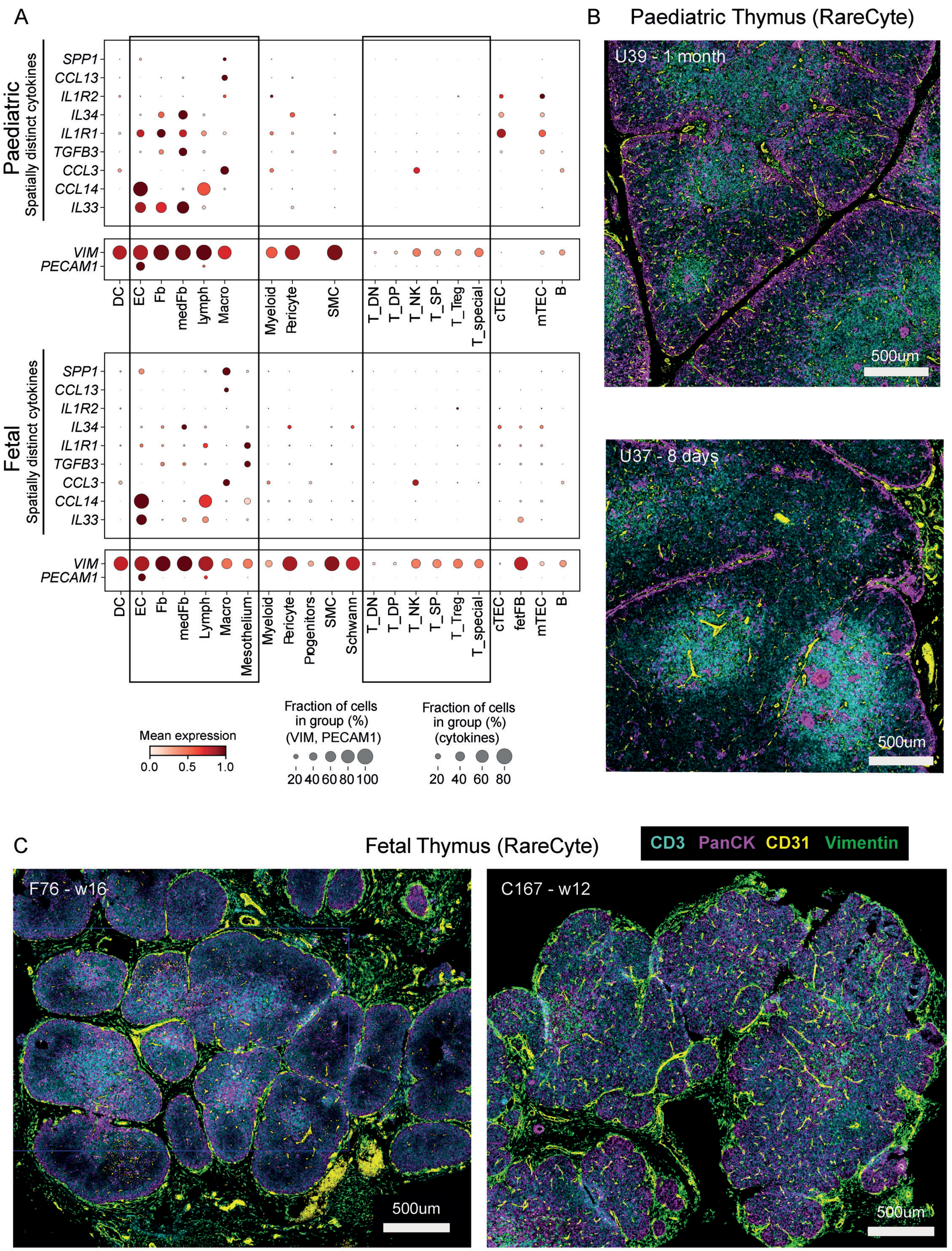
A
B
Paediatric Thymus (RareCyte)
Paediatric
Spatially distinct cytokines
SPP1
CCL13
IL1R2
IL34
IL1R1
TGFB3
CCL3
CCL14
IL33
VIM
PECAM1
DC
EC
Fb
medFb
Lymph
Macro
Myeloid
Pericyte
SMC
T_DN
T_DP
T_NK
T_SP
T_Treg
T_special
cTEC
mTEC
B
Fetal
Mesothelium
Progenitors
Schwann
fetFB
U39 - 1 month
U37 - 8 days
500um
Mean expression
0.0
0.5
1.0
Fraction of cells in group (%) (VIM, PECAM1)
20
40
60
80
100
Fraction of cells in group (%) (cytokines)
20
40
60
80
C
Fetal Thymus (RareCyte)
CD3
PanCK
CD31
Vimentin
F76 - w16
C167 - w12

**Extended Data Figure 6. Stromal variance in fetal and paediatric thymus. A.** Dot plot representation of scRNA-seq data by “cell_type_level_1” annotation showing expression profiles of cytokines, which displayed a high degree of divergence between fetal and paediatric Visium datasets in Figure 3F. Vimentin (*VIM*) and *PECAM1* expression are shown as reference for stromal cell staining in RareCyte imaging in (B) and (C). Boxes highlight stromal and T cell subtypes. **B.** RareCyte staining of 4 markers on paediatric thymus: CD3 (cyan) for thymocytes, PanCK (magenta) for TECs, CD31 (yellow) for endothelial cells and VIM (green) as a broad stromal marker. **C.** RareCyte staining of the same markers as in (B) for fetal thymus.

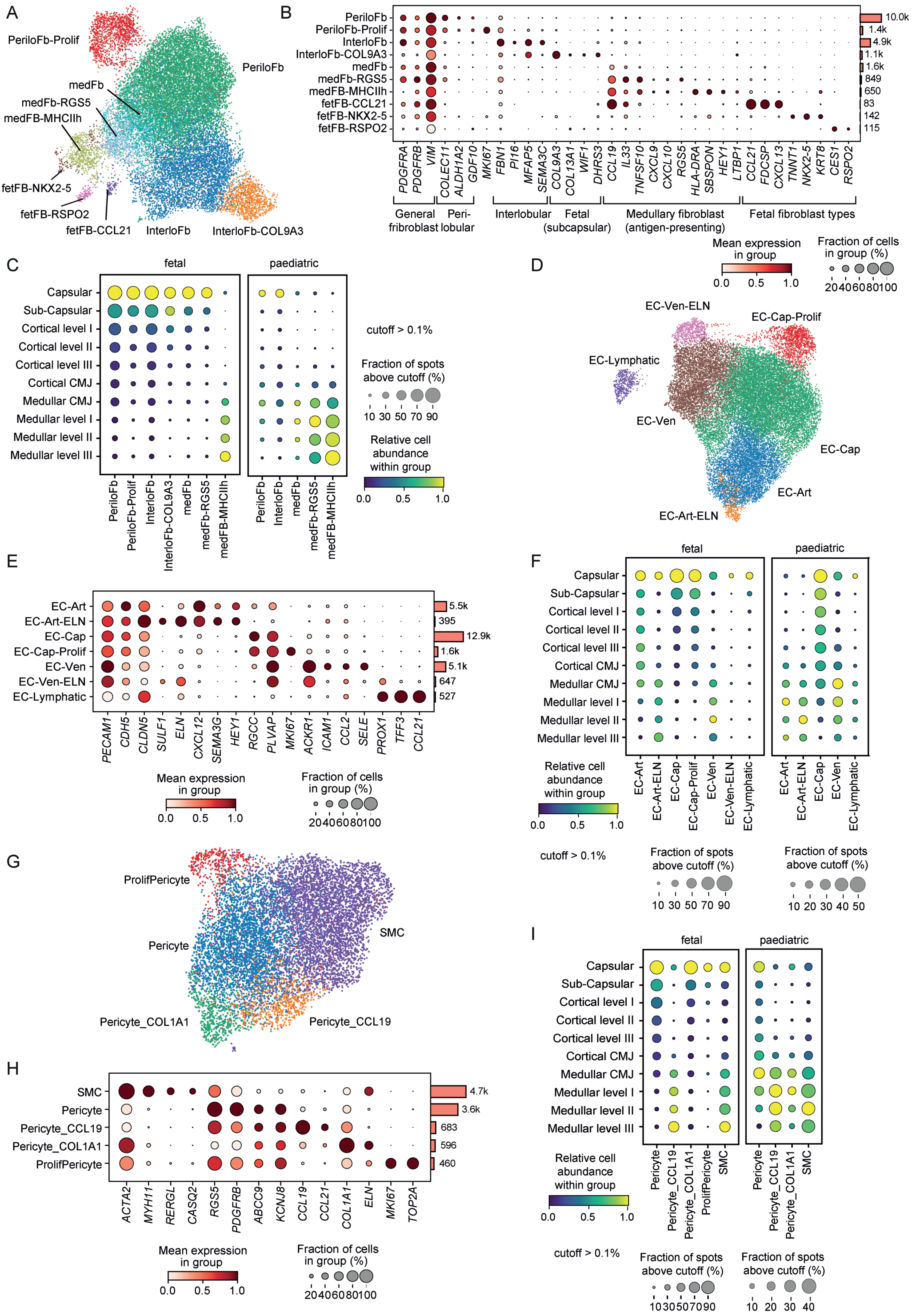
A
PeriloFb-Prolif
PeriloFb
medFb
medFb-RGS5
medFB-MHCIIh
fetFB-NKX2-5
fetFB-RSPO2
fetFB-CCL21
InterloFb
InterloFb-COL9A3
B
PeriloFb
PeriloFb-Prolif
InterloFb
InterloFb-COL9A3
medFb
medFb-RGS5
medFB-MHCIIh
fetFB-CCL21
fetFB-NKX2-5
fetFB-RSPO2
10.0k
1.4k
4.9k
1.1k
1.6k
849
650
83
142
115
PDGFRA PDGFRB VIM COLEC11 ALDH1A2 GDF10 MKI67 FBN1 PI16 MFAP5 SEMA3C COL9A3 COL13A1 WIF1 DHRS3 CCL19 IL33 TNFSF10 CXCL9 CXCL10 RGS5 HLA-DRA SBSPON HEY1 LTBP1 CCL21 FDCSP CXCL13 TNNT1 NKX2-5 KRT8 CES1 RSPO2
General fibroblast
Peri-lobular
Interlobular
Fetal (subcapsular)
Medullary fibroblast (antigen-presenting)
Fetal fibroblast types
Mean expression in group
0.0 0.5 1.0
Fraction of cells in group (%)
20 40 60 80 100
C
fetal
paediatric
Capsular
Sub-Capsular
Cortical level I
Cortical level II
Cortical level III
Cortical CMJ
Medullar CMJ
Medullar level I
Medullar level II
Medullar level III
PeriloFb PeriloFb-Prolif InterloFb InterloFb-COL9A3 medFb medFb-RGS5 medFB-MHCIIh
PeriloFb InterloFb medFb medFb-RGS5 medFB-MHCIIh
cutoff > 0.1%
Fraction of spots above cutoff (%)
10 30 50 70 90
Relative cell abundance within group
0.0 0.5 1.0
D
EC-Ven-ELN
EC-Cap-Prolif
EC-Lymphatic
EC-Ven
EC-Cap
EC-Art
EC-Art-ELN
E
EC-Art
EC-Art-ELN
EC-Cap
EC-Cap-Prolif
EC-Ven
EC-Ven-ELN
EC-Lymphatic
5.5k
395
12.9k
1.6k
5.1k
647
527
PECAM1 CDH5 CLDN5 SULF1 ELN CXCL12 SEMA3G HEY1 RGCC PLVAP MKI67 ACKR1 ICAM1 CCL2 SELE PROX1 TFF3 CCL21
Mean expression in group
0.0 0.5 1.0
Fraction of cells in group (%)
20 40 60 80 100
F
fetal
paediatric
Capsular
Sub-Capsular
Cortical level I
Cortical level II
Cortical level III
Cortical CMJ
Medullar CMJ
Medullar level I
Medullar level II
Medullar level III
EC-Art EC-Art-ELN EC-Cap EC-Cap-Prolif EC-Ven EC-Ven-ELN EC-Lymphatic
EC-Art EC-Art-ELN EC-Cap EC-Ven EC-Lymphatic
Relative cell abundance within group
0.0 0.5 1.0
cutoff > 0.1%
Fraction of spots above cutoff (%)
10 30 50 70 90
Fraction of spots above cutoff (%)
10 20 30 40 50
G
ProlifPericyte
Pericyte
SMC
Pericyte_COL1A1
Pericyte_CCL19
H
SMC
Pericyte
Pericyte_CCL19
Pericyte_COL1A1
ProlifPericyte
4.7k
3.6k
683
596
460
ACTA2 MYH11 RERGL CASQ2 RGS5 PDGFRB ABCC9 KCNJ8 CCL19 CCL21 COL1A1 ELN MKI67 TOP2A
Mean expression in group
0.0 0.5 1.0
Fraction of cells in group (%)
20 40 60 80 100
I
fetal
paediatric
Capsular
Sub-Capsular
Cortical level I
Cortical level II
Cortical level III
Cortical CMJ
Medullar CMJ
Medullar level I
Medullar level II
Medullar level III
Pericyte Pericyte_CCL19 Pericyte_COL1A1 ProlifPericyte SMC
Pericyte Pericyte_CCL19 Pericyte_COL1A1 SMC
Relative cell abundance within group
0.0 0.5 1.0
cutoff > 0.1%
Fraction of spots above cutoff (%)
10 30 50 70 90
Fraction of spots above cutoff (%)
10 20 30 40

**Extended Data Figure 7. Annotation and spatial mapping of thymic fibroblasts and vascular cells. A.** UMAP embedding of thymic fibroblast scRNA-seq data with annotations. **B.** Dot plot showing expression of marker genes in the annotated thymic fibroblast subsets. Total cell numbers per cell type are indicated by bar graph. **C.** Predicted spatial mapping of thymic fibroblasts after CMA calculation and deconvolution of fetal and paediatric Visium data. Cutoff indicates the minimum proportion of the respective cell type in a Visium spot for the spot to be included. Dot size represents the proportion of spots meeting the cutoff and colour indicates the relative cell abundance. **D.** UMAP embedding of thymic endothelial cell scRNA-seq data with annotations. **E.** Dot plot showing expression of marker genes in the annotated thymic endothelial cell subsets. Total cell numbers per cell type are indicated by bar graph. **F.** Predicted spatial mapping of thymic endothelial cells after CMA calculation and deconvolution of fetal and paediatric Visium data. Cutoff indicates the minimum proportion of the respective cell type in a Visium spot for the spot to be included. Dot size represents the proportion of spots meeting the cutoff and colour indicates the relative cell abundance. **G.** UMAP embedding of thymic smooth muscle cell scRNA-seq data with annotations. **H.** Dot plot showing expression of marker genes in the annotated thymic smooth muscle cell subsets. Total cell numbers per cell type are indicated by bar graph. **I.** Predicted spatial mapping of thymic smooth muscle cells after CMA calculation and deconvolution of fetal and paediatric Visium data. Cutoff indicates the minimum proportion of the respective cell type in a Visium spot for the spot to be included. Dot size represents the proportion of spots meeting the cutoff and colour indicates the relative cell abundance.

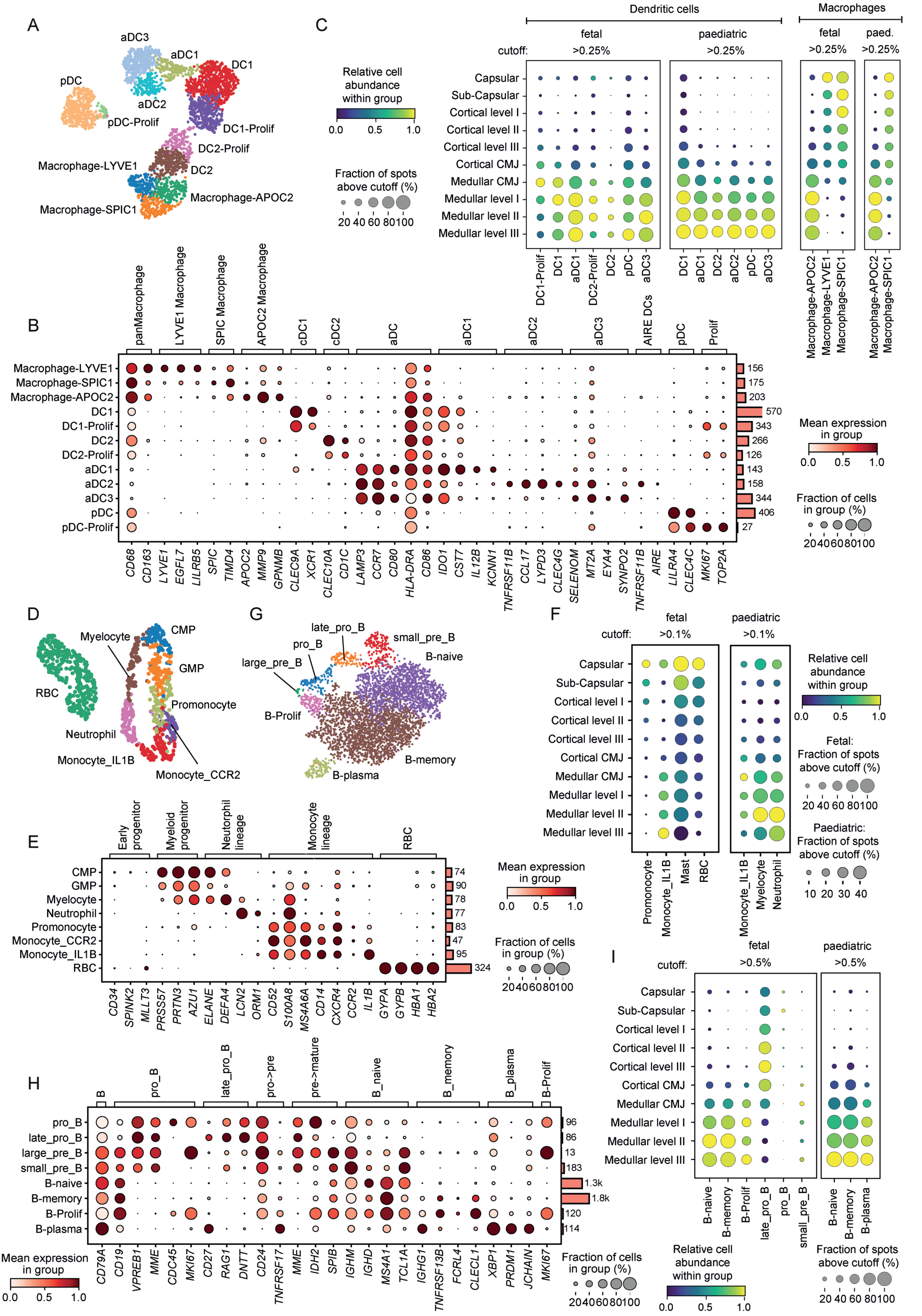

A
aDC3
aDC1
DC1
pDC
aDC2
pDC-Prolif
DC1-Prolif
DC2-Prolif
Macrophage-LYVE1
DC2
Macrophage-APOC2
Macrophage-SPIC1
C
Dendritic cells
Macrophages
fetal
paediatric
fetal
paed.
cutoff:
>0.25%
Relative cell abundance within group
0.0 0.5 1.0
Fraction of spots above cutoff (%)
20 40 60 80100
Capsular
Sub-Capsular
Cortical level I
Cortical level II
Cortical level III
Cortical CMJ
Medullar CMJ
Medullar level I
Medullar level II
Medullar level III
DC1-Prolif
DC1
aDC1
DC2-Prolif
DC2
pDC
aDC3
aDC2
Macrophage-APOC2
Macrophage-LYVE1
Macrophage-SPIC1
B
panMacrophage
LYVE1 Macrophage
SPIC Macrophage
APOC2 Macrophage
cDC1
cDC2
aDC
aDC1
aDC2
aDC3
AIRE DCs
pDC
Prolif
Macrophage-LYVE1
Macrophage-SPIC1
Macrophage-APOC2
DC1
DC1-Prolif
DC2
DC2-Prolif
aDC1
aDC2
aDC3
pDC
pDC-Prolif
156
175
203
570
343
266
126
143
158
344
406
27
CD68 CD163 LYVE1 EGFL7 LILRB5 SPIC TIMD4 APOC2 MMP9 GPNMB CLEC9A XCR1 CLEC10A CD1C LAMP3 CCR7 CD80 HLA-DRA CD86 IDO1 CST7 IL12B KCNN1 TNFRSF11B CCL17 LYPD3 CLEC4G SELENOM MT2A EYA4 SYNPO2 TNFRSF11B AIRE LILRA4 CLEC4C MKI67 TOP2A
Mean expression in group
0.0 0.5 1.0
Fraction of cells in group (%)
20406080100
D
Myelocyte
CMP
GMP
RBC
Promonocyte
Neutrophil
Monocyte_IL1B
Monocyte_CCR2
G
late_pro_B
small_pre_B
pro_B
B-naive
large_pre_B
B-Prolif
B-memory
B-plasma
F
fetal
paediatric
cutoff:
>0.1%
Relative cell abundance within group
0.0 0.5 1.0
Fetal: Fraction of spots above cutoff (%)
20 40 60 80100
Paediatric: Fraction of spots above cutoff (%)
10 20 30 40
Promonocyte
Monocyte_IL1B
Mast
RBC
Monocyte_IL1B
Myelocyte
Neutrophil
E
Early progenitor
Myeloid progenitor
Neutrophil lineage
Monocyte lineage
RBC
CMP
GMP
Myelocyte
Neutrophil
Promonocyte
Monocyte_CCR2
Monocyte_IL1B
RBC
74
90
78
77
83
47
95
324
CD34 SPINK2 MLLT3 PRSS57 PRTN3 AZU1 ELANE DEFA4 LCN2 ORM1 CD52 S100A8 MS4A6A CD14 CXCR4 CCR2 IL1B GYPA GYPB HBA1 HBA2
Mean expression in group
0.0 0.5 1.0
Fraction of cells in group (%)
20406080100
I
fetal
paediatric
cutoff:
>0.5%
B-naive
B-memory
B-Prolif
late_pro_B
pro_B
small_pre_B
B-naive
B-memory
B-plasma
Relative cell abundance within group
0.0 0.5 1.0
Fraction of spots above cutoff (%)
20 40 60 80100
H
B
pro_B
late_pro_B
pro->pre
pre->mature
B_naive
B_memory
B_plasma
B-Prolif
pro_B
late_pro_B
large_pre_B
small_pre_B
B-naive
B-memory
B-Prolif
B-plasma
96
86
13
183
1.3k
1.8k
120
114
CD79A CD19 VPREB1 MME CDC45 MKI67 CD27 RAG1 DNTT CD24 TNFRSF17 MME IDH2 SPIB IGHM IGHD MS4A1 TCL1A IGHG1 TNFRSF13B FCRL4 CLECL1 XBP1 PRDM1 JCHAIN MKI67
Mean expression in group
0.0 0.5 1.0
Fraction of cells in group (%)
20406080100

**Extended Data Figure 8. Annotation and spatial mapping of hematopoietic cells in the fetal and paediatric human thymus. A.** UMAP embedding scRNA-seq data for thymic macrophages and DCs with annotations. **B.** Dot plot showing expression of marker genes in the annotated DC and macrophage subsets. Total cell numbers per cell type are indicated by bar graph. **C.** Predicted spatial mapping of DCs and macrophages after CMA calculation and deconvolution of fetal and paediatric Visium data. Cutoff indicates the minimum proportion of the respective cell type in a Visium spot for the spot to be included. Dot size represents the proportion of spots meeting the cutoff and colour indicates the relative cell abundance. **D.** UMAP embedding of scRNA-seq data for additional thymic myeloid cells with annotations. **E.** Dot plot showing expression of marker genes in the annotated thymic myeloid cell subsets. Total cell numbers per cell type are indicated by bar graph. **F.** Predicted spatial mapping of thymic myeloid cells after CMA calculation and deconvolution of fetal and paediatric Visium data. Cutoff indicates the minimum proportion of the respective cell type in a Visium spot for the spot to be included. Dot size represents the proportion of spots meeting the cutoff and colour indicates the relative cell abundance. **G.** UMAP embedding of thymic B cell scRNA-seq data with annotations. **H.** Dot plot showing expression of marker genes in the annotated thymic B cell subsets. Total cell numbers per cell type are indicated by bar graph. **I.** Predicted spatial mapping of thymic B cells after CMA calculation and deconvolution of fetal and paediatric Visium data. Cutoff indicates the minimum proportion of the respective cell type in a Visium spot for the spot to be included. Dot size represents the proportion of spots meeting the cutoff and colour indicates the relative cell abundance.

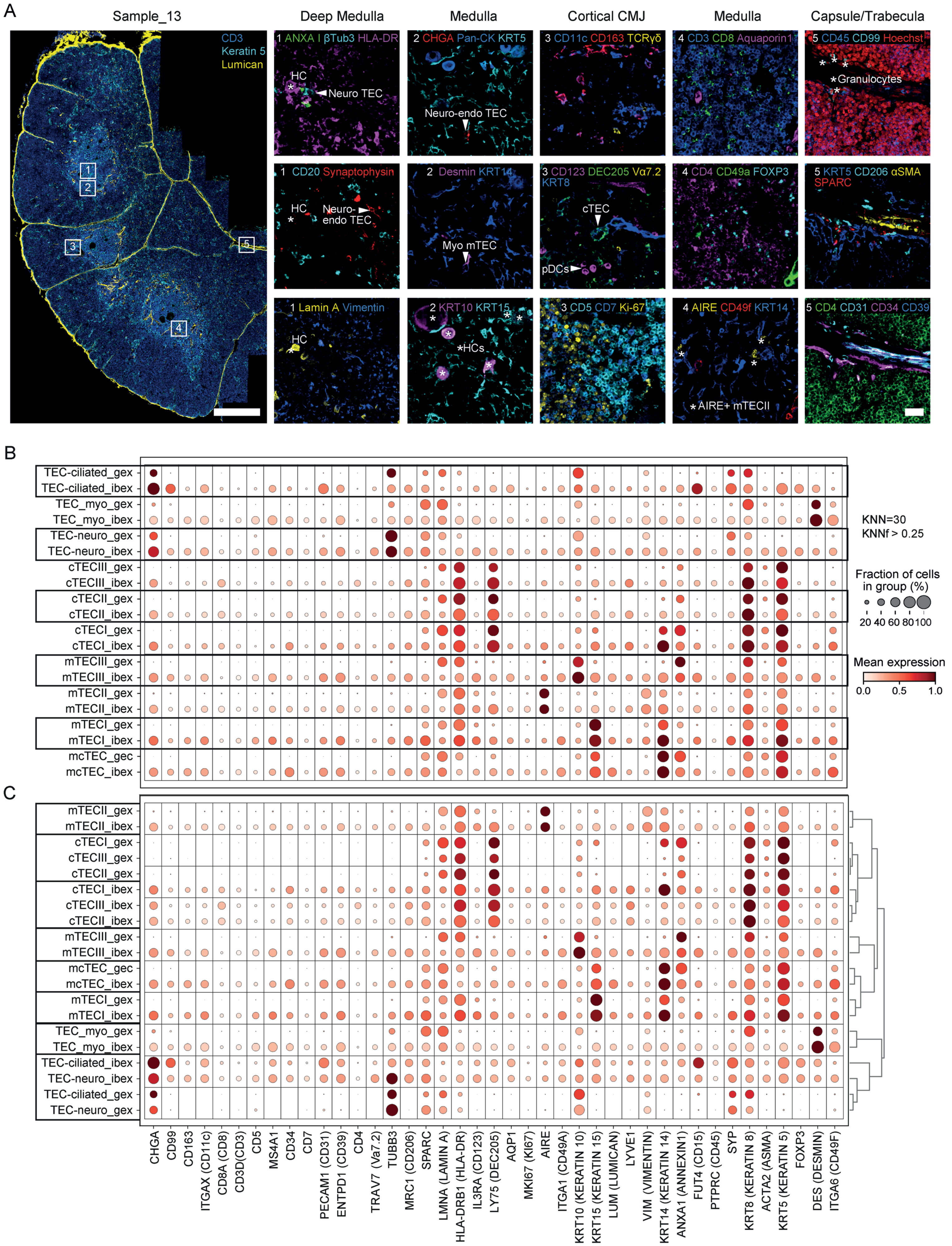
A
Sample_13
CD3
Keratin 5
Lumican
Deep Medulla
Medulla
Cortical CMJ
Medulla
Capsule/Trabecula
1 ANXA I βTub3 HLA-DR
HC
Neuro TEC
2 CHGA Pan-CK KRT5
Neuro-endo TEC
3 CD11c CD163 TCRγδ
4 CD3 CD8 Aquaporin1
5 CD45 CD99 Hoechst
Granulocytes
1 CD20 Synaptophysin
HC
Neuro-endo TEC
2 Desmin KRT14
Myo mTEC
3 CD123 DEC205 Vα7.2 KRT8
cTEC
pDCs
4 CD4 CD49a FOXP3
5 KRT5 CD206 αSMA SPARC
1 Lamin A Vimentin
HC
2 KRT10 KRT15
HCs
3 CD5 CD7 Ki-67
4 AIRE CD49f KRT14
AIRE+ mTECII
5 CD4 CD31 CD34 CD39
B
TEC-ciliated_gex
TEC-ciliated_ibex
TEC_myo_gex
TEC_myo_ibex
TEC-neuro_gex
TEC-neuro_ibex
cTECIII_gex
cTECIII_ibex
cTECII_gex
cTECII_ibex
cTECI_gex
cTECI_ibex
mTECIII_gex
mTECIII_ibex
mTECII_gex
mTECII_ibex
mTECI_gex
mTECI_ibex
mcTEC_gec
mcTEC_ibex
KNN=30
KNNf > 0.25
Fraction of cells in group (%)
20 40 60 80 100
Mean expression
0.0 0.5 1.0
C
mTECII_gex
mTECII_ibex
cTECI_gex
cTECIII_gex
cTECII_gex
cTECI_ibex
cTECIII_ibex
cTECII_ibex
mTECIII_gex
mTECIII_ibex
mcTEC_gec
mcTEC_ibex
mTECI_gex
mTECI_ibex
TEC_myo_gex
TEC_myo_ibex
TEC-ciliated_ibex
TEC-neuro_ibex
TEC-ciliated_gex
TEC-neuro_gex
CHGA
CD99
CD163
ITGAX (CD11c)
CD8A (CD8)
CD3D(CD3)
CD5
MS4A1
CD34
CD7
PECAM1 (CD31)
ENTPD1 (CD39)
CD4
TRAV7 (Vα7.2)
TUBB3
MRC1 (CD206)
SPARC
LMNA (LAMIN A)
HLA-DRB1 (HLA-DR)
IL3RA (CD123)
LY75 (DEC205)
AQP1
MKI67 (KI67)
AIRE
ITGA1 (CD49A)
KRT10 (KERATIN 10)
KRT15 (KERATIN 15)
LUM (LUMICAN)
LYVE1
VIM (VIMENTIN)
KRT14 (KERATIN 14)
ANXA1 (ANNEXIN1)
FUT4 (CD15)
PTPRC (CD45)
SYP
KRT8 (KERATIN 8)
ACTA2 (ASMA)
KRT5 (KERATIN 5)
FOXP3
DES (DESMIN)
ITGA6 (CD49F)

**Extended Data Figure 9. Identification of stromal and TEC subtypes using IBEX imaging. A.** Representative IBEX confocal images from 2-month-old female thymus showing anatomical structures and cell types defined by 44-plex antibody panel. Not shown: CD15 and LYVE-1. Large overview image shows typical region of interest captured in each IBEX experiment (2-3 lobules). Scale bar large overview: 500 μm, scale bars small insets: 25 μm. Annexin I (ANXA I), Chromogranin A (CHGA), pan-cytokeratin (Pan-CK), KRT (Keratin). **B.** Dot plot showing expression of proteins profiled via IBEX and of corresponding genes in scRNA-seq data in TEC subsets. Depicted cell types represent those annotated in scRNA-seq atlas ('_gex', see Figure 4A, B) or the corresponding cell types predicted in IBEX data based on KNN matching ('_ibex'). Expression was normalised per row. Boxes highlight corresponding cell types in the two data sets. **C.** Same as in B but with rows clustered by mutual similarity dendrogram linkage to show similarity level between cell types within and across data sets. Boxes highlight cell types with highest similarity according to dendrogram.

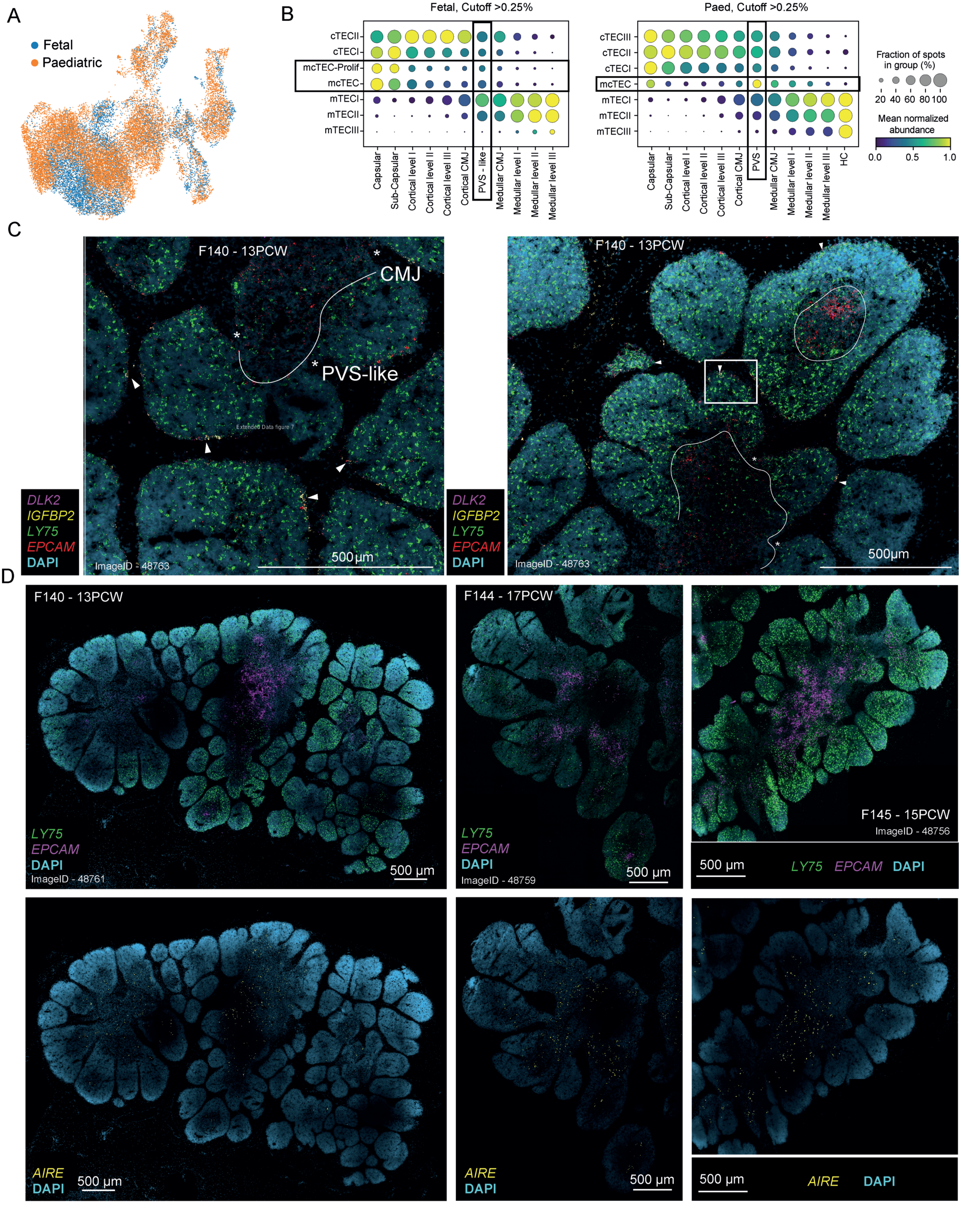

A
Fetal
Paediatric
B
Fetal, Cutoff >0.25%
cTECII
cTECI
mcTEC-Prolif
mcTEC
mTECI
mTECII
mTECIII
Capsular
Sub-Capsular
Cortical level I
Cortical level II
Cortical level III
Cortical CMJ
PVS - like
Medullar CMJ
Medullar level I
Medullar level II
Medullar level III
Paed, Cutoff >0.25%
cTECIII
cTECII
cTECI
mcTEC
mTECI
mTECII
mTECIII
Capsular
Sub-Capsular
Cortical level I
Cortical level II
Cortical level III
Cortical CMJ
PVS
Medullar CMJ
Medullar level I
Medullar level II
Medullar level III
HC
Fraction of spots in group (%)
20 40 60 80 100
Mean normalized abundance
0.0 0.5 1.0
C
F140 - 13PCW
CMJ
PVS-like
DLK2
IGFBP2
LY75
EPCAM
DAPI
ImageID - 48763
500 µm
F140 - 13PCW
DLK2
IGFBP2
LY75
EPCAM
DAPI
ImageID - 48763
500µm
D
F140 - 13PCW
LY75
EPCAM
DAPI
ImageID - 48761
500 µm
F144 - 17PCW
LY75
EPCAM
DAPI
ImageID - 48759
500 µm
F145 - 15PCW
ImageID - 48756
500 µm
LY75 EPCAM DAPI
AIRE
DAPI
500 µm
AIRE
DAPI
500 µm
500 µm
AIRE DAPI

**Extended Data Figure 10. Distribution of mcTEC, mTEC and cTECs in the fetal thymus. A.** UMAP embedding for all TECs in the scRNA-seq data set as shown in Figure 4A, here coloured by donor age. **B.** Relative cell distribution and enrichment of TECs in CMA bins as well as HC and PVS spots based on deconvolved Visium data. Boxes highlight mcTECs and PVS annotations. Note that proliferating mcTECs were only found in fetal thymus and cTECIII was exclusively detected in paediatric data. Cutoff indicates the minimum proportion of the respective cell type in a Visium spot for the spot to be included. Note that “Fetal-HC” is data from a single Visium spot due to the scarcity of HCs in the fetal thymus. **C.** 4-plex RNAscope staining of a thymus tissue section for mcTEC markers *DLK2* (magenta) and *IGFBP2* (yellow) as well as the cTEC marker *LY75* (green) and the mTEC marker *EPCAM* (red). DAPI (cyan) was used to identify nuclei. Arrows indicate capsular mcTECs. Asterisk indicates a location of a structure that is equivalent to the paediatric PVS (invagination zone between the cortex and medulla), although no vascularisation is seen in this region in the fetal thymus. Line indicates the location of the CMJ. Note that mcTEC *DLK2* and *IGFBP6* staining is predominantly capsular and subcapsular and is absent in the PVS-like structure. Inset corresponds to Figure 4G. **D.** RNAscope staining on thymic tissue sections from three individual fetal donors. Staining was performed for the cTEC marker *LY75* (green), the mTEC marker *EPCAM* (magenta), and the mTECII marker *AIRE* (yellow). DAPI (cyan) was used for identification of nuclei.

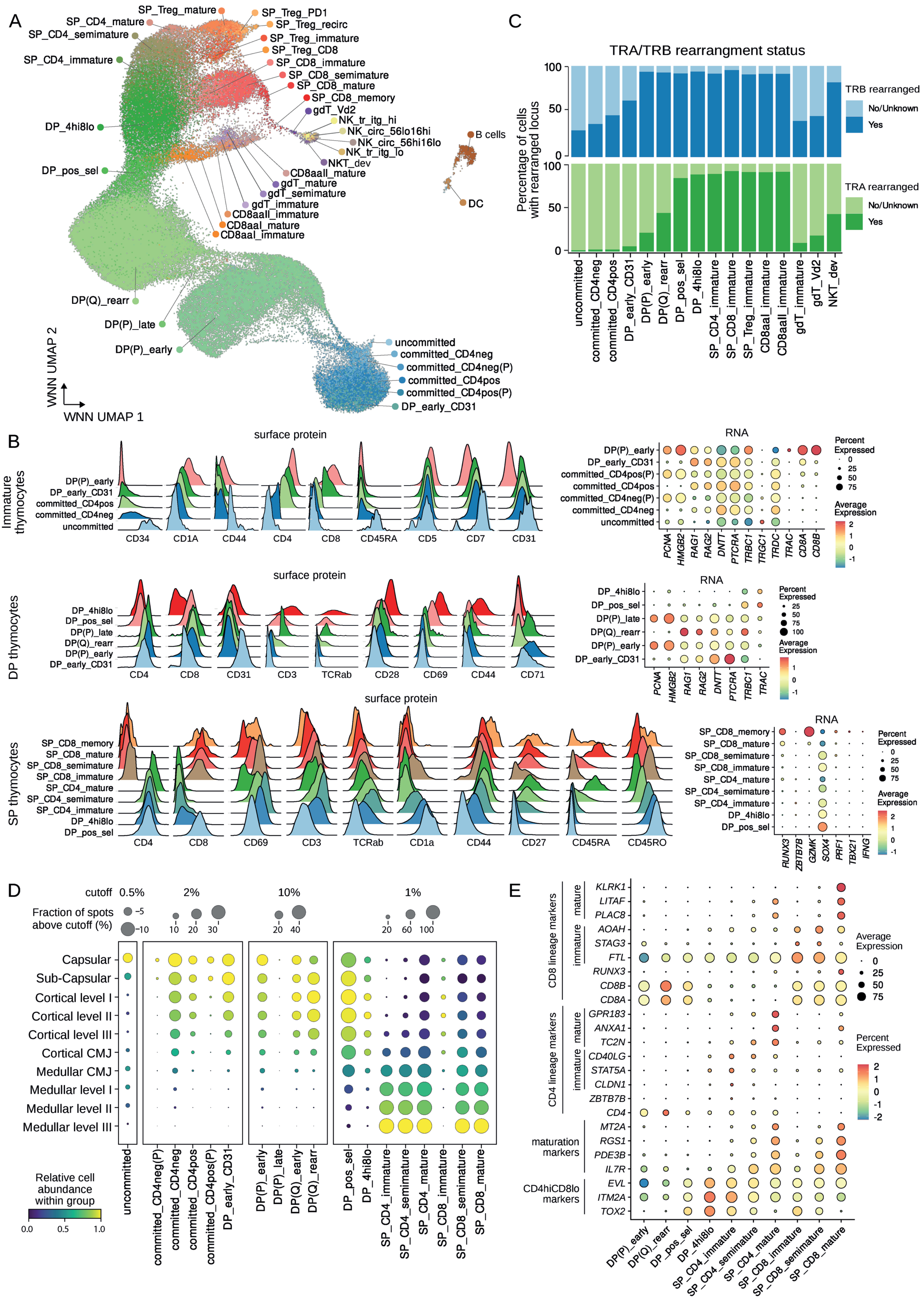
A
SP_Treg_mature
SP_CD4_mature
SP_CD4_semimature
SP_CD4_immature
DP_4hi8lo
DP_pos_sel
DP(Q)_rearr
DP(P)_late
DP(P)_early
SP_Treg_PD1
SP_Treg_recirc
SP_Treg_immature
SP_Treg_CD8
SP_CD8_immature
SP_CD8_semimature
SP_CD8_mature
SP_CD8_memory
gdT_Vd2
NK_tr_itg_hi
NK_circ_56lo16hi
NK_circ_56hi16lo
NK_tr_itg_lo
NKT_dev
CD8aaII_mature
gdT_mature
gdT_semimature
gdT_immature
CD8aaII_immature
CD8aaI_mature
CD8aaI_immature
B cells
DC
uncommitted
committed_CD4neg
committed_CD4neg(P)
committed_CD4pos
committed_CD4pos(P)
DP_early_CD31
WNN UMAP 2
WNN UMAP 1
C
TRA/TRB rearrangment status
Percentage of cells with rearranged locus
TRB rearranged
No/Unknown
Yes
TRA rearranged
No/Unknown
Yes
uncommitted
committed_CD4neg
committed_CD4pos
DP_early_CD31
DP(P)_early
DP(Q)_rearr
DP_pos_sel
DP_4hi8lo
SP_CD4_immature
SP_CD8_immature
SP_Treg_immature
CD8aaI_immature
CD8aaII_immature
gdT_immature
gdT_Vd2
NKT_dev
B
Immature thymocytes
surface protein
DP(P)_early
DP_early_CD31
committed_CD4pos
committed_CD4neg
uncommitted
CD34 CD1A CD44 CD4 CD8 CD45RA CD5 CD7 CD31
RNA
DP(P)_early
DP_early_CD31
committed_CD4pos(P)
committed_CD4pos
committed_CD4neg(P)
committed_CD4neg
uncommitted
PCNA HMGB2 RAG1 RAG2 DNTT PTCRA TRBC1 TRGC1 TRDC TRAC CD8A CD8B
Percent Expressed
Average Expression
DP thymocytes
surface protein
DP_4hi8lo
DP_pos_sel
DP(P)_late
DP(Q)_rearr
DP(P)_early
DP_early_CD31
CD4 CD8 CD31 CD3 TCRab CD28 CD69 CD44 CD71
RNA
PCNA HMGB2 RAG1 RAG2 DNTT PTCRA TRBC1 TRAC
SP thymocytes
surface protein
SP_CD8_memory
SP_CD8_mature
SP_CD8_semimature
SP_CD8_immature
SP_CD4_mature
SP_CD4_semimature
SP_CD4_immature
DP_4hi8lo
DP_pos_sel
CD4 CD8 CD69 CD3 TCRab CD1a CD44 CD27 CD45RA CD45RO
RNA
RUNX3 ZBTB7B GZMK SOX4 PRF1 TBX21 IFNG
D
cutoff 0.5% 2% 10% 1%
Fraction of spots above cutoff (%)
Capsular
Sub-Capsular
Cortical level I
Cortical level II
Cortical level III
Cortical CMJ
Medullar CMJ
Medullar level I
Medullar level II
Medullar level III
Relative cell abundance within group
uncommitted
committed_CD4neg(P)
committed_CD4neg
committed_CD4pos
committed_CD4pos(P)
DP_early_CD31
DP(P)_early
DP(P)_late
DP(Q)_early
DP(Q)_rearr
DP_pos_sel
DP_4hi8lo
SP_CD4_immature
SP_CD4_semimature
SP_CD4_mature
SP_CD8_immature
SP_CD8_semimature
SP_CD8_mature
E
CD8 lineage markers
mature
immature
KLRK1 LITAF PLAC8 AOAH STAG3 FTL RUNX3 CD8B CD8A
CD4 lineage markers
GPR183 ANXA1 TC2N CD40LG STAT5A CLDN1 ZBTB7B CD4
maturation markers
MT2A RGS1 PDE3B IL7R
CD4hiCD8lo markers
EVL ITM2A TOX2
Average Expression
Percent Expressed
DP(P)_early
DP(Q)_rearr
DP_pos_sel
DP_4hi8lo
SP_CD4_immature
SP_CD4_semimature
SP_CD4_mature
SP_CD8_immature
SP_CD8_semimature
SP_CD8_mature

**Extended Data Figure 11. Surface protein-guided annotations and spatial mapping of maturation stages of conventional T lineage cells in the paediatric thymus. A.** WNN UMAP of the full CITE-seq dataset with cell annotations. **B.** Expression of selected markers relevant for the distinction and annotation of differentiation stages of immature, DP and SP thymocytes. Density histograms indicate surface protein levels after denoising and normalisation (left). Dot plots show scaled normalised pseudo bulk RNA levels for selected relevant genes (right). DP: double positive, SP: single positive, pos_sel: positive-selected, (P): proliferating, (Q): quiescent, rearr: TCR-rearranging. **C.** Bar graph illustrating the proportion of cells with rearranged *TRB* and *TRA* locus at selected maturation stages based on TCR-seq data after analysis with Dandelion. **D.** CMA mapping of the annotated stages of conventional T lineage development by using cell2location with paediatric Visium sections. Stages were plotted separately with adjusted cutoff to aid visualisation of smaller subsets. Cutoff indicates the minimum proportion of the respective cell type in a Visium spot for the spot to be included. Dot size represents the proportion of spots meeting the cutoff and colour indicates the relative cell abundance. **E.** Marker gene expression in the surface protein-derived maturation stages of conventional T cells from DP stage onwards.

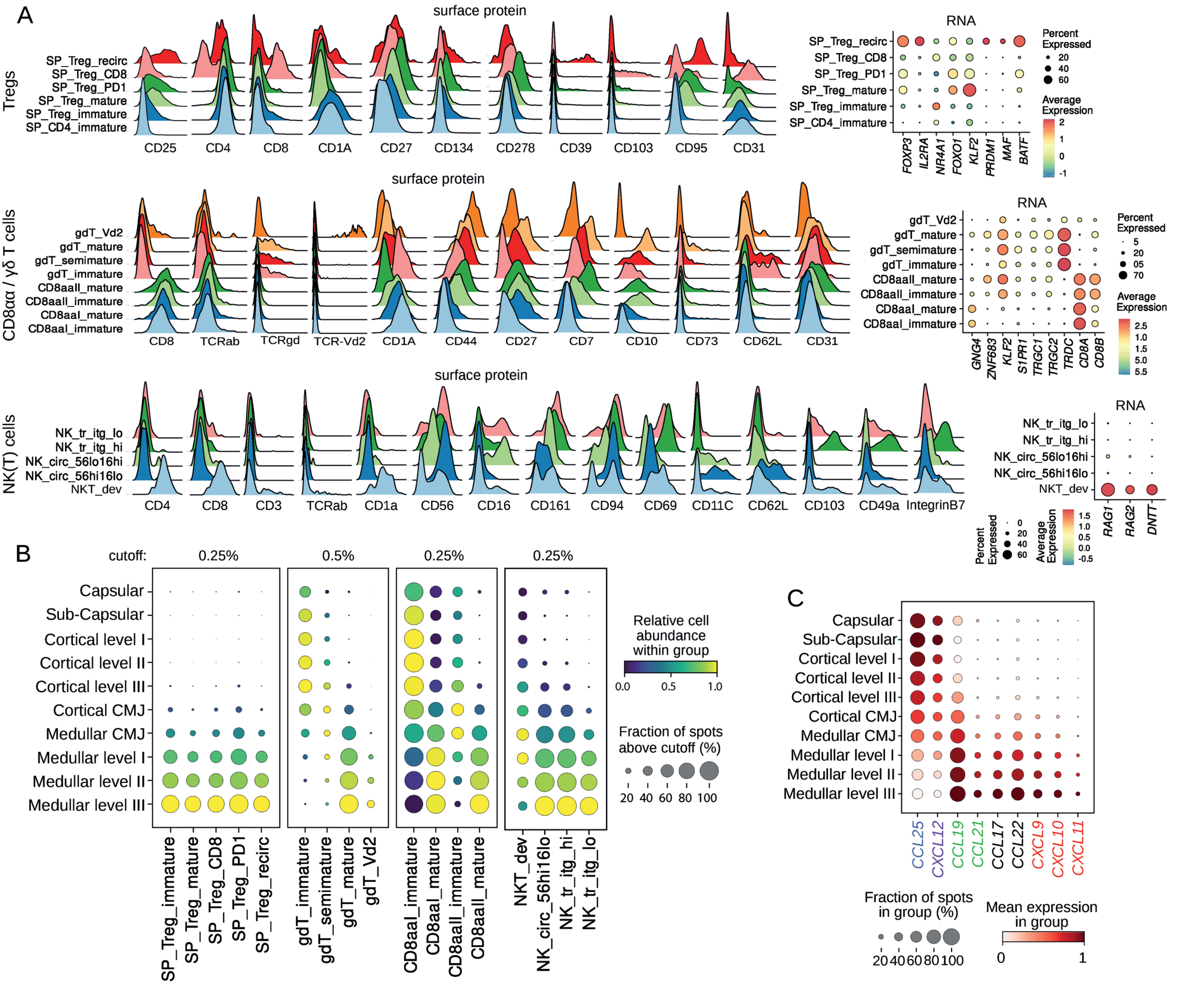
A
surface protein
Tregs
SP_Treg_recirc
SP_Treg_CD8
SP_Treg_PD1
SP_Treg_mature
SP_Treg_immature
SP_CD4_immature
CD25
CD4
CD8
CD1A
CD27
CD134
CD278
CD39
CD103
CD95
CD31
RNA
FOXP3
IL2RA
NR4A1
FOXO1
KLF2
PRDM1
MAF
BATF
Percent Expressed
20
40
60
Average Expression
2
1
0
-1
surface protein
CD8αα / γδ T cells
gdT_Vd2
gdT_mature
gdT_semimature
gdT_immature
CD8aaII_mature
CD8aaII_immature
CD8aaI_mature
CD8aaI_immature
CD8
TCRab
TCRgd
TCR-Vd2
CD1A
CD44
CD27
CD7
CD10
CD73
CD62L
CD31
RNA
GNG4
ZNF683
KLF2
S1PR1
TRGC1
TRGC2
TRDC
CD8A
CD8B
Percent Expressed
5
20
05
70
Average Expression
2.5
1.0
1.5
5.0
5.5
surface protein
NK(T) cells
NK_tr_itg_lo
NK_tr_itg_hi
NK_circ_56lo16hi
NK_circ_56hi16lo
NKT_dev
CD4
CD8
CD3
TCRab
CD1a
CD56
CD16
CD161
CD94
CD69
CD11C
CD62L
CD103
CD49a
IntegrinB7
RNA
RAG1
RAG2
DNTT
Percent Expressed
0
20
40
60
Average Expression
1.5
1.0
0.5
0.0
-0.5
B
cutoff:
0.25%
0.5%
0.25%
0.25%
Capsular
Sub-Capsular
Cortical level I
Cortical level II
Cortical level III
Cortical CMJ
Medullar CMJ
Medullar level I
Medullar level II
Medullar level III
SP_Treg_immature
SP_Treg_mature
SP_Treg_CD8
SP_Treg_PD1
SP_Treg_recirc
gdT_immature
gdT_semimature
gdT_mature
gdT_Vd2
CD8aaI_immature
CD8aaI_mature
CD8aaII_immature
CD8aaII_mature
NKT_dev
NK_circ_56hi16lo
NK_tr_itg_hi
NK_tr_itg_lo
Relative cell abundance within group
0.0
0.5
1.0
Fraction of spots above cutoff (%)
20
40
60
80
100
C
Capsular
Sub-Capsular
Cortical level I
Cortical level II
Cortical level III
Cortical CMJ
Medullar CMJ
Medullar level I
Medullar level II
Medullar level III
CCL25
CXCL12
CCL19
CCL21
CCL17
CCL22
CXCL9
CXCL10
CXCL11
Fraction of spots in group (%)
20
40
60
80
100
Mean expression in group
0
1

**Extended Data Figure 12. Surface protein-guided annotations and spatial mapping of unconventional T and NK cells. A.** Expression of selected markers relevant for the distinction and annotation of subtypes and differentiation stages of regulatory T cells, CD8αα T cells, γδ T cells, and NK(T) cells. Density histograms indicate surface protein levels after denoising and normalisation (left). Dot plots show scaled normalised pseudo bulk RNA levels for selected relevant genes (right). Recirc: recirculating, tr: tissue resident, circ: circulating, itg: integrin. **B.** CMA mapping of the annotated stages of conventional T lineage development by using cell2location with paediatric Visium sections. Subtypes were plotted separately with adjusted cutoff to aid visualisation of rare subsets. Cutoff indicates the minimum proportion of the respective cell type in a Visium spot for the spot to be included. Dot size represents the proportion of spots meeting the cutoff and colour indicates the relative cell abundance. **C.** Expression of key chemokines along the CMA as inferred from Visium spot data. Label colours indicate corresponding chemokines for the receptors shown in Figure 6F.
