## Supplementary material for "A spatial human thymus cell atlas mapped to a continuous tissue axis"

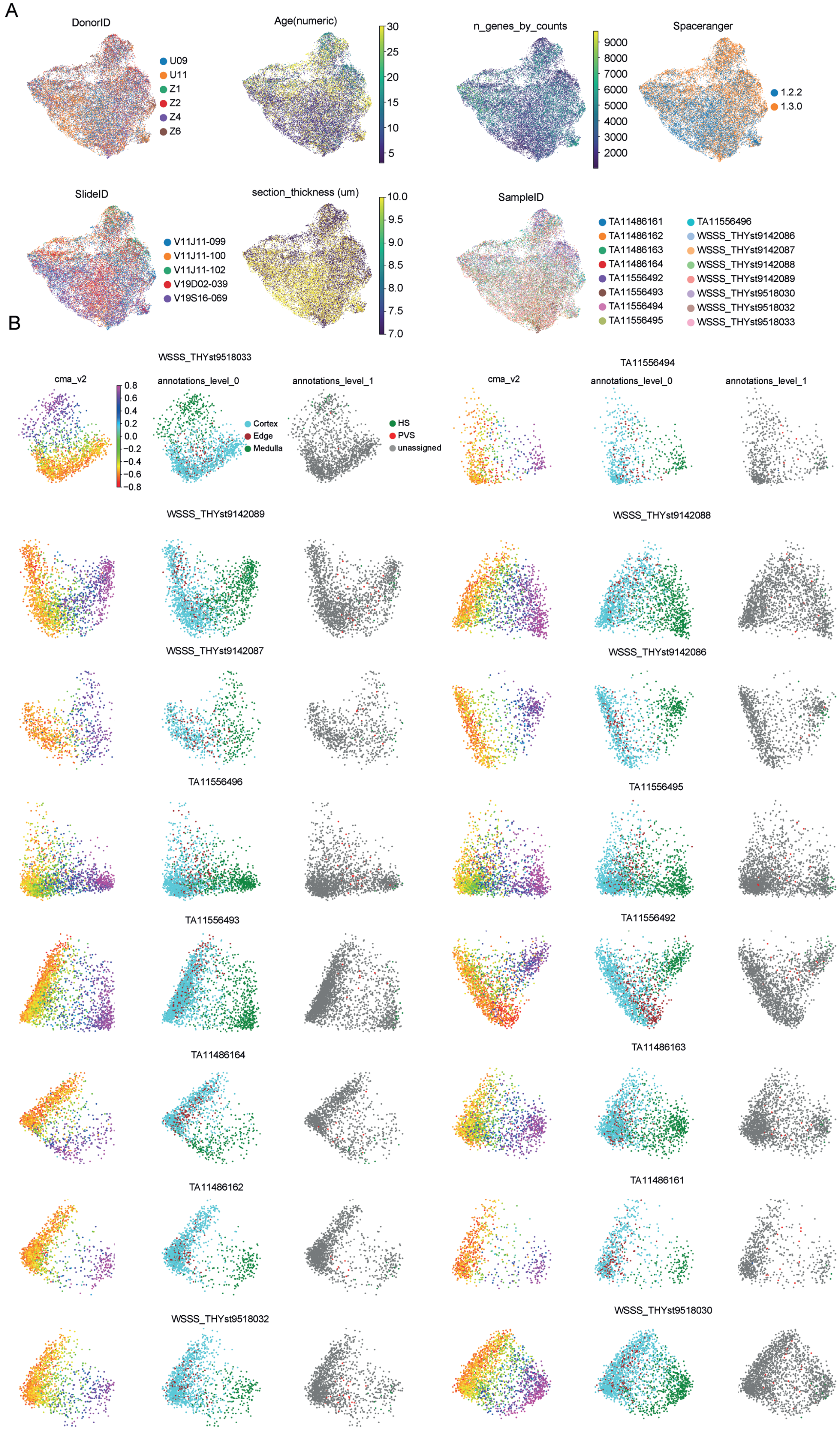
A
DonorID
U09
U11
Z1
Z2
Z4
Z6
Age(numeric)
30
25
20
15
10
5
n_genes_by_counts
9000
8000
7000
6000
5000
4000
3000
2000
Spaceranger
1.2.2
1.3.0
SlideID
V11J11-099
V11J11-100
V11J11-102
V19D02-039
V19S16-069
section_thickness (um)
10.0
9.5
9.0
8.5
8.0
7.5
7.0
SampleID
TA11486161
TA11486162
TA11486163
TA11486164
TA11556492
TA11556493
TA11556494
TA11556495
TA11556496
WSSS_THYst9142086
WSSS_THYst9142087
WSSS_THYst9142088
WSSS_THYst9142089
WSSS_THYst9518030
WSSS_THYst9518032
WSSS_THYst9518033
B
WSSS_THYst9518033
cma_v2
0.8
0.6
0.4
0.2
0.0
−0.2
−0.4
−0.6
−0.8
annotations_level_0
Cortex
Edge
Medulla
annotations_level_1
HS
PVS
unassigned
TA11556494
cma_v2
annotations_level_0
annotations_level_1
WSSS_THYst9142089
WSSS_THYst9142088
WSSS_THYst9142087
WSSS_THYst9142086
TA11556496
TA11556495
TA11556493
TA11556492
TA11486164
TA11486163
TA11486162
TA11486161
WSSS_THYst9518032
WSSS_THYst9518030

**Supplementary Figure 1. Meta data and annotations of paediatric Visium samples. A.** Donor meta data and sample composition for paediatric Visium data illustrated in UMAP space. **B.** PCA plots for 16 individual paediatric Visium samples highlighting the major tissue annotations: CMA, annotation_level_0 and annotation_level_1.

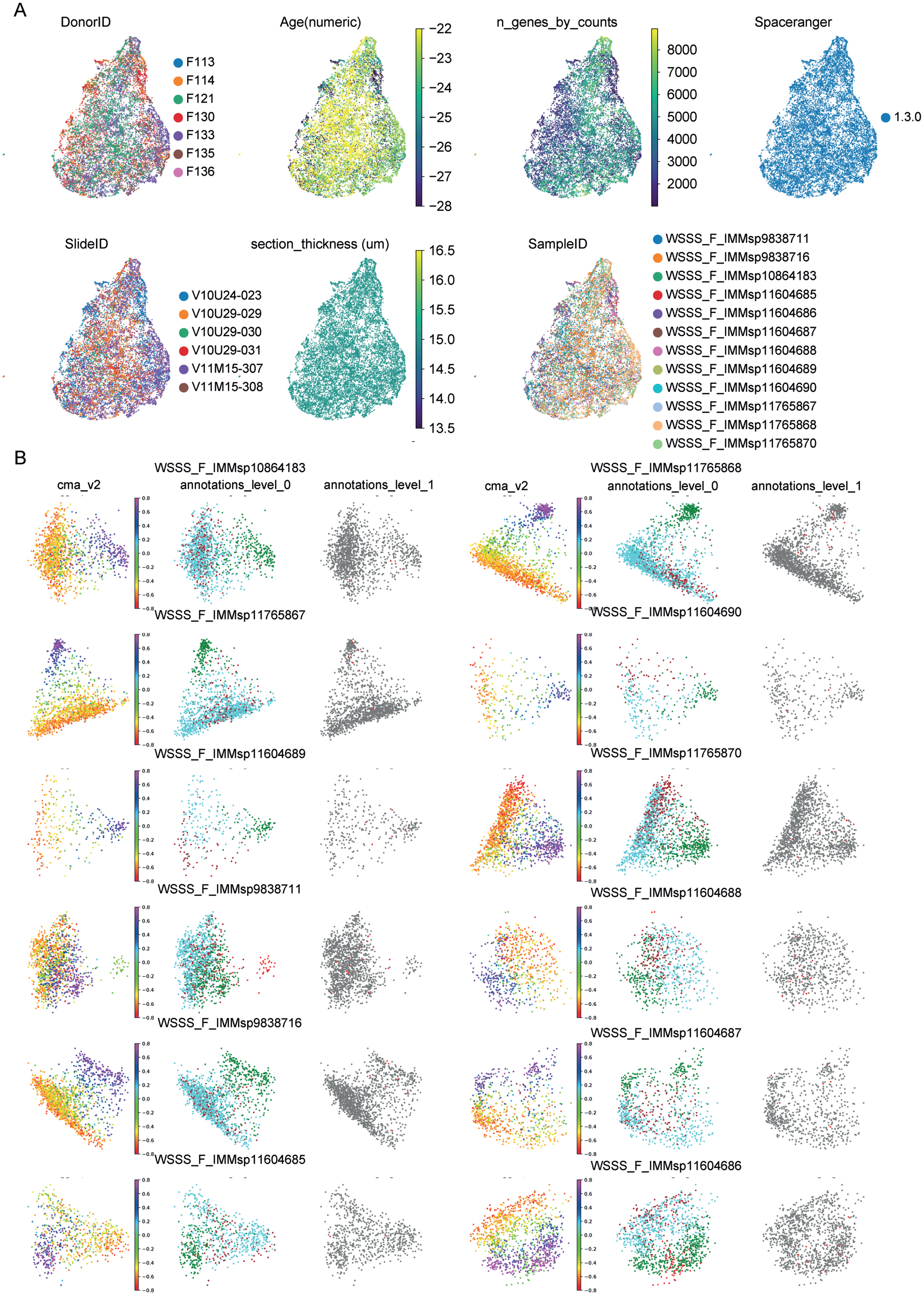
A
DonorID
F113
F114
F121
F130
F133
F135
F136
Age(numeric)
−22
−23
−24
−25
−26
−27
−28
n_genes_by_counts
8000
7000
6000
5000
4000
3000
2000
Spaceranger
1.3.0
SlideID
V10U24-023
V10U29-029
V10U29-030
V10U29-031
V11M15-307
V11M15-308
section_thickness (um)
16.5
16.0
15.5
15.0
14.5
14.0
13.5
SampleID
WSSS_F_IMMsp9838711
WSSS_F_IMMsp9838716
WSSS_F_IMMsp10864183
WSSS_F_IMMsp11604685
WSSS_F_IMMsp11604686
WSSS_F_IMMsp11604687
WSSS_F_IMMsp11604688
WSSS_F_IMMsp11604689
WSSS_F_IMMsp11604690
WSSS_F_IMMsp11765867
WSSS_F_IMMsp11765868
WSSS_F_IMMsp11765870
B
WSSS_F_IMMsp10864183
cma_v2
annotations_level_0
annotations_level_1
WSSS_F_IMMsp11765868
cma_v2
annotations_level_0
annotations_level_1
WSSS_F_IMMsp11765867
WSSS_F_IMMsp11604690
WSSS_F_IMMsp11604689
WSSS_F_IMMsp11765870
WSSS_F_IMMsp9838711
WSSS_F_IMMsp11604688
WSSS_F_IMMsp9838716
WSSS_F_IMMsp11604687
WSSS_F_IMMsp11604685
WSSS_F_IMMsp11604686

**Supplementary Figure 2. Meta data and annotations of fetal Visium samples. A.** Donor meta data and sample composition for fetal Visium data illustrated in UMAP space. **B.** PCA plots for 12 individual fetal Visium samples highlighting the major tissue annotations: CMA, annotation_level_0 and annotation_level_1.

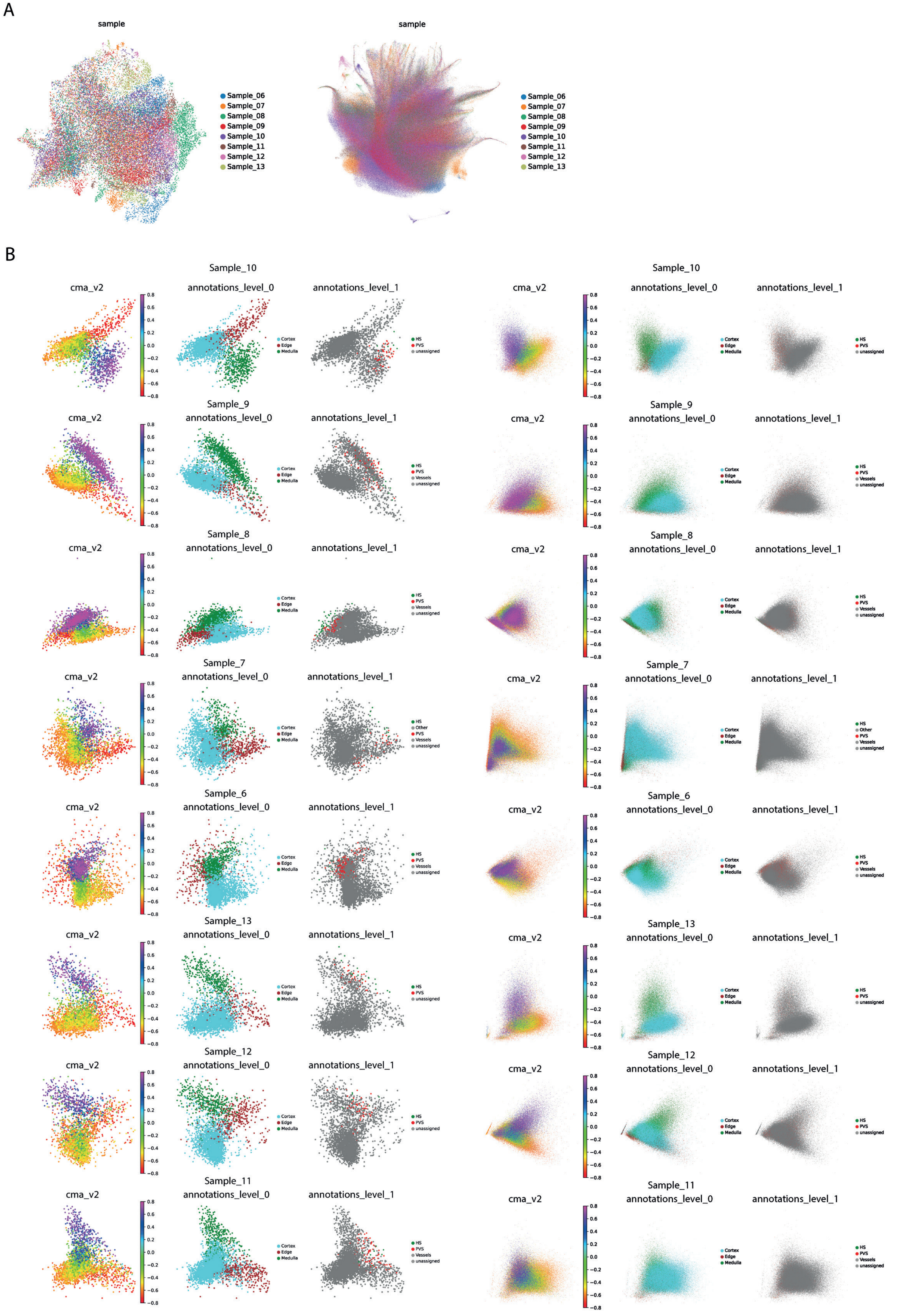
A
sample
Sample_06
Sample_07
Sample_08
Sample_09
Sample_10
Sample_11
Sample_12
Sample_13
B
Sample_10
Sample_9
Sample_8
Sample_7
Sample_6
Sample_13
Sample_12
Sample_11
cma_v2
annotations_level_0
annotations_level_1
Cortex
Edge
Medulla
HS
Other
PVS
Vessels
unassigned

**Supplementary Figure 3. Meta data and annotations of paediatric IBEX 50 µm spot grid and single cell segmentation data. A.** Donor meta data and sample composition for paediatric IBEX spot grid (left) and single cell (right) data illustrated in UMAP space. **B.** PCA plots for spot grid (left) and single cell (right) data from 8 individual paediatric IBEX samples highlighting the major tissue annotations: CMA, annotation_level_0 and annotation_level_1.

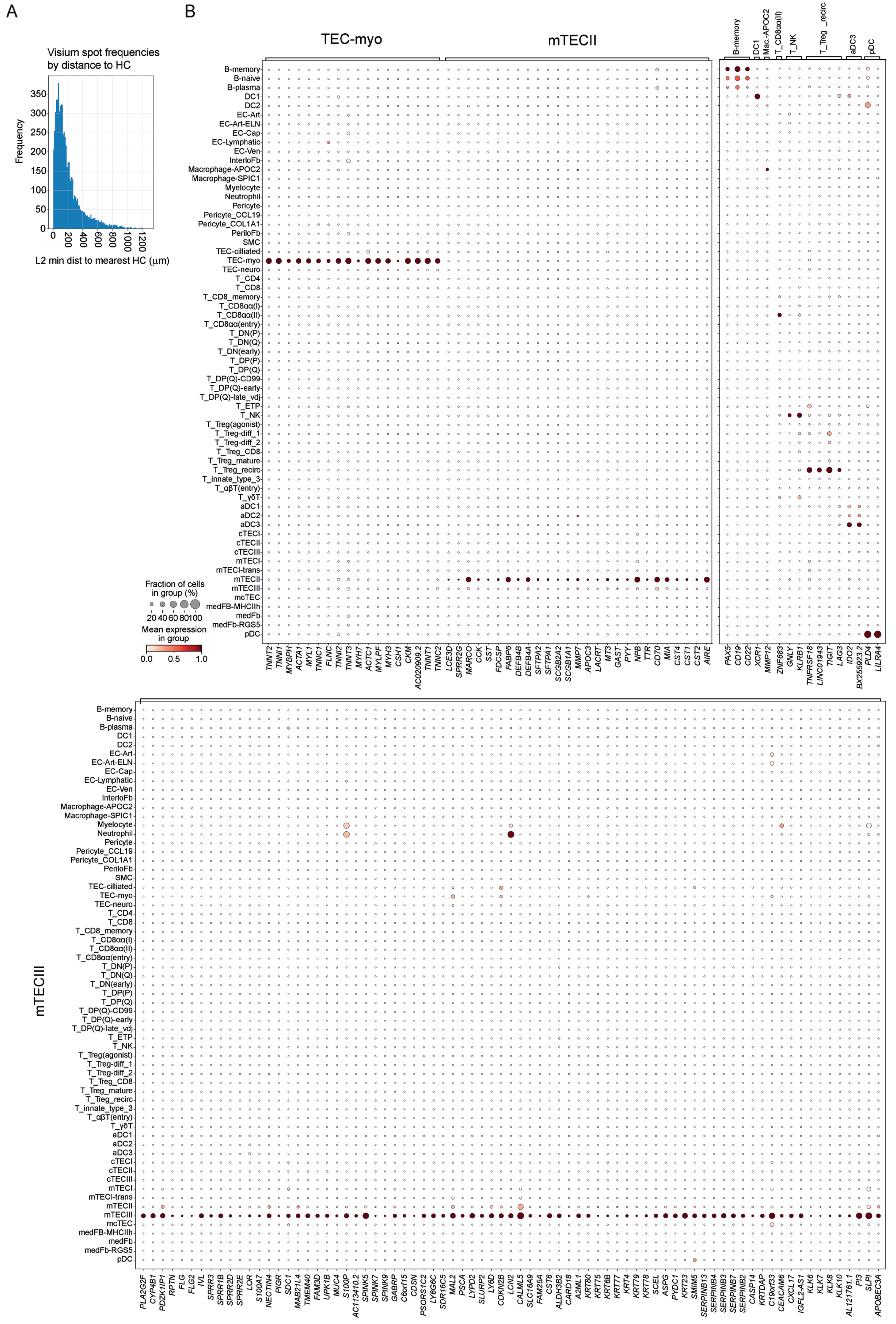
A
B
Visium spot frequencies by distance to HC
Frequency
L2 min dist to mearest HC (μm)
TEC-myo
mTECII
mTECIII
B-memory
DC1
Mac.-APOC2
T_CD8αα(II)
T_NK
T_Treg _recirc
aDC3
pDC
Fraction of cells in group (%)
20 40 60 80 100
Mean expression in group
0.0 0.5 1.0
TNNT2 TNNI1 MYBPH ACTA1 MYL1 TNNC1 FLNC TNNI2 TNNT3 MYH7 ACTC1 MYLPF MYH3 CSH1 CKM AC020909.2 TNNT1 TNNC2
LCE3D SPRR2G MARCO CCK SST FDCSP FABP6 DEFB4B DEFB4A SFTPA2 SFTPA1 SCGB2A2 SCGB1A1 MMP7 APOC3 LACRT MT3 GAST PYY NPB TTR CD70 MIA CST4 CST1 CST2 AIRE
PAX5 CD19 CD22 XCR1 MMP12 ZNF683 GNLY KLRB1 TNFRSF18 LINC01943 TIGIT LAG3 IDO2 BX255923.2 PLD4 LILRA4
PLA2G2F CYP4B1 PDZK1IP1 RPTN FLG FLG2 IVL SPRR3 SPRR1B SPRR2D SPRR2E LOR S100A7 NECTIN4 PIGR SDC1 MAB21L4 TMEM40 FAM3D UPK1B MUC4 S100P AC113410.2 SPINK5 SPINK7 SPINK9 GABRP C6orf15 CDSN PSORS1C2 LY6G6C SDR16C5 MAL2 PSCA LYPD2 SLURP2 LY6D CDKN2B LCN2 CALML5 SLC16A9 FAM25A CST6 ALDH3B2 CARD18 A2ML1 KRT80 KRT75 KRT6B KRT77 KRT4 KRT79 KRT78 SCEL ASPG PYDC1 KRT23 SMIM5 SERPINB13 SERPINB4 SERPINB3 SERPINB7 SERPINB2 CASP14 KRTDAP C19orf33 CEACAM6 CXCL17 IGFL2-AS1 KLK6 KLK7 KLK8 KLK10 AL121761.1 PI3 SLPI APOBEC3A

**Supplementary Figure 4. Analysis of HCs in the paediatric thymus. A**. Histogram showing the frequency distribution of medullary spots by distance from HCs. **B** Specialisation genes selected by unique expression profiles within a cell type shown across all cell types. Note the high proportion of genes for mTECIII cells.

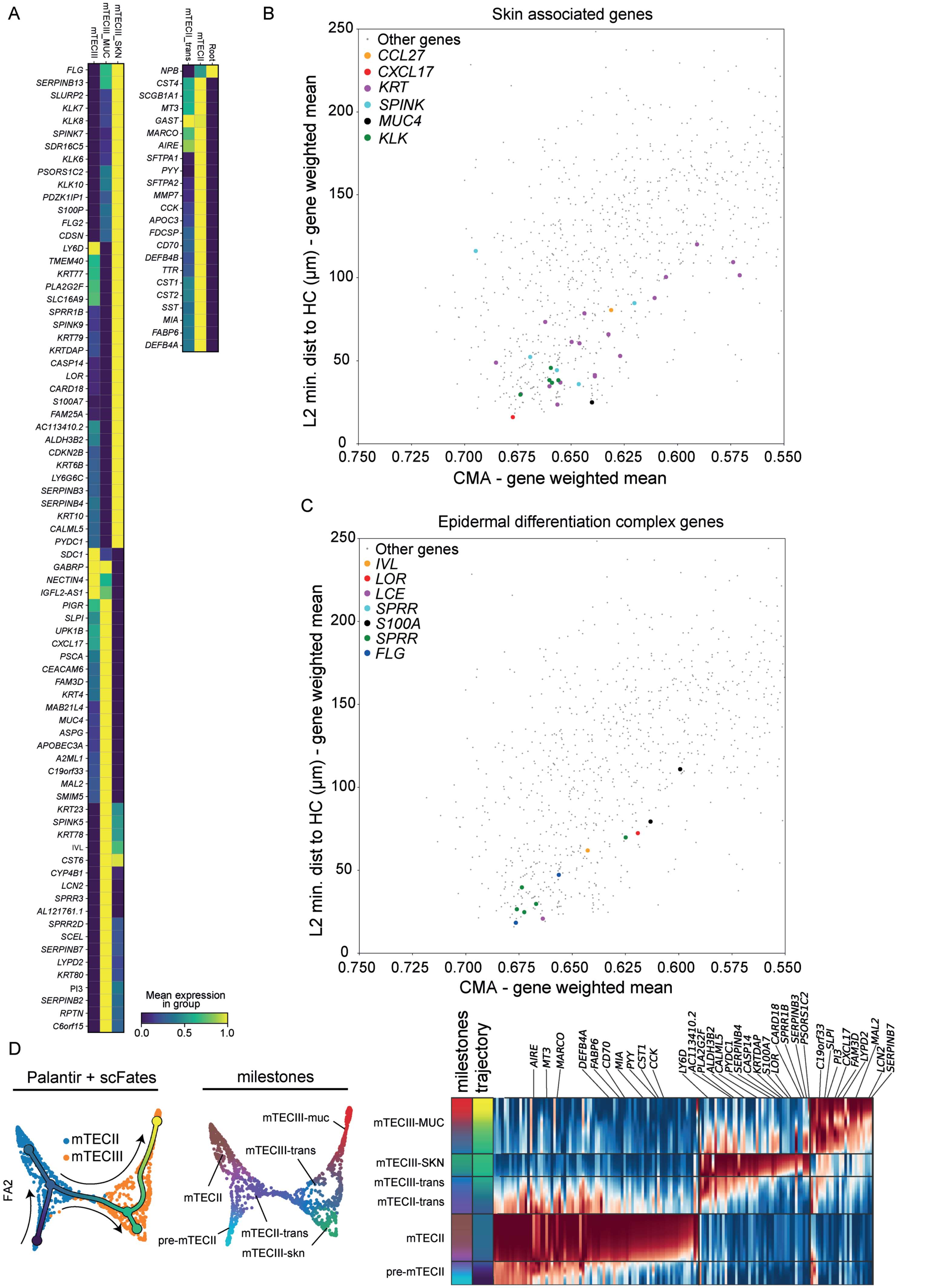
A
mTECIII
mTECIII_MUC
mTECIII_SKN
FLG
SERPINB13
SLURP2
KLK7
KLK8
SPINK7
SDR16C5
KLK6
PSORS1C2
KLK10
PDZK1IP1
S100P
FLG2
CDSN
LY6D
TMEM40
KRT77
PLA2G2F
SLC16A9
SPRR1B
SPINK9
KRT79
KRTDAP
CASP14
LOR
CARD18
S100A7
FAM25A
AC113410.2
ALDH3B2
CDKN2B
KRT6B
LY6G6C
SERPINB3
SERPINB4
KRT10
CALML5
PYDC1
SDC1
GABRP
NECTIN4
IGFL2-AS1
PIGR
SLPI
UPK1B
CXCL17
PSCA
CEACAM6
FAM3D
KRT4
MAB21L4
MUC4
ASPG
APOBEC3A
A2ML1
C19orf33
MAL2
SMIM5
KRT23
SPINK5
KRT78
IVL
CST6
CYP4B1
LCN2
SPRR3
AL121761.1
SPRR2D
SCEL
SERPINB7
LYPD2
KRT80
PI3
SERPINB2
RPTN
C6orf15
mTECII_trans
mTECII
Root
NPB
CST4
SCGB1A1
MT3
GAST
MARCO
AIRE
SFTPA1
PYY
SFTPA2
MMP7
CCK
APOC3
FDCSP
CD70
DEFB4B
TTR
CST1
CST2
SST
MIA
FABP6
DEFB4A
Mean expression in group
0.0
0.5
1.0
B
Skin associated genes
Other genes
CCL27
CXCL17
KRT
SPINK
MUC4
KLK
L2 min. dist to HC (μm) - gene weighted mean
CMA - gene weighted mean
250
200
150
100
50
0
0.750
0.725
0.700
0.675
0.650
0.625
0.600
0.575
0.550
C
Epidermal differentiation complex genes
Other genes
IVL
LOR
LCE
SPRR
S100A
SPRR
FLG
L2 min. dist to HC (μm) - gene weighted mean
CMA - gene weighted mean
D
Palantir + scFates
mTECII
mTECIII
FA2
milestones
mTECIII-muc
mTECIII-trans
mTECII
pre-mTECII
mTECII-trans
mTECIII-skn
milestones
trajectory
AIRE
MT3
MARCO
DEFB4A
FABP6
CD70
MIA
PYY
CST1
CCK
LY6D
AC113410.2
PLA2G2F
ALDH3B2
CALML5
PYDC1
SERPINB4
CASP14
KRTDAP
S100A7
LOR
CARD18
SPRR1B
SERPINB3
PSORS1C2
C19orf33
SLPI
PI3
CXCL17
FAM3D
LYPD2
MAL2
LCN2
SERPINB7
mTECIII-MUC
mTECIII-SKN
mTECIII-trans
mTECII-trans
mTECII
pre-mTECII

**Supplementary Figure 5. Expression and spatial localisation of SGs. A.** SGs of mTECIII (left) and mTECII (right) subgroups defined by trajectory analysis in (D). **B**. Distribution of all 851 medullary genes along the CMA and with respect to HC distance. Skin related gene families are represented by large dots, which are coloured according to the gene category. **C.** Distribution of all 851 medullary genes along the CMA and with respect to HC distance. Epidermal differentiation complex gene families are represented by large dots, which are coloured according to the gene category. **D.** scFates trajectory analysis on mTECII and mTECIII as in Figure 5H-J. Gene enrichment analysis for distinct expression patterns of genes across the cell trajectory is shown.

A

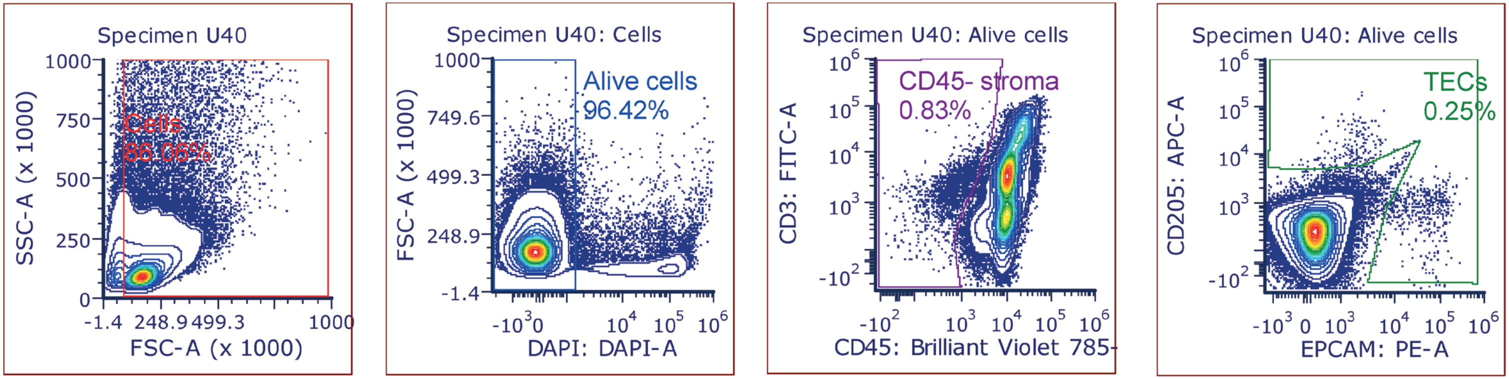


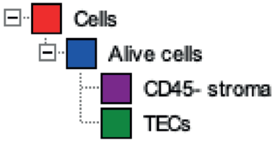


| Gate | # of Events | X Geometric Mean | Y Geometric Mean | % of all cells |
|---|---|---|---|---|
| No Gate | 304395 | n/a | 118683.54 | 100.00 |
| Cells | 261969 | 176065.37 | 112319.54 | 86.06 |
| Alive cells | 252590 | 178393.42 | 109786.99 | 82.98 |
| CD45- stroma | 2109 | 224202.52 | 359447.29 | 0.69 |
| TECs | 629 | 304347.30 | 410541.04 | 0.21 |

B

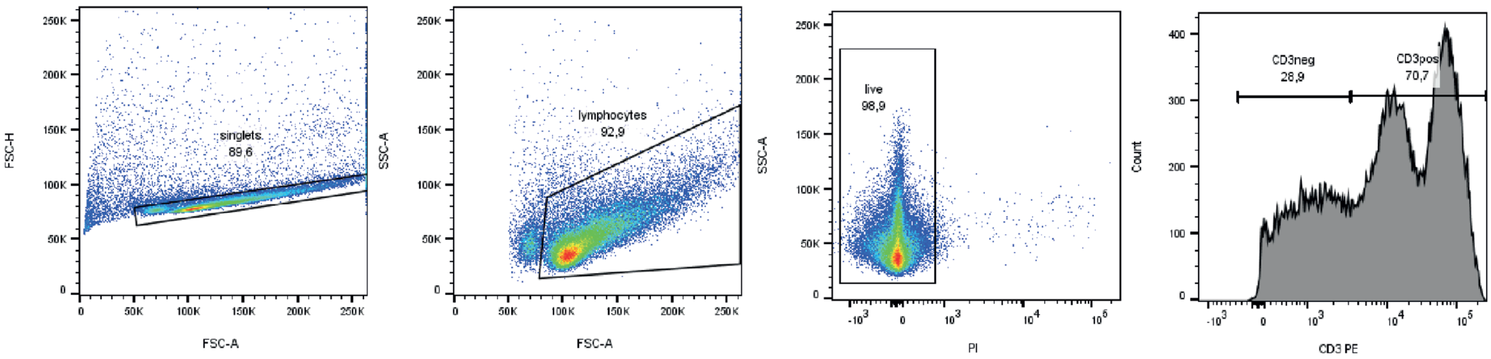

**Supplementary Figure 6. Gating strategy for FACS sorting. A.** Sorting scheme for stroma enrichment experiments. Cells were stained with a mix of anti-EPCAM PE, anti-CD205 APC, anti-CD45 BV785, anti-CD3 FITC and DAPI was used as dead cell marker. No multiplet filtering was done to allow capture of large cTECs. First, CD45- cells (L) were sorted to obtain total stroma. Next, a gate including either CD205+ or EPCAM+ cells (K) was used to sort total thymic epithelial cells. **B.** Sorting scheme for CITE-seq experiments. Cells were stained with anti-CD3 PE and PI was used as dead cell marker. For each sample debris, doublets and PI+ cells were gated out and CD3+ and CD3- cells were sorted and collected separately. Plot is representative for all samples used in CITE-seq experiments.

**Supplementary Table 1. Single cell meta data.** Composition of all dissociated datasets used in this study. 'Age' denotes pcw for fetal samples and time after birth for infant/paediatric samples. 'age_numeric' was calculated as follows: for fetal samples: 'age_numeric' = (pcw - 40) and for paediatric samples: 'age_numeric' = (age in months). See table - 1_SuppTable_sc_meta.csv

**Supplementary Table 2. Spatial sample meta data.** Composition of all spatial data used in this study separated to: Visium, IBEX, RareCyte and RNAscope imaging. See table - 2_SuppTable_spatial_meta.xlsx

**Supplementary Table 3. IBEX antibody and imaging details.** Antibodies and imaging details for the custom IBEX panel implemented for all samples in this study:

| Cycle | Marker | Clone | Conjugate | Vendor | Cat. Number | Dilution | RRID |
|---|---|---|---|---|---|---|---|
| 1 | Hoechst | NA | NA | Biotium | 40046 | 1:5000 | NA |
| 1 | CD8 | SK1 | AF488 | BioLegend | 344716 | 1:25 | AB_10549301 |
| 1 | Chromogranin A | LK2H10 | AF532 | Novus Biologicals | NBP2-34672AF532 | 1:50 | AB_2892553 |
| 1 | CD163 | GHI/61 | PE | BioLegend | 333606 | 1:100 | AB_1134002 |
| 1 | CD3 | UCHT1 | iF594 | Caprico Biotechnologies | 1053136 | 1:300 | AB_2892737 |
| 1 | CD99 | 12E7 | NA | Abcam | Ab8855 | 1:200 | AB_306816 |
| 1 | Alpaca anti-Mouse | CTK0103/CTK0104 | AF647 | Thermo Fisher Scientific | SA5-10333 | 1:400 | AB_2868380 |
| 1 | CD11c | EP1347Y | NA | Abcam | Ab216655 | 1:100 | AB_2864379 |

| | | | | | | | |
|---|---|---|---|---|---|---|---|
| 1 | Donkey anti-Rabbit | Polyclonal | AF680 | Thermo Fisher Scientific | A10043 | 1:200 | AB_2534018 |
| 2 | Hoechst | NA | NA | Biotium | 40046 | 1:5000 | NA |
| 2 | CD20 | L26 | AF488 | Thermo Fisher Scientific | 53-0202-82 | 1:100 | AB_10734358 |
| 2 | CD7 | HIT7 | AF555 | AAT Bioquest | 10070161 | 1:200 | NA |
| 2 | CD34 | 4H11 | iF594 | AAT Bioquest | 103400C0 | 1:50 | AB_2892741 |
| 2 | CD5 | UCHT2 | AF647 | BioLegend | 300616 | 1:200 | AB_493037 |
| 2 | CD31 | WM59 | AF700 | BioLegend | 303133 | 1:10 | AB_2566326 |
| 3 | Hoechst | NA | NA | Biotium | 40046 | 1:5000 | NA |
| 3 | CD39 | A1 | FITC | BioLegend | 328206 | 1:10 | AB_940425 |
| 3 | β-tubulin III | TUJ1 | AF532 | BioLegend | NA, Custom | 1:100 | NA |
| 3 | Vα7.2 | 3C10 | PE | BioLegend | 351706 | 1:25 | AB_10899577 |
| 3 | CD4 | RPA-T4 | iF594 | AAT Bioquest | 100410C0 | 1:50 | NA |
| 3 | FOXP3 | 259D | AF647 | BioLegend | 320213 | 1:15 | AB_492985 |
| 3 | Donkey anti-Rat | Polyclonal | AF680 | Jackson ImmunoResearch | 712-625-150 | 1:200 | AB_2340698 |
| 4 | Hoechst | NA | NA | Biotium | 40046 | 1:5000 | NA |
| 4 | Lamin A | 133A2 | FITC | Thermo Fisher Scientific | MUB1101L1 | 1:10 | NA |
| 4 | CD123 | 6H6 | PE | BioLegend | 306005 | 1:25 | AB_314579 |
| 4 | HLA-DR | L243 | iF594 | Caprico Biotechnologies | 1032136 | 1:400 | AB_2892750 |
| 4 | CD206 | GH1/61 | AF647 | Thermo Fisher Scientific | MA5-44147 | 1:400 | AB_2913079 |
| 4 | SPARC | Polyclonal | NA | R&D Systems | AF941 | 1:200 | AB_2892754 |
| 4 | Donkey anti-Goat | Polyclonal | AF680 | Thermo Fisher Scientific | A21084 | 1:200 | AB_2535741 |
| 5 | Hoechst | NA | NA | Biotium | 40046 | 1:5000 | NA |
| 5 | Desmin | Y66 | AF488 | Abcam | ab185033 | 1:500 | AB_2892748 |
| 5 | DEC205 | JB87-35 | NA | Novus Biologicals | NBP2-75467 | 1:200 | NA |
| 5 | Zenon IgG | NA | AF532 | Thermo Fisher Scientific | Z25303 | NA | NA |
| 5 | AIRE | TM-724 | eF570 | Thermo Fisher Scientific | 41-9534-82 | 1:40 | AB_2573623 |
| 5 | Aquaporin 1 | EPR11588(B) | AF647 | Abcam | ab225225 | 1:3000 | NA |
| 5 | Ki-67 | B56 | AF700 | BD Biosciences | 561277 | 1:25 | AB_10611571 |
| 6 | Hoechst | NA | NA | Biotium | 40046 | 1:5000 | NA |
| 6 | CD49f | GoH3 | AF488 | BioLegend | 313607 | 1:25 | AB_493634 |
| 6 | LYVE-1 | Polyclonal | NA | R&D Systems | AF2089 | 1:250 | AB_355144 |
| 6 | Keratin 10 | DE-K10 | PE | Novus Biologicals | NBP2-54402PE | 1:500 | NA |
| 6 | Lumican | Polyclonal | iF594 | R&D Systems | AF2846 | 1:200 | AB_2139484 |
| 6 | CD49a | TS2/7 | AF647 | BioLegend | 328310 | 1:100 | AB_2129242 |
| 6 | Keratin 15 | Poly18339 | NA | BioLegend | 833904 | 1:600 | AB_2616894 |
| 6 | Goat anti-Chicken | Polyclonal | AF680 | Thermo Fisher Scientific | A32934 | 1:400 | AB_2762846 |
| 7 | Hoechst | NA | NA | Biotium | 40046 | 1:5000 | NA |
| 7 | Annexin 1 | REA112 | FITC | Miltenyi Biotec | 130-119-352 | 1:100 | AB_2857448 |

| | | | | | | | |
|---|---|---|---|---|---|---|---|
| 7 | Vimentin | 091D3 | AF532 | BioLegend | NA, Custom | 1:800 | AB_2892753 |
| 7 | Keratin 14 | Poly9060 | AF647 | BioLegend | NA, Custom | 1:1000 | NA |
| 8 | Hoechst | NA | NA | Biotium | 40046 | 1:5000 | NA |
| 8 | CD15 | MMA | AF532 | BioLegend | Custom | 1:400 | NA |
| 8 | γδ TCR | B1 | PE | BioLegend | 331209 | 1:50 | AB_1089219 |
| 8 | CD45 | F10-89-4 | iF594 | Caprico Biotechnologies | 1016136 | 1:300 | AB_2892743 |
| 8 | Synaptophysin D | SP17 | NA | BioLegend | 837103 | 1:200 | AB_2783410 |
| 8 | Donkey anti-Mouse | Polyclonal | AF647 | Jackson ImmunoResearch | 715-605-020 | 1:200 | AB_2340860 |
| 9 | Hoechst | NA | NA | Biotium | 40046 | 1:5000 | NA |
| 9 | Pan-Cytokeratin | AE1/AE3 | AF488 | Thermo Fisher Scientific | 53-9003-82 | 1:100 | AB_1834350 |
| 9 | Keratin 5 | Poly19055 | AF532 | BioLegend | NA, Custom | 1:300 | NA |
| 9 | α-SMA | 1A4 | eF570 | Thermo Fisher Scientific | 41-9760-82 | 1:200 | AB_2573631 |
| 9 | Keratin 8 | TS1 | AF647 | Novus Biologicals | NBP2-34267 | 1:200 | NA |
| 8 | γδ TCR | B1 | PE | BioLegend | 331209 | 1:50 | AB_1089219 |
| 8 | CD45 | F10-89-4 | iF594 | CapricoBio | 1016136 | 1:300 | AB_2892743 |
| 8 | Synaptophysin | SP17 | NA | BioLegend | 837103 | 1:200 | AB_2783410 |
| 8 | Donkey anti-Mouse | Polyclonal | AF647 | Jackson | 715-605-020 | 1:200 | AB_2340860 |
| 9 | Hoechst | NA | NA | Biotium | 40046 | 1:5000 | NA |
| 9 | Pan-Cytokeratin | AE1/AE3 | AF488 | Thermo | 53-9003-82 | 1:100 | AB_1834350 |
| 9 | Keratin 5 | Poly19055 | AF532 | BioLegend | NA, Custom | 1:300 | NA |
| 9 | α-SMA | 1A4 | eF570 | Thermo | 41-9760-82 | 1:200 | AB_2573631 |
| 9 | Keratin 8 | TS1 | AF647 | Novus | NBP2-34267 | 1:200 | NA |

**Supplementary Table 4. IBEX CITE-seq KNN matching details.** Matched IBEX protein to single cell RNA-seq data by RNA genes or CITEseq proteins (when available)

| Protein (IBEX) | Gene (scRNA-seq) | CITEseq |
|---|---|---|
| CHGA | CHGA | CHGA |
| CD99 | CD99 | cite_CD99 |
| CD163 | CD163 | cite_CD163 |
| CD11C | ITGAX | cite_CD11c |
| CD8 | CD8A | cite_CD8 |
| CD3 | CD3D | cite_CD3 |
| CD5 | CD5 | cite_CD5 |
| CD20 | MS4A1 | cite_CD20 |
| CD34 | CD34 | cite_CD34 |
| CD7 | CD7 | cite_CD7 |
| CD31 | PECAM1 | cite_CD31 |

| | | |
|---|---|---|
| CD39 | ENTPD1 | cite_CD39 |
| CD4 | CD4 | cite_CD4 |
| VA7.2 | TRAV7 | cite_TCR-Va7.2 |
| TUBB3 | TUBB3 | TUBB3 |
| CD206 | MRC1 | MRC1 |
| SPARC | SPARC | SPARC |
| LAMIN_A | LMNA | LMNA |
| HLADR | HLA-DRB1 | cite_HLA.DR |
| CD123 | IL3RA | cite_CD123 |
| DEC205 | LY75 | LY75 |
| AQP1 | AQP1 | AQP1 |
| KI67 | MKI67 | MKI67 |
| AIRE | AIRE | AIRE |
| CD49A | ITGA1 | cite_CD49a |
| KERATIN_10 | KRT10 | KRT10 |
| KERATIN_15 | KRT15 | KRT15 |
| LUMICAN | LUM | LUM |
| LYVE1 | LYVE1 | LYVE1 |
| VIMENTIN | VIM | VIM |
| KERATIN_14 | KRT14 | KRT14 |
| ANNEXIN1 | ANXA1 | ANXA1 |
| CD15 | FUT4 | FUT4 |
| CD45 | PTPRC | cite_CD45 |
| SYP | SYP | SYP |
| TCRGD | TCRG | cite_TCRgd |
| KERATIN_8 | KRT8 | KRT8 |
| PANCYTO | None | None |
| ASMA | ACTA2 | ACTA2 |
| KERATIN_5 | KRT5 | KRT5 |
| FOXP3 | FOXP3 | FOXP3 |
| DESMIN | DES | DES |
| CD49F | ITGA6 | cite_CD49f |

**Supplementary Table 5. RareCyte antibody panel.** RareCyte antibody composition details.

| Antibody | Flourophore (nm) | Details | Product code |
|---|---|---|---|
| CD31 | 515 | Anti-Human CD31 (EPR3094)-ArgoFluor™ 515 | 52-1005-501 |
| CD68 | 535 | Anti-Human CD68 (D4B9C)-ArgoFluor™ 535 | 52-1008-501 |
| KI67 | 555 | Anti-Human Ki-67 (D3B5)-ArgoFluor™ 555L | 52-1013-501 |
| CD4 | 572 | Anti-Human CD4 (N1UG0)-ArgoFluor™ 572 | 52-1002-501 |
| CD163 | 580 | Anti-Human CD163 (EPR14643)-ArgoFluor™ 580L | 52-1009-501 |
| CD8a | 602 | Anti-Human CD8a (AMC908)-ArgoFluor™602 | 52-1003-601 |
| CD20 | 660 | Anti-Human CD20 (L26)-ArgoFluor™ 660L | 52-1004-601 |
| FOXP3 | 662 | Anti-Human FOXP3 (236A/E7)-ArgoFluor™ 662 | 52-1012-601 |
| CD3e | 686 | Anti-Human CD3e (D7A6E)-ArgoFluor™ 686 | 52-1001-601 |
| CD45 | 810 | Anti-Human CD45 (D9M8I)-ArgoFluor™ 810 | 52-1006-801 |
| PanCK | 845 | Anti-Human Pan-CK (C-11, AE1/AE3)-ArgoFluor™ 845 | 52-1015-801 |
| VIM | 874 | Anti-Human Vimentin (O91D3)-ArgoFluor™ 874 | 52-1019-801 |

**Supplementary Table 6. RNAscope probes, conditions and controls.** See table - 6_SuppTable_RNAscope_procedures.xlsx

**Supplementary Table 7. CITE-seq antibody panel.** Information about Biolegend TotalSeq-C antibodies used in this study.

| Cat.no. | Barcode | Target | Clone | TotalSeq-C antibody |
|---|---|---|---|---|
| 305447 | C0006 | CD86 | IT2.2 | Human Universal Cocktail v1 |
| 329751 | C0007 | CD274 | 29E.2A3 | Human Universal Cocktail v1 |
| 318815 | C0020 | CD270 | 122 | Human Universal Cocktail v1 |
| 337635 | C0023 | CD155 | SKII.4 | Human Universal Cocktail v1 |
| 337419 | C0024 | CD112 | TX31 | Human Universal Cocktail v1 |
| 323131 | C0026 | CD47 | CC2C6 | Human Universal Cocktail v1 |
| 336711 | C0029 | CD48 | BJ40 | Human Universal Cocktail v1 |
| 334348 | C0031 | CD40 | 5C3 | Human Universal Cocktail v1 |
| 310849 | C0032 | CD154 | 24-31 | Human Universal Cocktail v1 |
| 316021 | C0033 | CD52 | HI186 | Human Universal Cocktail v1 |
| 300479 | C0034 | CD3 | UCHT1 | Human Universal Cocktail v1 |
| 344753 | C0046 | CD8 | SK1 | Human Universal Cocktail v1 |
| 362559 | C0047 | CD56 | 5.1H11 | Human Universal Cocktail v1 |
| 302265 | C0050 | CD19 | HIB19 | Human Universal Cocktail v1 |
| 366633 | C0052 | CD33 | P67.6 | Human Universal Cocktail v1 |
| 371521 | C0053 | CD11c | S-HCL-3 | Human Universal Cocktail v1 |
| 343541 | C0054 | CD34 | 581 | Individual TotalSeq-C antibody |
| 311449 | C0058 | HLA-A,B,C | W6/32 | Human Universal Cocktail v1 |
| 312233 | C0062 | CD10 | HI10a | Individual TotalSeq-C antibody |

| | | | | |
|---|---|---|---|---|
| 304163 | C0063 | CD45RA | HI100 | Human Universal Cocktail v1 |
| 306045 | C0064 | CD123 | 6H6 | Human Universal Cocktail v1 |
| 343127 | C0066 | CD7 | CD7-6B7 | Human Universal Cocktail v1 |
| 323227 | C0068 | CD105 | 43A3 | Human Universal Cocktail v1 |
| 313635 | C0070 | CD49f | GoH3 | Human Universal Cocktail v1 |
| 359425 | C0071 | CD194 | L291H4 | Human Universal Cocktail v1 |
| 300567 | C0072 | CD4 | RPA-T4 | Human Universal Cocktail v1 |
| 103063 | C0073 | CD44 | IM7 | Human Universal Cocktail v1 |
| 301859 | C0081 | CD14 | M5E2 | Human Universal Cocktail v1 |
| 302065 | C0083 | CD16 | 3G8 | Human Universal Cocktail v1 |
| 302649 | C0085 | CD25 | BC96 | Human Universal Cocktail v1 |
| 304259 | C0087 | CD45RO | UCHL1 | Human Universal Cocktail v1 |
| 329963 | C0088 | CD279 | EH12.2H7 | Human Universal Cocktail v1 |
| 372729 | C0089 | TIGIT | A15153G | Human Universal Cocktail v1 |
| 400187 | C0090 | Mouse IgG1, κ isotype Ctrl | MOPC-21 | Human Universal Cocktail v1 |
| 400293 | C0091 | Mouse IgG2a, κ isotype Ctrl | MOPC-173 | Human Universal Cocktail v1 |
| 400381 | C0092 | Mouse IgG2b, κ isotype Ctrl | MPC-11 | Human Universal Cocktail v1 |
| 400677 | C0095 | Rat IgG2b, κ Isotype Ctrl | RTK4530 | Human Universal Cocktail v1 |
| 302363 | C0100 | CD20 | 2H7 | Human Universal Cocktail v1 |
| 331941 | C0101 | CD335 | 900 | Human Universal Cocktail v1 |
| 303139 | C0124 | CD31 | WM59 | Human Universal Cocktail v1 |
| 361033 | C0134 | CD146 | P1H12 | Human Universal Cocktail v1 |
| 314547 | C0136 | IgM | MHM-88 | Human Universal Cocktail v1 |
| 300637 | C0138 | CD5 | UCHT2 | Human Universal Cocktail v1 |
| 331231 | C0139 | TCRgd | B1 | Individual TotalSeq-C antibody |
| 353747 | C0140 | CD183 | G025H7 | Human Universal Cocktail v1 |
| 359137 | C0141 | CD195 | J418F1 | Human Universal Cocktail v1 |
| 303225 | C0142 | CD32 | FUN-2 | Human Universal Cocktail v1 |
| 353440 | C0143 | CD196 | G034E3 | Human Universal Cocktail v1 |
| 356939 | C0144 | CD185 | J252D4 | Human Universal Cocktail v1 |
| 350233 | C0145 | CD103 | Ber-ACT8 | Human Universal Cocktail v1 |
| 310951 | C0146 | CD69 | FN50 | Human Universal Cocktail v1 |
| 304851 | C0147 | CD62L | DREG-56 | Human Universal Cocktail v1 |
| 353251 | C0148 | CD197 | G043H7 | Individual TotalSeq-C antibody |
| 339947 | C0149 | CD161 | HP-3G10 | Human Universal Cocktail v1 |
| 369621 | C0151 | CD152 | BNI3 | Human Universal Cocktail v1 |
| 369335 | C0152 | CD223 | 11C3C65 | Human Universal Cocktail v1 |
| 367737 | C0153 | KLRG1 | SA231A2 | Human Universal Cocktail v1 |
| 302853 | C0154 | CD27 | O323 | Human Universal Cocktail v1 |
| 328649 | C0155 | CD107a | H4A3 | Human Universal Cocktail v1 |
| 305651 | C0156 | CD95 | DX2 | Human Universal Cocktail v1 |
| 350035 | C0158 | CD134 | Ber-ACT35 (ACT35) | Human Universal Cocktail v1 |
| 307663 | C0159 | HLA-DR | L243 | Human Universal Cocktail v1 |
| 331547 | C0160 | CD1c | L161 | Human Universal Cocktail v1 |
| 301359 | C0161 | CD11b | ICRF44 | Human Universal Cocktail v1 |

| | | | | |
|---|---|---|---|---|
| 305045 | C0162 | CD64 | 10.1 | Human Universal Cocktail v1 |
| 344125 | C0163 | CD141 | M80 | Human Universal Cocktail v1 |
| 350319 | C0164 | CD1d | 51.1 | Human Universal Cocktail v1 |
| 320837 | C0165 | CD314 | 1D11 | Human Universal Cocktail v1 |
| 333409 | C0167 | CD35 | E11 | Human Universal Cocktail v1 |
| 393321 | C0168 | CD57 | QA17A04 | Human Universal Cocktail v1 |
| 344527 | C0170 | CD272 | MIH26 | Human Universal Cocktail v1 |
| 313553 | C0171 | CD278 | C398.4A | Human Universal Cocktail v1 |
| 330921 | C0174 | CD58 | TS2/9 | Human Universal Cocktail v1 |
| 328237 | C0176 | CD39 | A1 | Human Universal Cocktail v1 |
| 355705 | C0179 | CX3CR1 | K0124E1 | Human Universal Cocktail v1 |
| 311143 | C0180 | CD24 | ML5 | Human Universal Cocktail v1 |
| 354923 | C0181 | CD21 | Bu32 | Human Universal Cocktail v1 |
| 350617 | C0185 | CD11a | TS2/4 | Human Universal Cocktail v1 |
| 341417 | C0187 | CD79b | CB3-1 | Human Universal Cocktail v1 |
| 329529 | C0189 | CD244 | C1.7 | Human Universal Cocktail v1 |
| 346021 | C0206 | CD169 | 7-239 | Human Universal Cocktail v1 |
| 353809 | C0207 | CD370 | 8F9 | Individual TotalSeq-C antibody |
| 372617 | C0208 | XCR1 | S15046E | Individual TotalSeq-C antibody |
| 352115 | C0213 | Notch 1 | MHN1-519 | Individual TotalSeq-C antibody |
| 321229 | C0214 | Integrin b7 | FIB504 | Human Universal Cocktail v1 |
| 316927 | C0215 | CD268 | 11C1 | Human Universal Cocktail v1 |
| 303941 | C0216 | CD42b | HIP1 | Human Universal Cocktail v1 |
| 353127 | C0217 | CD54 | HA58 | Human Universal Cocktail v1 |
| 304945 | C0218 | CD62P | AK4 | Human Universal Cocktail v1 |
| 308613 | C0219 | CD119 | GIR-208 | Human Universal Cocktail v1 |
| 306743 | C0224 | TCRab | IP26 | Human Universal Cocktail v1 |
| 345413 | C0233 | Notch3 | MHN3-21 | Individual TotalSeq-C antibody |
| 400467 | C0236 | Rat IgG1, kappa Isotype Ctrl | RTK2071 | Human Universal Cocktail v1 |
| 400577 | C0238 | Rat IgG2a, kappa Isotype Ctrl | RTK2758 | Human Universal Cocktail v1 |
| 400977 | C0241 | Armenian Hamster IgG Isotype Ctrl | HTK888 | Human Universal Cocktail v1 |
| 339021 | C0246 | CD122 | TU27 | Human Universal Cocktail v1 |
| 311915 | C0247 | CD267 | 1A1 | Human Universal Cocktail v1 |
| 334645 | C0352 | FceRIa | AER-37 (CRA-1) | Human Universal Cocktail v1 |
| 303739 | C0353 | CD41 | HIP8 | Human Universal Cocktail v1 |
| 309839 | C0355 | CD137 | 4B4-1 | Human Universal Cocktail v1 |
| 333637 | C0358 | CD163 | GHI/61 | Human Universal Cocktail v1 |
| 305341 | C0359 | CD83 | HB15e | Human Universal Cocktail v1 |
| 371227 | C0360 | CD357 | 108-17 | Individual TotalSeq-C antibody |
| 355011 | C0363 | CD124 | G077F6 | Human Universal Cocktail v1 |
| 301733 | C0364 | CD13 | WM15 | Human Universal Cocktail v1 |
| 309231 | C0367 | CD2 | TS1/8 | Human Universal Cocktail v1 |
| 338337 | C0368 | CD226 | 11A8 | Human Universal Cocktail v1 |
| 303029 | C0369 | CD29 | TS2/16 | Human Universal Cocktail v1 |
| 354241 | C0370 | CD303 | 201A | Human Universal Cocktail v1 |

| | | | | |
|---|---|---|---|---|
| 359317 | C0371 | CD49b | P1E6-C5 | Human Universal Cocktail v1 |
| 349523 | C0373 | CD81 | 5A6 | Human Universal Cocktail v1 |
| 348245 | C0384 | IgD | IA6-2 | Human Universal Cocktail v1 |
| 302129 | C0385 | CD18 | TS1/18 | Human Universal Cocktail v1 |
| 302963 | C0386 | CD28 | CD28.2 | Human Universal Cocktail v1 |
| 303543 | C0389 | CD38 | HIT2 | Human Universal Cocktail v1 |
| 351356 | C0390 | CD127 | A019D5 | Human Universal Cocktail v1 |
| 304068 | C0391 | CD45 | HI30 | Human Universal Cocktail v1 |
| 363516 | C0393 | CD22 | S-HCL-1 | Human Universal Cocktail v1 |
| 334125 | C0394 | CD71 | CY1G4 | Human Universal Cocktail v1 |
| 302722 | C0396 | CD26 | BA5b | Human Universal Cocktail v1 |
| 300135 | C0402 | CD1a | HI149 | Individual TotalSeq-C antibody |
| 336227 | C0407 | CD36 | 5-271 | Human Universal Cocktail v1 |
| 372111 | C0408 | CD172a | 15-414 | Individual TotalSeq-C antibody |
| 339517 | C0420 | CD158 | HP-MA4 | Human Universal Cocktail v1 |
| 328319 | C0575 | CD49a | TS2/7 | Human Universal Cocktail v1 |
| 304345 | C0576 | CD49d | 9F10 | Human Universal Cocktail v1 |
| 344031 | C0577 | CD73 | AD2 | Human Universal Cocktail v1 |
| 351735 | C0581 | TCR-Va7.2 | 3C10 | Human Universal Cocktail v1 |
| 331435 | C0582 | TCR-Vd2 | B6 | Human Universal Cocktail v1 |
| 342925 | C0584 | TCR-Va24-Ja18 | 6B11 | Individual TotalSeq-C antibody |
| 358613 | C0591 | LOX-1 | 15C4 | Human Universal Cocktail v1 |
| 312619 | C0592 | CD158b/j | DX27 | Human Universal Cocktail v1 |
| 312725 | C0599 | CD158e1 | DX9 | Human Universal Cocktail v1 |
| 337113 | C0805 | CD226 | TX25 | Human Universal Cocktail v1 |
| 331823 | C0830 | CD319 | 162.1 | Human Universal Cocktail v1 |
| 358921 | C0843 | CD199 | L053E8 | Individual TotalSeq-C antibody |
| 371321 | C0845 | CD99 | 3B2/TA8 | Human Universal Cocktail v1 |
| 353615 | C0853 | CD371 | 50C1 | Human Universal Cocktail v1 |
| 317213 | C0864 | CD352 | NT-7 | Human Universal Cocktail v1 |
| 305523 | C0867 | CD94 | DX22 | Human Universal Cocktail v1 |
| 316533 | C0894 | Ig light chain kappa | MHK-49 | Human Universal Cocktail v1 |
| 333725 | C0896 | CD85j | GHI/75 | Human Universal Cocktail v1 |
| 338525 | C0897 | CD23 | EBVCS-5 | Human Universal Cocktail v1 |
| 316629 | C0898 | Ig light chain lambda | MHL-38 | Human Universal Cocktail v1 |
| 339213 | C0902 | CD328 | 6-434 | Human Universal Cocktail v1 |
| 358209 | C0912 | GPR56 | CG4 | Human Universal Cocktail v1 |
| 342619 | C0918 | HLA-E | 3D12 | Human Universal Cocktail v1 |
| 342115 | C0920 | CD82 | ASL-24 | Human Universal Cocktail v1 |
| 331017 | C0944 | CD101 | BB27 | Human Universal Cocktail v1 |
| 344319 | C1046 | CD88 | S5/1 | Human Universal Cocktail v1 |
| 394307 | C1052 | CD224 | KF29 | Human Universal Cocktail v1 |

**Supplementary Table 8. Cell Cycle Genes.** Genes excluded prior to scVI integration. See 8_SuppTable_8_cellcyclegenes.csv
